# Deep-tissue spatial omics: imaging whole-embryo transcriptomics and subcellular structures at high spatial resolution

**DOI:** 10.1101/2024.05.17.594641

**Authors:** Valentina Gandin, Jun Kim, Liang-Zhong Yang, Yumin Lian, Takashi Kawase, Amy Hu, Konrad Rokicki, Greg Fleishman, Paul Tillberg, Alejandro Aguilera Castrejon, Carsen Stringer, Stephan Preibisch, Zhe J. Liu

**Author notes:** Equal contribution (Valentina Gandin, Jun Kim). Correspondence (Z.J.L.).

## Abstract

The inherent limitations of fluorescence microscopy, notably the restricted number of color channels, have long constrained comprehensive spatial analysis in biological specimens. Here, we introduce cycleHCR technology that leverages multicycle DNA barcoding and Hybridization Chain Reaction (HCR) to surpass the conventional color barrier. cycleHCR facilitates high-specificity, single-shot imaging per target for RNA and protein species within thick specimens, mitigating the molecular crowding issues encountered with other imaging-based spatial omics techniques. We demonstrate whole-mount transcriptomics imaging of 254 genes within an E6.5∼7.0 mouse embryo, achieving precise three-dimensional gene expression and cell fate mapping across a specimen depth of ∼ 310 µm. Utilizing expansion microscopy alongside protein cycleHCR, we unveil the complex network of 10 subcellular structures in primary mouse embryonic fibroblasts. Furthermore, in mouse hippocampal slice, we image 8 protein targets and profile the transcriptome of 120 genes, uncovering complex gene expression gradients and cell-type specific nuclear structural variances. cycleHCR provides a unifying framework for multiplex RNA and protein imaging, offering a quantitative solution for elucidating spatial regulations in deep tissue contexts for research and potentially diagnostic applications.

## Main text

Understanding the spatial organization of molecular components within complex tissue samples is crucial for deciphering the biological regulations that underpin animal development and disease states. Significant advancements in microscopy, tissue clearing and expansion techniques over the past decades have enabled deeper imaging with enhanced clarity and resolution (*1–3*). However, a fundamental limitation of fluorescent imaging remains - it cannot simultaneously image multiple molecular components or color channels due to the restricted availability of labeling dyes and their spectrum overlaps.

Recent advances in high-throughput spatial omics, notably single-molecule *in situ* hybridization techniques like MERFISH and seqFISH (*4, 5*), as well as *in situ* sequencing methods such as FISSEQ and STARmap (*6, 7*), have markedly improved our capacity to spatially decode molecular identities, particularly for RNA species. Despite these innovations, these approaches encounter significant challenges. One such challenge is molecular crowding, which can restrict the decoding capacity for targets that are either abundant or exhibit heterogeneous distributions. Additionally, the necessity to detect dim single molecules with high numerical aperture objectives severely limits axial imaging depth in *in situ* hybridization-based techniques (*8*). Rolling circle amplification (RCA), when combined with expansion microscopy, offers potential solutions to circumvent molecular crowding and enhance imaging depth (*9*). However, the inherently low detection efficiency of RCA limits sensitivity and coverage for target detection. Moreover, these methods require precise cross-round registration with nanometer resolution for accurate assignment of detected spots to specific genes, a task that becomes progressively more difficult with increasing specimen size. The lack of empirical ground truth images for verification further complicates the interpretation of RNA target distributions generated by these methods, which rely on cross-round barcoding at the single-molecule level (*8*).

The development of split Hybridization Chain Reaction (HCR) techniques has begun to address several critical challenges in deep tissue imaging: First, the proximity binding requirement of split probes to initiate HCR allows for high specificity in the probe hybridization process. This feature enables single-shot imaging of RNA and protein molecules without the need for cross-round decoding and proofing (*10–12*). Second, HCR based signal amplification permits the use of objectives with low numerical apertures and long working distances, enabling reliable deep-tissue detection. Third, the inherent single-shot per target nature of HCR imaging obviates the need for cross-round decoding, making densely labeled targets. Achieving multi-round HCR imaging has been demonstrated through the time-consuming process of primary probe removal by DNase treatment and rehybridization (*13*). Currently, this method allows for only one round of imaging every 3 – 5 days, creating a bottleneck in the workflow and limiting the number of targets that can be effectively examined.

### cycleHCR: concept and implementation

Here, we introduce cycleHCR technology that enables high-throughput, single-shot imaging of RNA and protein species within thick tissue specimens. At the core of cycleHCR RNA imaging lies the optimization and selection of 45bp split primary probes (Fig. 1A and fig. S1) Distinguished by their high melting temperatures (>80°C) for probe-RNA interactions, these probes ensure robust and stable binding under stringent conditions efficiently clearing away other components such as HCR chains and barcoding probes (fig. S1-S3). Another distinction of cycleHCR is the introduction of a barcoding phase utilizing pairs of 14bp Left (L) and Right (R) DNA barcoding probes equipped with split HCR initiators (Fig. 1A). These probes are designed to trigger HCR reactions only when perfectly matched to the barcode on the target (Fig. 1B), achieving high-specificity target recognition both in cultured cells and thick tissue specimens (Fig. 1B, 1C and 1E). Through the combination of 30 unique L and 30 unique R probes per color channel, cycleHCR facilitates the generation of up to 900 distinct barcodes, thereby enabling the potential encoding of 2,700 targets across three channels (Fig. 1D).

**Fig. 1.**
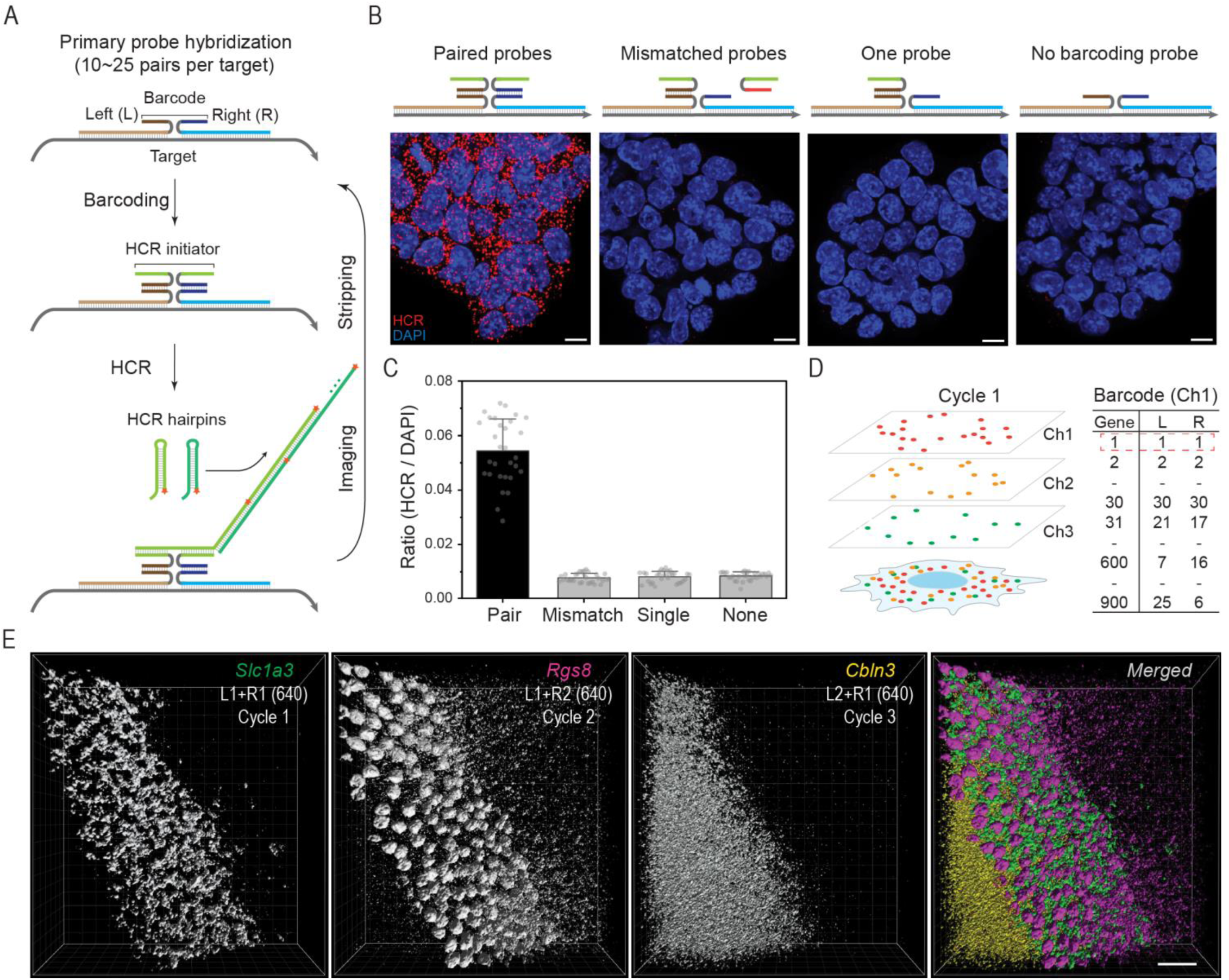
cycleHCR RNA imaging. **(A)** To target each RNA sequence with high specificity, we engineered DNA probe libraries with 10 to 25 pairs of 45-bp probes. Each pair consists of left and right probes with unique 14-bp readout sequences for binding barcoded left (L) and right (R) readout probes with split HCR initiators, triggering the hybridization chain reaction (HCR). **(B)** The HCR amplification occurs only with perfectly matched L and R readout probe pairs. Mismatched probes, single probes, or absence of probes prevent HCR initiation, minimizing false positives. Target: *Trim6*; L2+R2 (561). Scale bars: 10µm. **(C)** Quantification of images shown in (B) is performed by comparing the HCR signal to the nuclear DAPI signal. The error bars represent standard deviations. **(D)** The mixing of 30 left and 30 right probes, each with unique divergent sequences, allows for the creation of up to 900 distinct barcodes for each fluorescence channel. Images are acquired using three separate fluorescence channels (488nm, 561nm, 640nm), each channel harboring orthogonal B4, B2, and B3 HCR initiators. **(E)** This panel demonstrates cycleHCR labeling specificity in thick specimens for three distinct cell-type marker genes. Through sequential three-round cycleHCR imaging, we show specific labeling changes for *Slc1a3* (L1+R1 at 640nm, marking Bergmann glia cells), *Rgs8* (L1+R2 at 640nm, marking purkinje cells), and *Cbln3* (L2+R1 at 640nm, marking the granule layer) in cerebellum specimens approximately 200µm thick. 3D images are rendered using the normal shading mode in Imaris. Scale bar: 50µm.

**Fig. 2.**
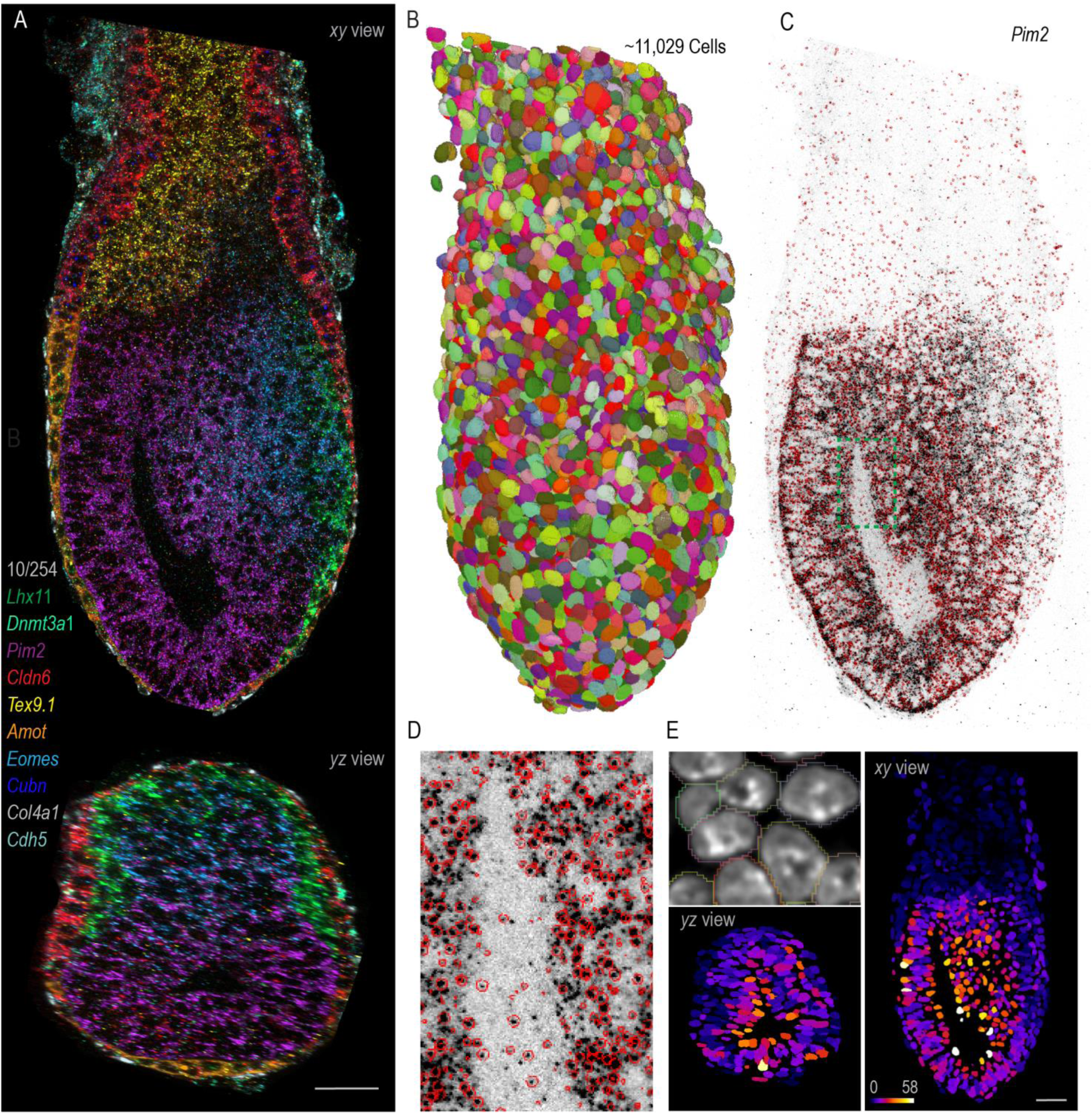
Whole-embryo transcriptomics imaging across a depth of ∼310µm. **(A)** Orthogonal *xy* and *yz* views of a 10-color composite image displaying the expression patterns of 10 genes (*Lhx1*, *Dhmt3a1*, *Pim2*, *Cldn6*, *Tex9.1*, *Amot*, *Eomes*, *Cubn*, *Col4a1*, and *Cdh5*) out of 254 genes imaged by cycleHCR (85 cycles) in an E6.5∼7.0 mouse embryo. The slice views were rendered by the maximal intensity projection (MIP) view using Imaris software. **(B)** Random colored masks for 11,029 cells segmented by Cellpose and rendered by ORS Dragonfly software. **(C)** An inverted raw image showing the detection of *Pim2* mRNA transcripts at the single-molecule level (black spots), with single-molecule localizations encircled in red. **(D)** A zoomed-in view of the region marked in (C), providing enhanced detail on the accuracy of single-molecule localization. **(E)** 3D spatial gene expression maps for *Pim2* mRNA, with spots assigned to individual cells based on Cellpose masks that are slightly dilated compared to DAPI labeled nucleus. Cells in the resulting *xy* and *yz* views are color-coded based on transcript counts indicated by the provided color map. Scale bars: 50µm.

**Fig. 3.**
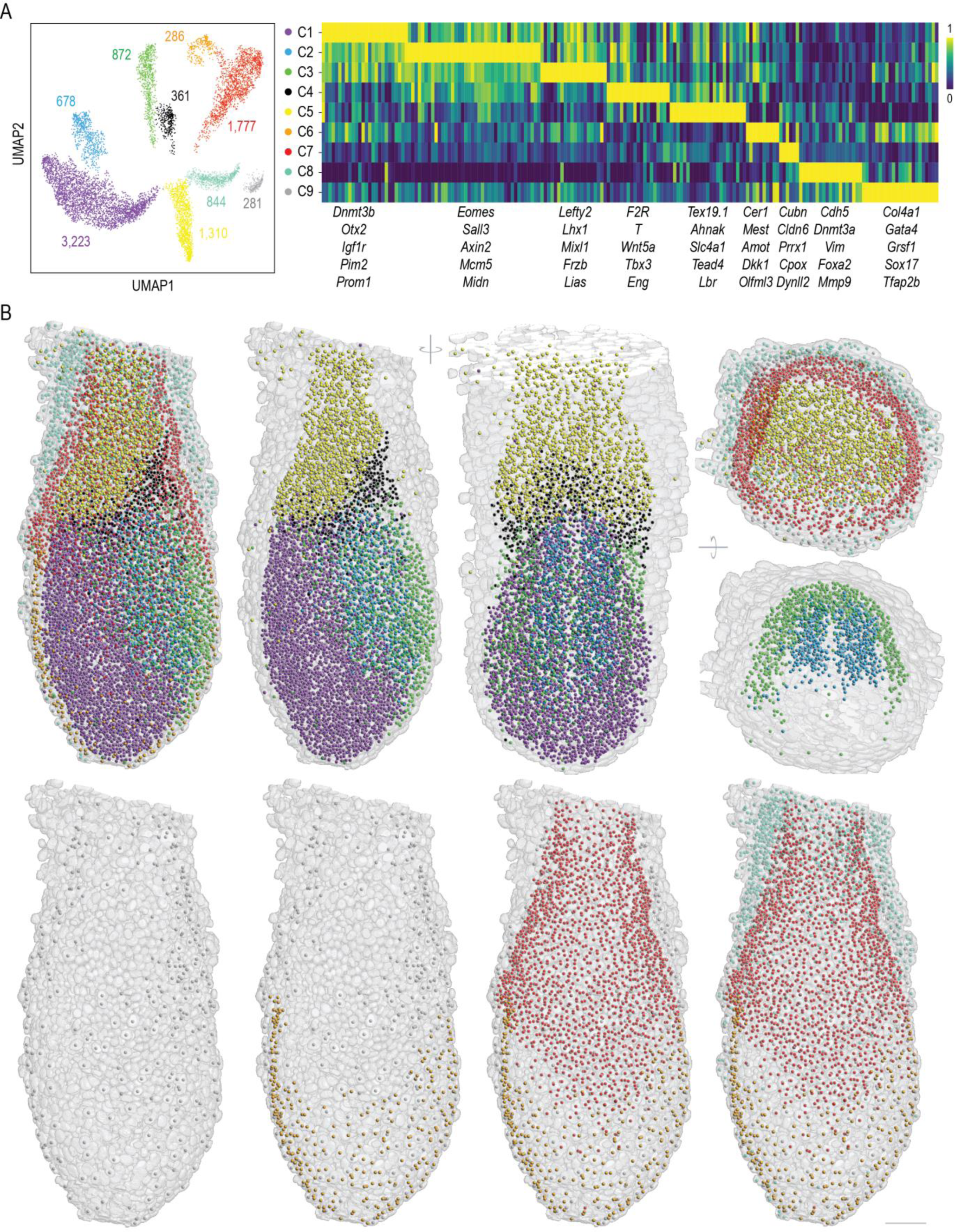
3D cell-map reconstruction. **(A)** UMAP analysis and gene clustering on single-cell transcript counts for 186 expressed genes detected by cycleHCR identify 9 distinct cell clusters. The number of cells within each cluster is indicated in the left panel. The right panel shows the clustering of the 186 genes, with 5 representative genes for individual clusters highlighted below. The color map represents the min-max scaled expression level for each gene across clusters. **(B)** A 3D cell-map, color-coded as per (A), displays the center of mass for each cell with a surrounding mesh for nucleus outlines. The left panel shows all 9 clusters. Upper middle panels provide views of clusters C1-5 from the same and a ∼90-degree rotated orientation, and the right panel displays the extra-embryonic layers (C5, C7, C8, and C9) alongside two closely situated intra-embryo clusters (C2 and C3). Lower panels detail the spatial distribution of outer layer clusters (C6-C9). Rendered using the napari Python package. Scale bar: 50µm.

The use of highly stable primary probes and precise barcoding negates the need for primary probe removal and rehybridization during multicycle HCR imaging. Through optimization, a consistent temperature of 32°C has been determined to allow for signal removal in 20 minutes (fig. S3) and HCR amplification in 1.5 hours (fig. S4), achieving about 51% of the signal intensity of overnight amplification. Efficient barcoding is accomplished in under 30 minutes using high concentrations (150 ∼ 200nM) of L + R probes at this temperature. These optimizations collectively reduce the duration of one detection cycle to within 4 hours. Additionally, employing an oxygen scavenger imaging buffer prevents photo-crosslinking during imaging (fig. S5), maintaining detection fidelity across multiple imaging cycles.

**Fig. 4.**
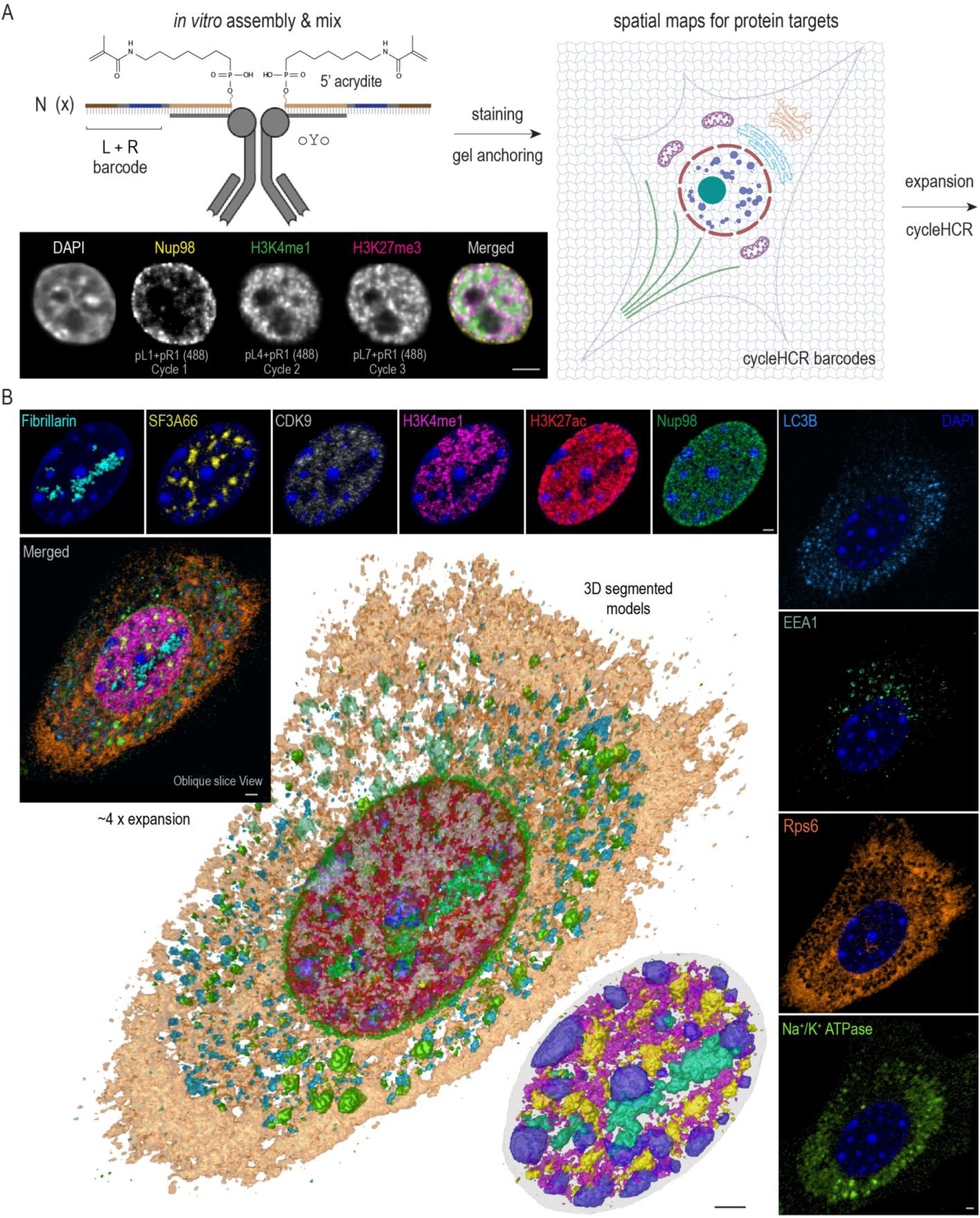
cycleHCR protein imaging with expansion microscopy. **(A)** cycleHCR protein labeling involves an *in vitro* tri-functional antibody complex assembly. This method covalently attaches a unique sequence to the antibody using the oYo linker. The sequence is then hybridized with a gel anchoring probe that contains a cycleHCR readout barcode and a 5’ acrydite group for attachment to a polyacrylamide gel. Following the *in vitro* assembly, the sample is incubated with the antibody mix. After gel embedding, the cycleHCR barcodes are covalently incorporated into the gel matrix, allowing for subsequent tissue expansion and protein mapping. The specificity of this protein labeling strategy is demonstrated through sequential three-cycle imaging by merely adjusting the right readout probes with the left readout probe fixed. The labeling shifts from the nuclear pore marker Nup98 (pL1+pR1 at 488nm) to the euchromatin marker H3K4me1 (pL4+pR1 at 488nm), and eventually to the heterochromatin marker H3K27me3 (pL7+pR1 at 488nm). **(B)** The integration of protein cycleHCR with expansion microscopy, illustrated by 3D surpass and oblique slice composite views (rendered by Imaris) of a primary mouse embryonic fibroblast after approximately 4X expansion. cycleHCR targets 10 proteins, including Fibrillarin (nucleolar fibrillar regions), SF3A66 (nuclear speckles), CDK9 (RNA Polymerase II elongation), H3K27ac (active chromatin), H3K4me1 (enhancer regions), Nup98 (nuclear pores), LC3B (autophagosomes), EEA1 (early endosomes), Rps6 (ribosome), and Na^+^/K^+^ ATPase. Central to the panel is the 3D segmented models rendered by ORS Dragonfly software with a detailed zoom-in on the nucleus (bottom left) showing the spatial relationships among heterochromatin, Fibrillarin, H3K4me1, and SF3A66. Scale bars: 5µm.

Transitioning towards full automation, our setup incorporates a programmable robotic arm for precise preparation of L + R readout mixes (fig. S2C), and an automated imaging and fluidic system capable of executing cycleHCR protocols and image acquisition without the need of human supervision (fig. S2B and S2D). This setup supports up to five imaging rounds analyzing 15 RNA species daily. Consistency and reproducibility in data analysis are facilitated by a scalable and portable Nextflow (*14*) image processing workflow that manages essential tasks such as image stitching, cross-cycle registration, single-molecule localization, 3D cell segmentation, and spot-to-cell assignment (fig. S6 and S7).

For high-resolution imaging of thick specimens, our setup utilizes spinning disk confocal microscopy equipped with silicone oil immersion objectives that offer long working distances up to 550 µm. Integrating hydrogel-embedding and enzymatic digestion-based tissue clearing with cycleHCR allows for detailed 3D visualization of complex sub-cellular RNA patterns within thick tissues (fig. S8A; Movie 1). For instance, within a ∼200µm section of mouse cerebellum, we observed dense *Slc1a3* mRNA clusters in Bergmann glia (fig. S8B), *Rgs8* gene transcriptional bursting sites in Purkinje nuclei (fig. S8C), and *Rgs8* transcripts along dendrites (fig. S8D). Interestingly, about half of Purkinje cells showed two *Rgs8* bursting sites, with no cell exceeding this number (fig. S8E). This observation hints at a diploid genome within Purkinje cells, offering insights into the discussion regarding their ploidy and genomic constitution (*15*).

### Whole-embryo transcriptomics imaging

To explore gene regulation and cell-fate determination in early development, we focused on an intact E6.5-7.0 mouse embryo, requiring ∼310 µm axial coverage. Leveraging cycleHCR, we performed whole-mount transcriptomics imaging of 254 lineage-specific genes through 85 imaging cycles over 20 days, achieving edge-to-edge imaging clarity and single-molecule detection sensitivity (Fig. 2A-D; Movie 2; Table S1). The synergy between high system stability and robust image registration enables precise 3D alignment of images throughout imaging cycles (fig. S9), ensuring the reliable spatial decoding of RNA distributions. The resulting single-shot images revealed distinct mRNA localization patterns (fig. S10-S16), highlighting the complex genetic interplay underlying early embryogenesis.

Our automated system standardizes cycleHCR reaction and imaging conditions, ensuring consistent RNA molecule detection across cycles and color channels using RS-FISH (*16*) (Fig. 2D and fig. S17). For single-cell gene expression analysis, we utilized Cellpose (*17*) for 3D nucleus segmentation, applying a specialist model trained on both *xy* and *yz* planes and refined through size filtering, reaching ∼87.6% initial accuracy. Manual curation further refined segmentation, identifying ∼11,029 nuclei (fig. S18). After evaluating various spot-to-cell assignment methods, we adopted the most conservative approach, assigning spots based on Cellpose masks which are slightly dilated compared to nuclei (Fig. 2E and fig. S19). This strategy only captures spots in proximity to or within the nucleus and minimizes contamination for accurate 3D gene expression mapping at the single-cell level (Table S2), paving the way for in-depth analysis.

### 3D cell-fate mapping

After initial filtering, we identified 186 actively transcribed genes and excluded 484 cells with low gene expression levels (fig. S20). Uniform Manifold Approximation and Projection (UMAP) (*18*) analysis on these genes revealed 9 distinct cell clusters (C1-C9) (Fig. 3A), expanding from ∼6 clusters previously recognized through single-cell sequencing at this developmental stage (*19–21*). These clusters span a wide range of cell counts, from 286 in the smallest cluster (C6) to 3,223 in the largest (C1), leaving only 876 cells unallocated. Gene clustering, UMAP imputation, and 3D spatial gene expression visualization showed extensive gene expression overlaps among clusters (Fig 3A; fig. S21-S29), indicating a complex genetic basis for cell-fate specification. For instance, the well-known primitive streak marker gene *T* was found to be over-represented in clusters C2, C3, and C4, with the highest enrichment observed in C4 (fig. S24).

Preliminary lineage assignment, based on marker gene analysis with comparison to previous studies (*22, 23*), suggests C1 as the epiblast (*Pim2*, *Dnmt3b* and *Prom1*), C2 as the primitive streak (*Eomes, Axin2 and Mcm5*), C3 as nascent mesoderm (*Lefty2*, *Mixl1* and *Frzb*), C4 as a cluster of mix lineages (e.g. *T* (primitive streak and allantois progenitors (*24*)), *Tbx3* (allantois progenitors) *and F2r* (blood progenitors)), C5 as extraembryonic ectoderm (*Tex19.1, Ahnak and Tead4*), C6 as anterior visceral endoderm (*Cer1* and *Dkk1*), C7, C9 as diverse endoderm lineages (*Col4a1*, *Sox17*, *Cubn*, *Gata4* and *Cldn6*) and C8 as putative endothelial cells (*Cdh5* and *Vim*).

To gain insights into the spatial organization and developmental context of these transcriptionally defined clusters, we proceeded to map them back to their 3D spatial locations within the embryo, revealing well-organized structures (Fig. 3B; Movie 3). Specifically, the 3D map shows the anatomical segregation of interior clusters - C1 (epiblast), C2 (primitive streak), C3 (nascent mesoderm), C4, and C5 (extraembryonic ectoderm) - into embryonic (C1, C2, C3) and extraembryonic (C5) domains, with C4 serving as a demarcation between the two domains. Within the embryo, the epiblast (C1), primitive streak (C2), and nascent mesoderm (C3), arrange into distinct, symmetrical layers across the midline. The endoderm clusters, C6, C7 and C9, also demonstrate spatial segregations; with C6 encasing the embryonic clusters (C1, C2, C3), C7 encircling the extraembryonic ectoderm (C5) and cells from C9 sparsely surround C6 and C7. Adding to this complex structure, the C8 covers endoderm layers, forming concentric rings.

The anatomical arrangement of identified clusters further refines the interpretation of potential cell fates. For example, the spatial positioning of C1 (Epiblast), C2 (Primitive Streak) and C3 (Nascent mesoderm) aligns with established developmental processes at this stage, including gastrulation and mesoderm migration (*25, 26*). Notably, C4, positioned between embryonic and extraembryonic domains, likely represents allantois and blood progenitors destined to form the umbilical cord connecting the fetus to the placenta (*27, 28*). The extraembryonic ectoderm (C5) is set to become the placental cone (*29*). Gene expression and spatial positioning identify C6 and C7 as visceral endoderm subtypes directly interacting with the embryo (*30*), with C6 more concentrated near the epiblast and C7 laterally positioned and enriched in the extraembryonic region. C9, lacking direct embryo contact and with highly specific expression of the gene encoding extracellular matrix protein - Col4a1 (fig. S29), likely represents the parietal endoderm, eventually contributing to the parietal yolk sac (*31*). The putative endothelial cells (C8) likely mark emerging blood vasculature (*32*), crucial for supporting both embryonic and extraembryonic structures. The precise enumeration and localization of minor or nascent cell groups, such as C4, C6 and C9, highlight the exceptional sensitivity of cycleHCR for *in situ* spatial analysis of cell types in intact specimens.

### cycleHCR protein imaging

To harness the high specificity of cycleHCR for multiplex protein imaging, we have developed an antibody complex that simultaneously recognizes the target and anchors a cycleHCR barcode within a polyacrylamide gel for imaging readout (Fig. 4A). Specifically, the *in vitro* assembly begins by attaching a docking sequence to the antibody using a light-activated oYo linker (*33*), followed by annealing a gel anchoring probe that binds through DNA complementarity. The gel anchoring probe carries a cycleHCR barcode and a 5’ acrydite group for gel integration. Once anchored, the barcode permanently incorporates into the gel matrix, eliminating the need for the presence of the antibody. We have developed a unique set of 25 Left and 25 Right protein cycleHCR barcodes, orthogonal to our RNA-focused set, enabling potential joint imaging of RNA and proteins. This method facilitates precise, antibody-based protein detection over multiple cycleHCR rounds in cultured cells and mouse hippocampal slices (Fig. 4A; fig. S30-31).

Embedding barcoding oligos into the gel preserves spatial information during later processing steps, making cycleHCR ideal for coupling with tissue clearing and expansion microscopy. To integrate protein cycleHCR imaging with expansion microscopy, we stabilize the expansion gel with a second, non-expandable gel overlay. This ensures stable imaging and reliable image alignment through multiple cycles (fig. S32-S34), allowing us to capture high-resolution images of 10 subcellular structures in primary mouse embryonic fibroblasts (Fig. 4B; Movie 4). 3D segmentation reveals the complex spatial organization of these structures (Fig. 4B; fig. S32-33; Movie 5), including distinct nuclear regions enriched for heterochromatin, nucleolar fibrillar centers, nuclear speckles, and active enhancers. These results highlight the potential utility of cycleHCR protein imaging in characterizing subcellular structures and their interrelations.

### Multiplex RNA and protein imaging via cycleHCR

Next, we investigated the potential of cycleHCR technology for multiplex RNA and protein imaging to elucidate cell-fate map and sub-cellular structures within the hippocampal slice. A library targeting 120 genes that label diverse cell types in the hippocampus was constructed (Table S3). A 40-60 µm thick hippocampal slice was first stained with 8 cycleHCR barcoded antibodies. After gel embedding, tissue clearing, and hybridization of the cycleHCR RNA library, 44 readout cycles were conducted over 11 days, with 40 cycles targeting RNA and 4 cycles targeting proteins. High system stability and precise cross-round image registration (fig. S35) ensured high-quality visualization of RNA and protein signals through cycleHCR rounds (Fig. 5A-D; fig. S36-S37; Movie 6). Accurate 3D cell segmentation and spot identification facilitated single-cell gene expression (fig. S38-S39; Table S4). Utilizing UMAP-based clustering, we uncovered 6 transcriptionally and spatially segregated clusters (denoted as C1-C6), corresponding to distinct neuronal and glial cell types (Fig. 5E; fig. S40-S45). Notably, we observed complex gene expression gradients along the middle line of C1-C2-C3, with 33 genes displaying descending expression levels and 7 genes showing ascending levels (Fig. 5F-G; fig. S46). The unified cycleHCR readout of RNA and protein compositions enabled cell-type characterization and sub-cellular structural investigation within the same specimen. This approach unveiled cluster-specific nuclear structural variances (Fig. 5H and fig. S47), suggestive of a potential link between cell fate and nuclear architectures.

**Fig. 5.**
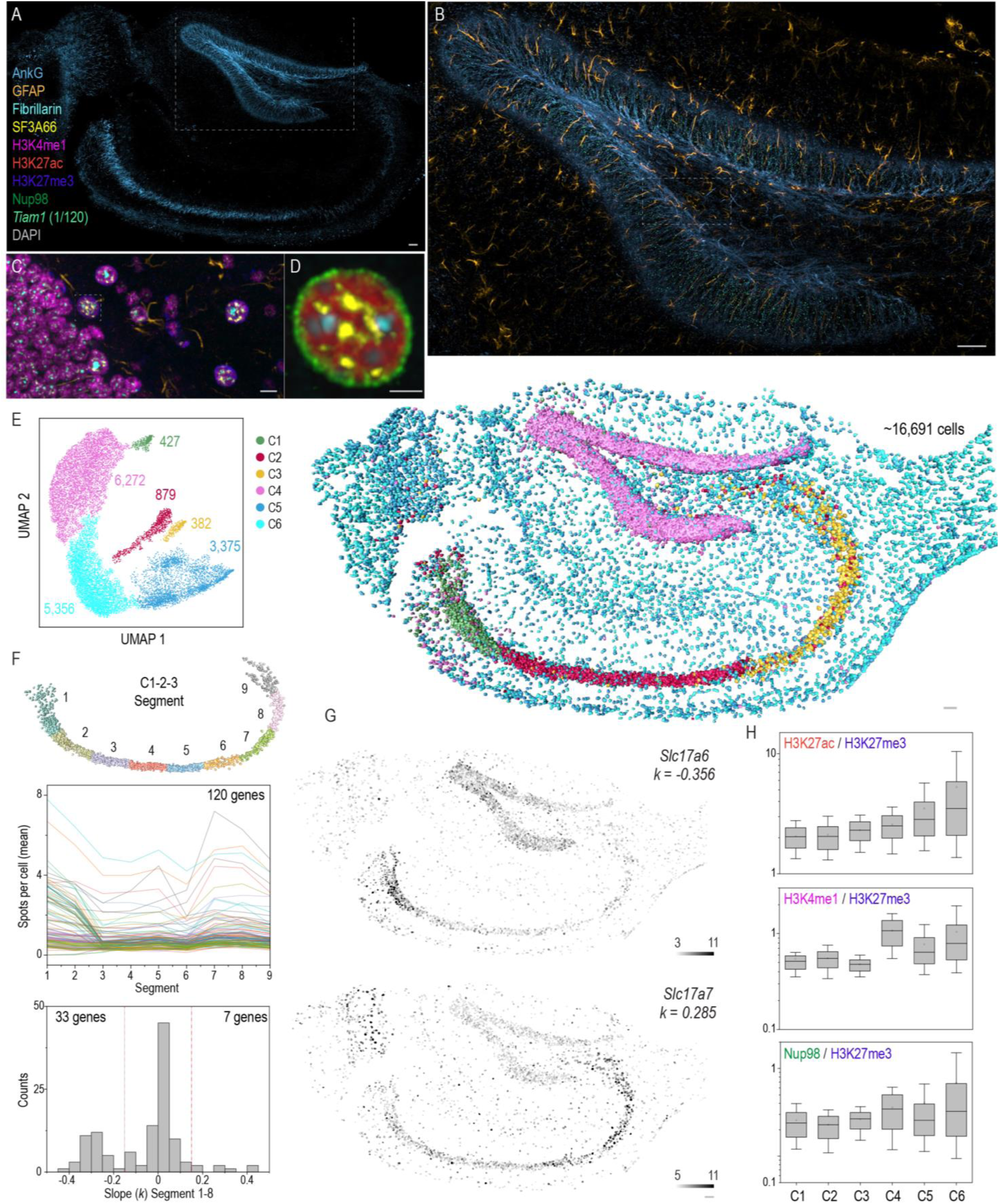
Multiplex protein and RNA imaging via cycleHCR in hippocampal slice. **(A-D)** Hierarchical images displaying structures at various length scales, with the axonal initial segment marker AnkG staining featured in (A). A closer examination of AnkG, astrocyte marker GFAP, and *Tiam1* cycleHCR RNA within the boxed region is shown in (B). Further zooming reveals Fibrillarin, SF3A66, H3K4me1, H3K27me3, and GFAP in the boxed region within (B), as depicted in (C). Zoomed-in views of Fibrillarin, SF3A66, H3K27ac, Nup98, and DAPI within the boxed region in (C) are presented in (D). Scale bars: (A-B) 50µm; (C) 10µm; (D) 5µm. (E) UMAP analysis of single-cell transcript counts identifies 6 distinct cell clusters in the hippocampus. The number of cells within each cluster is indicated in the panel, with spatial distributions of approximately 16,691 assigned cells shown on the right panel, matching cluster colors on the left panel. (F) Top: refined cells in clusters 1-3 are divided into 9 distinct segments along their well layout middle line from left to right side. Middle: The average transcript count per cell for 120 genes along Segments 1 to 9 are plotted using random colors, with multiple genes exhibiting peak gene expression at Segment 1, 7 or 8. Bottom: The distribution of linear slopes for all genes was calculated using the first 8 segments. 33 genes display decreased gene expression along segments with *k* <= -0.15, while 7 genes show increased gene expression along segments with *k* >= 0.15. (G) The solute carrier family protein *Slc17c6* showed descending gene expression levels along the C1-3 axis, while the opposite is observed for *Slc17c7*. (H) Boxplots of nuclear antigen levels (H3K27ac - active chromatin; H3K4me1 – active enhancers; nuclear pore complex subunit – Nup98) normalized to inactive chromatin (H3K27me3) intensity for all clusters. The lower and upper whiskers represent 10% and 90% values; the box represents the range from 25% to 75% percentile; the center line represents the median; the dotted line indicates the mean. The number of cells for each cluster is annotated in (E). Scale bars (E-G): 50µm.

## Discussion

Biological systems are complex networks of interconnected components, making it crucial fluorescence microscopy, constrained by fluorescent dye limitations, only allows for the observation of a limited number of components, limiting our comprehension of complex spatial regulations. Deep tissue imaging further faces challenges such as high autofluorescence and the necessity for objectives with low numerical apertures and long working distances, which are not well-suited for detecting dim single molecules. Existing imaging based spatial omics techniques require barcoding at the single-molecule level, which, combined with the need for cross-round spot-level registration, poses significant challenges in identifying targets with variable densities in thick specimens. As a result, these techniques are largely limited to thin tissue sections (*4, 5, 8*), leading to a loss of three-dimensional context.

The development of split HCR systems enhances deep tissue imaging by providing high-specificity target recognition and significant signal amplification. Built on these achievements, we introduce cycleHCR technology, which utilizes DNA barcoding to enable unified high-throughput imaging of both RNA and proteins. This approach leverages high system stability and robust image registration, ensuring accurate image alignment through multiple cycles. This synergy allows cycleHCR to map spatial information of molecular compositions and subcellular structures over successive imaging cycles, effectively overcoming the color barrier. The capability of cycleHCR to pinpoint and resolve minor or nascent cell groups in thick specimens underscores its significance in developmental biology and oncology research. Furthermore, the integration of RNA and protein imaging in a single platform, along with its compatibility with expansion microscopy, positions cycleHCR as a tool for both detailed transcriptomic studies and the characterization of high-resolution molecular architectures. This integrated approach bridges the gap between identifying cell types and understanding their sub-cellular arrangements, offering an integral view of complex biological systems.

The comparison of cycleHCR to other spatial omics methods brings to light both its unique advantages and its primary limitation, which is the lower throughput. This limitation arises mainly because of the Hybridization Chain Reaction (HCR) amplification process and its reliance on a single-shot approach for each target. However, cycleHCR uniquely bypasses the issue of molecular crowding, as it generates empirical images for both RNA and protein species without requiring spot-level registration and decoding between rounds. This is a significant advantage in studies where preserving the spatial integrity and avoiding the complications of molecular crowding are crucial.

Furthermore, the HCR amplification inherent to cycleHCR enhances signal brightness and contrast against background. This feature allows for the use of low-aperture objectives for deep imaging and rapid acquisition over large volumes. This capability is particularly valuable for studying thick tissue samples or whole organisms where both depth penetration and volume imaging are required. Future improvements in imaging depth are expected by using longer-working-distance water immersion objectives. Additionally, light-sheet microscopy (*34*), known for its efficient illumination and reduced photobleaching, could complement cycleHCR, enabling deeper imaging while preserving high resolution and reducing sample damage, further enhancing its utility.

## Supplementary Figure and Figure Legends S1 to S47

**Fig. S1.**
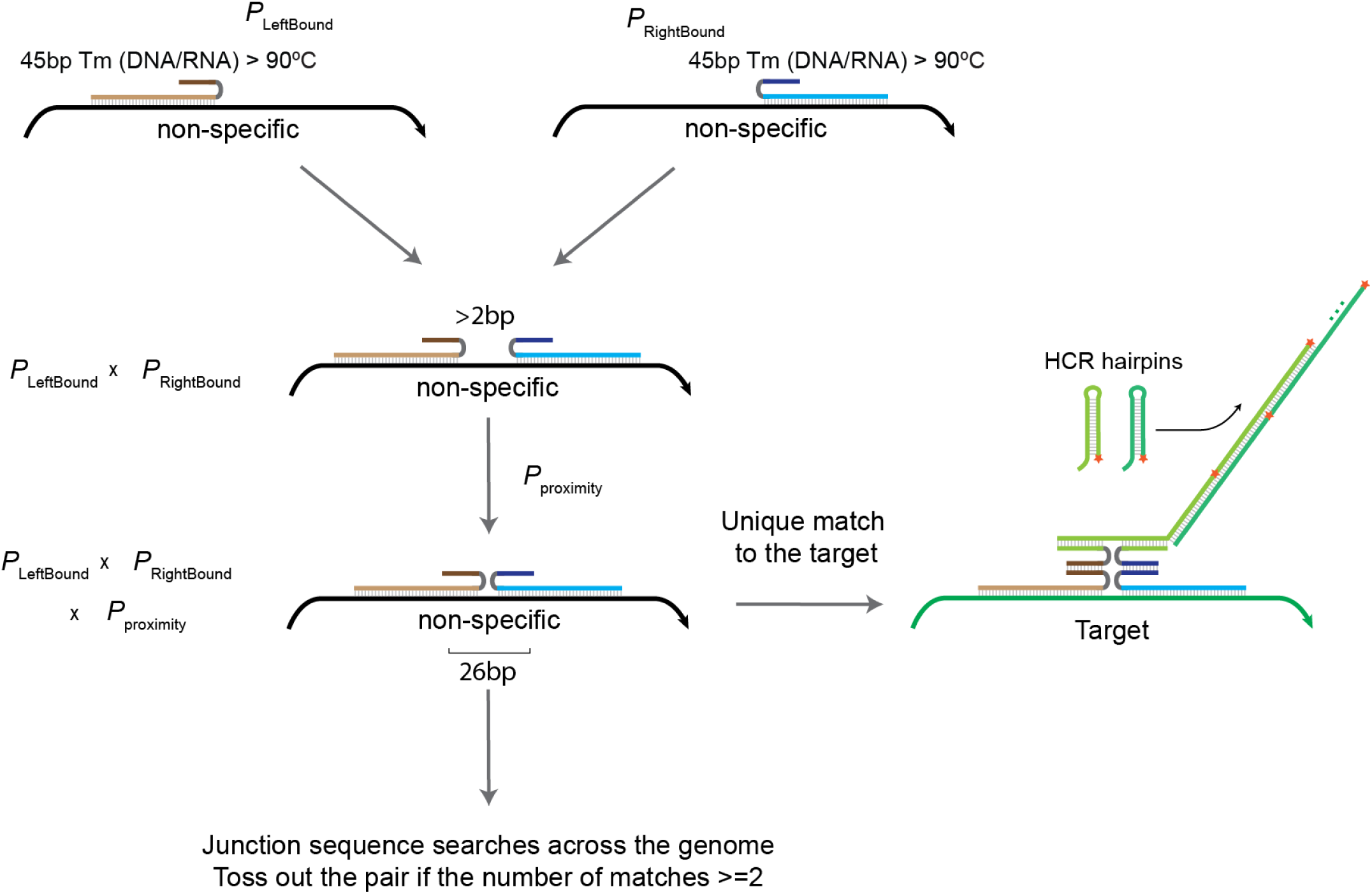
Primary probe selection for single-shot cycleHCR RNA imaging. To achieve high-specificity RNA imaging in a single-shot with cycleHCR, a multistep probe selection strategy is employed for each target transcript. This involves: 1) Sliding window search: utilizing a sliding window, we identify a 92bp sequence optimal for probe design, divided into 45bp segments for the left and right probe pairs, separated by a 2bp gap. 2) Melting temperature (Tm) estimation: The Tm for DNA-RNA probe binding is calculated to ensure that each probe half binds with a Tm above 90°C. High Tm values contribute to the stability of probe-target interactions even under stringent stripping conditions. 3) Final screening for specificity: a final specificity check involves screening for 26bp junction sequences across the genome sequence. Any probe pair with more than one genomic match is excluded to avoid non-specific targeting. To further illustrate the specificity of split primary probes, we assign the non-specific binding probabilities for both left (PLeftBound) and right (PRightBound) probes. The overall likelihood of coincidental binding to the same non-specific target is calculated as the product of PLeftBound and PRightBound. Given that HCR activation relies on the proximal binding of both probe halves, the probability of non-target HCR initiation is a product of PLeftBound, PRightBound, and the proximity probability (PProximity), highlighting the strategic approach to ensure specificity.

**Fig. S2.**
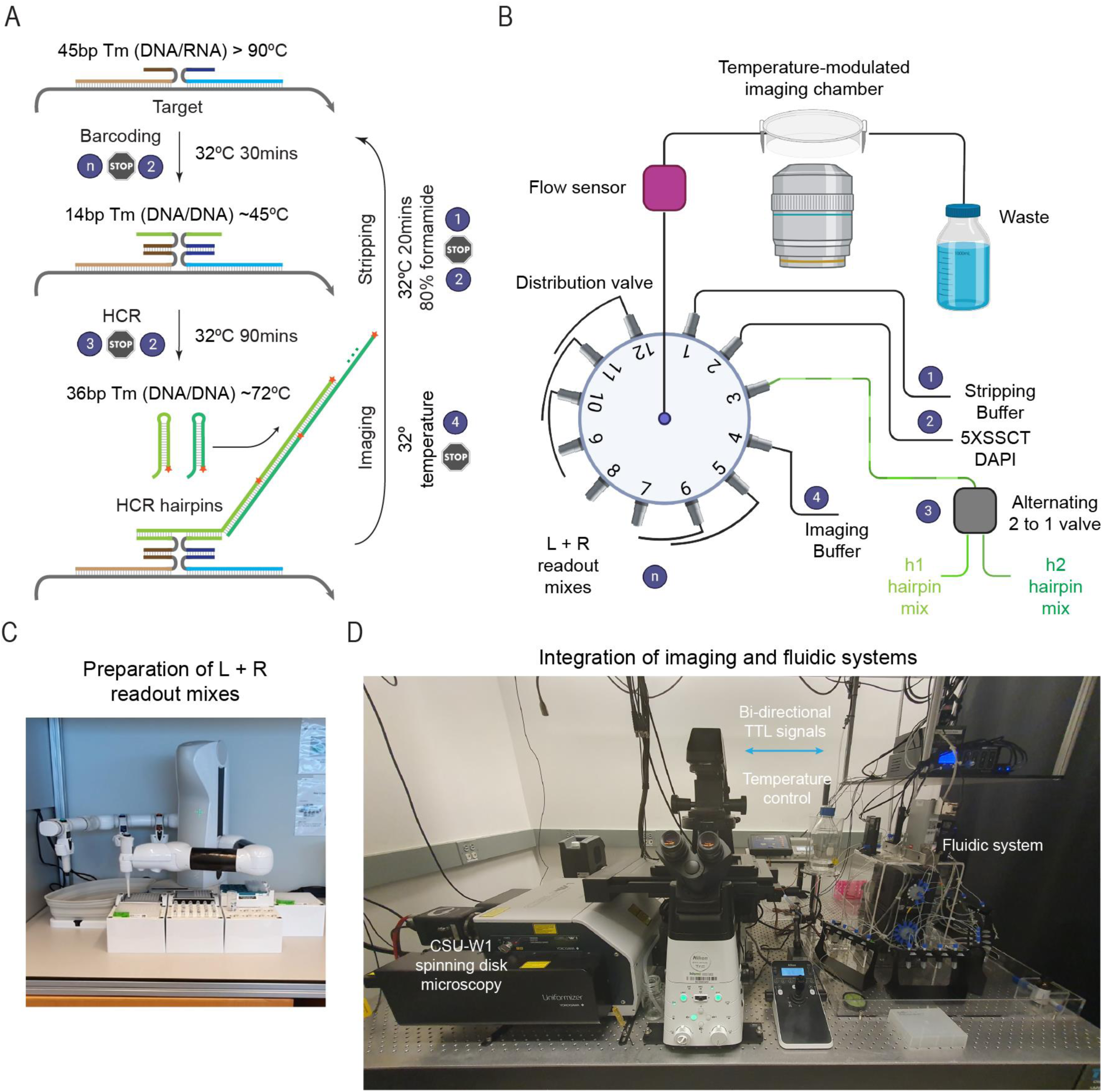
Automation of cycleHCR procedures. **(A)** The cycleHCR protocol employs a consistent incubation temperature of 32°C across all steps after primary probe hybridization. Left and right barcoding probes, each carrying split HCR initiators, are hybridized to the primary probe for 30 minutes. Following a washing step, the HCR amplification process is initiated with h1 and h2 hairpins for 1 hour and 30 minutes. Post-amplification, a final washing step precedes imaging. A tailored stripping step then selectively removes HCR hairpins and left-right barcodes without disrupting the robust primary probe:RNA interactions, due to their high melting temperatures (>90°C). **(B)** The fluidic system integrates an air pressure modulator with a flow sensor, ensuring precise control over the flow rate and volume of probes, washes, imaging, and h1/h2 solutions through a distribution valve. A 2-to-1 valve allows for real-time mixing of h1 and h2 solutions. These solutions then pass through a temperature-controlled imaging chamber before disposal into a waste collection bottle. **(C)** The preparation process of Left and Right readout mixes utilizes a programmable pipetting robotic arm. **(D)** The cycleHCR imaging setup includes a CSU-W1 spinning disk microscope, equipped with a temperature control system for both the chamber and objective, integrated with the fluidic system. Communication between the microscope and the fluidic system is facilitated via Transistor-to-Transistor Logic (TTL) communication protocol, ensuring synchronized operation between imaging and fluidic operations.

**Fig. S3.**
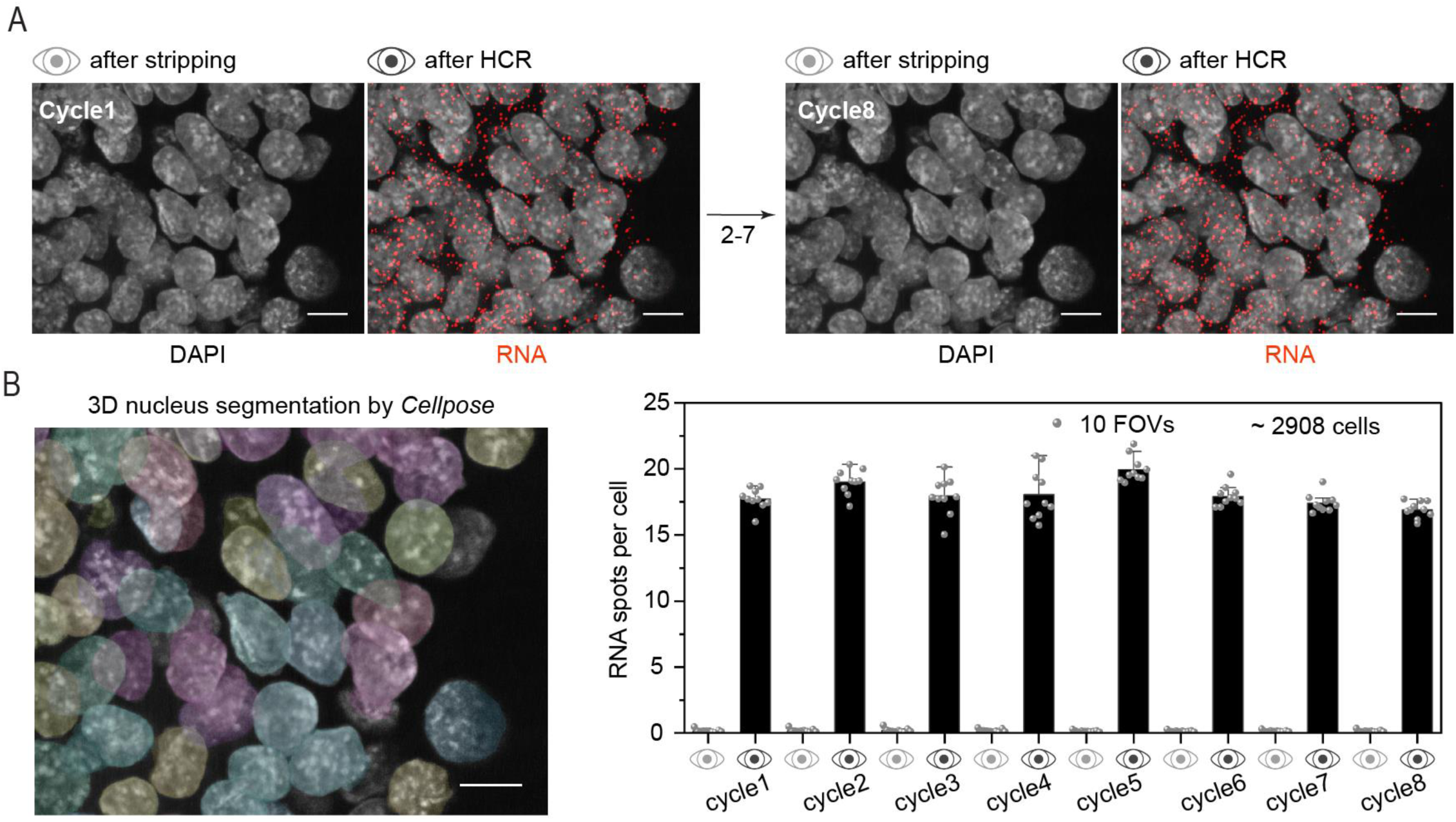
Evaluation of stripping efficiency and stability of primary probes on RNA targets. **(A)** The multicycle stripping and reprobing efficiency of cycleHCR is evaluated by repetitively stripping and reprobing the same RNA target (*Trim6*; L2+R2 at 561nm) across eight cycles. For each cycle, imaging data were captured post-stripping and post-HCR to assess the efficiency of probe removal and the consistency of signal amplification. **(B)** The stability of primary probe-RNA interactions was quantitatively evaluated using nuclear segmentation with Cellpose (left panel), enhanced by a custom-trained model, alongside 3D single molecule localization with RS-FISH. RNA spot counts were determined for each cell within 10 fields of view, totaling approximately 2,908 cells. The analysis revealed no significant decrease in signal intensity across the eight cycles (right panel). Scale bars: 10µm.

**Fig. S4.**
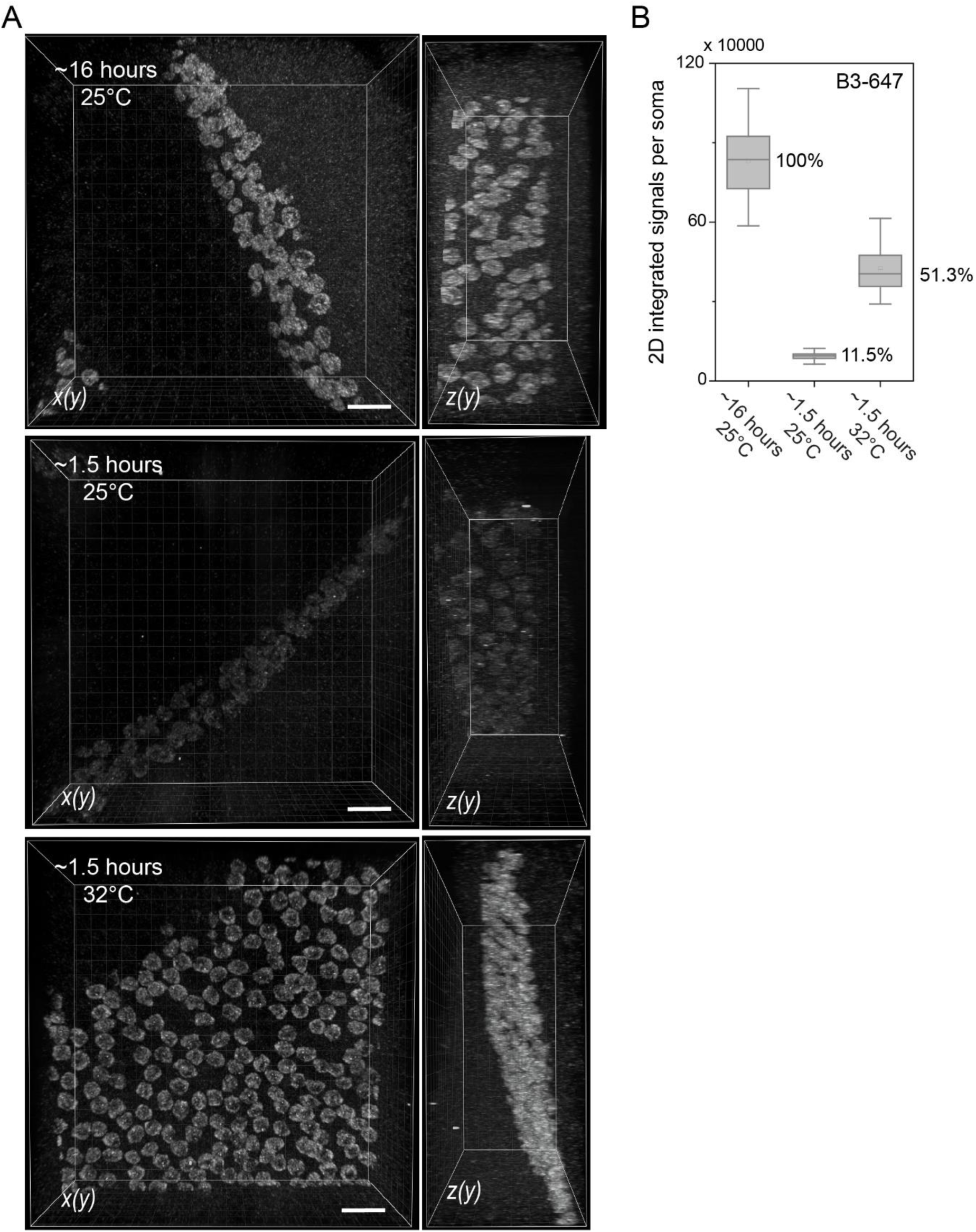
Optimizing amplification temperature for cycleHCR. **(A)** To optimize the cycleHCR protocol, we evaluated the impact of different amplification temperatures and durations on signal performance using a cerebellar section of approximately 200 µm thickness. *Rgs8* mRNA was repeatedly stripped and reprobed under various temperature conditions to assess signal intensity variations. Imaging was performed to document signal differences across these conditions. 3D Imging volumes are rendered by Imaris using MIP mode with the same intensity range for all images. Scale bars: 50µm. **(B)** Comparative analysis revealed that HCR amplification at 32°C for 1.5 hours achieved signal intensities about 51% of those obtained through overnight amplification at ambient room temperature (∼25°C). In comparison, a 1.5-hour amplification at 25°C resulted in only approximately 11% of the signal intensity relative to the overnight conditions. Signal quantification involved integrating signals above background levels per cell in 2D slices, with approximately 50 quantifications conducted for each condition. The upper and lower whiskers represent 90% and 10% values; the box represents the range from 25% to 75% percentile; the center line represents the median; the dotted line indicates the mean.

**Fig. S5.**
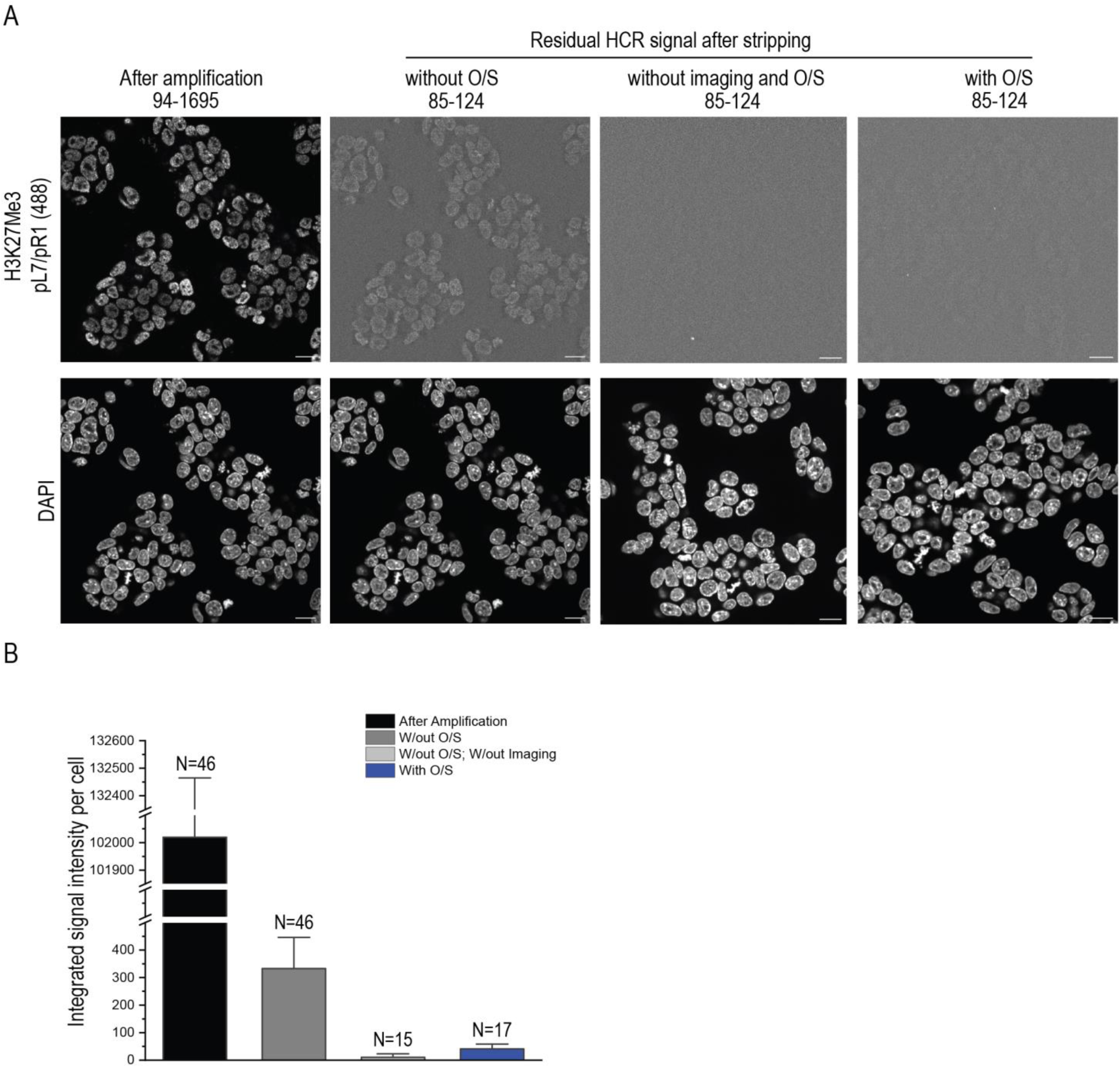
Mitigating photo-crosslinking in cycleHCR imaging using oxygen scavenger. **(A)** To evaluate photo-crosslinking effect, samples following protein cycleHCR targeting histone H3K27me3 underwent imaging at 100% laser power. Then, continuous illumination for 15 seconds was then applied both with and without the presence of oxygen scavenger (O/S) to determine its effect on photo-crosslinking of HCR fluorescent probes to the specimen. Fluorescence images captured after probe stripping allowed for the assessment of photo-crosslinking levels. The presence of O/S significantly reduces photo-crosslinking. The display range for these images is provided above. Scale bars: 20µm. **(B)** The impact of O/S on reducing photo-crosslinking was quantitatively evaluated by measuring residual fluorescence signals exceeding background levels per cell in 2D slices. The analysis confirms the effectiveness of oxygen scavenger in minimizing undesired photo-crosslinking effects. The error bars represent standard deviations.

**Fig. S6.**
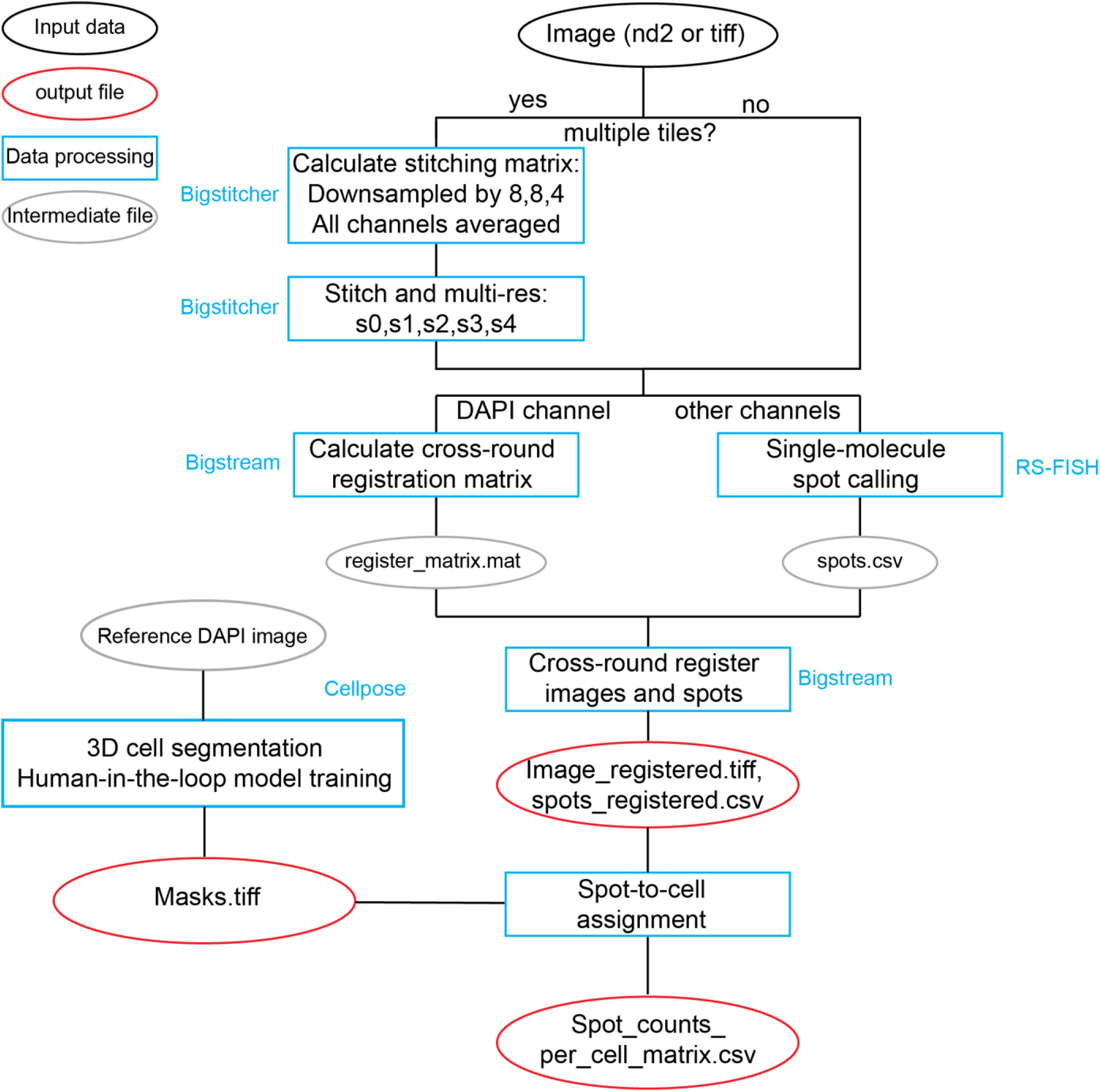
Nextflow orchestrated image processing workflow for cycleHCR. This figure presents a workflow designed for efficient processing of imaging data. The workflow is orchestrated with Nextflow and includes the following steps: 1) Initial categorization: raw image files, in Nd2 and Tiff formats, are first categorized based on the number of tiles present at each time point. 2) Image stitching: for instances where the number of tiles exceeds one, images undergo stitching via BigStitcher. Conversely, single-tile images bypass this stitching step. 3) RNA localization: RS-FISH is utilized to identify mRNA single molecule localizations 4) Cross-round registration: The BigStream tool calculates cross-round registration matrices, enabling the alignment of images and localization spots across different rounds using DAPI channel data for consistent registration. 5) Cell segmentation: in parallel with registration, cell segmentation is conducted using Cellpose using the reference DAPI channel, employing a custom model refined through a human-in-the-loop method to ensure accurate segmentation of nucleus boundaries. 6) Spot-to-cell assignment: following registration, localizations are assigned to each segmented cell, culminating in a comprehensive localization count per cell for all identified cell masks within the image.

**Fig. S7.**
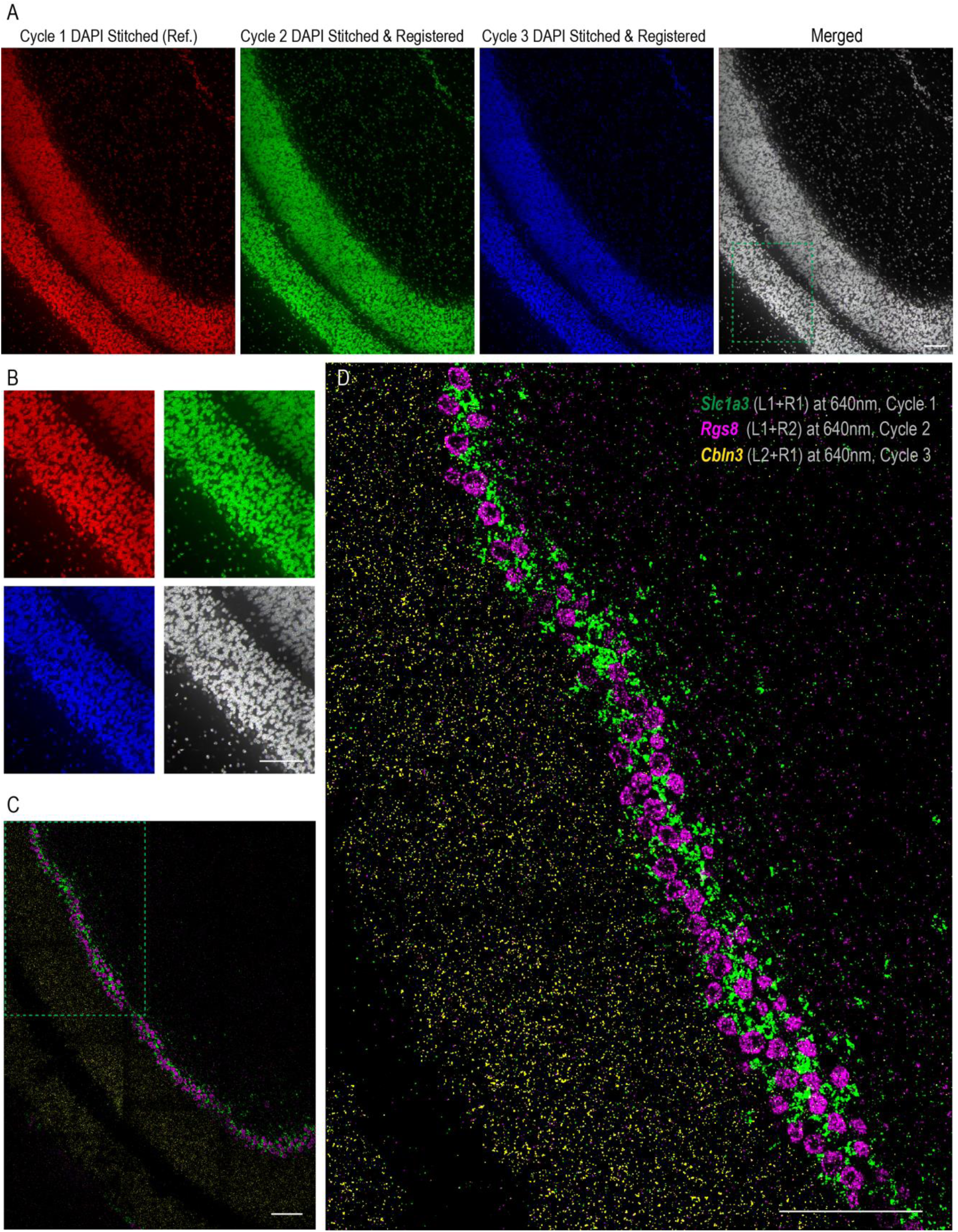
Cross-round registration validation in mouse cerebellar slice imaging. **(A)** Registration validation shows the alignment post-stitching, with the reference DAPI image (in red) overlapping with DAPI images from Cycle 2 (in green) and Cycle 3 (in blue) for the sequential three round cycleHCR showed in Figure 1E. **(B)** A closer inspection within the marked box in (A) reveals the precise overlaps of DAPI signals throughout the three cycleHCR imaging consecutive cycles. **(C)** An image displays the sequential cycleHCR RNA signals for three genes: *Slc1a3* (L1+R1 at 640nm), *Rgs8* (L1+R2 at 640nm), and *Cbln3* (L2+R1 at 640nm) across three cycles. **(D)** Enhanced view of RNA Localization by further zooming into the region highlighted in (C) provides a detailed look at the RNA signals for the specified genes. Scale bars, 100µm.

**Fig. S8.**
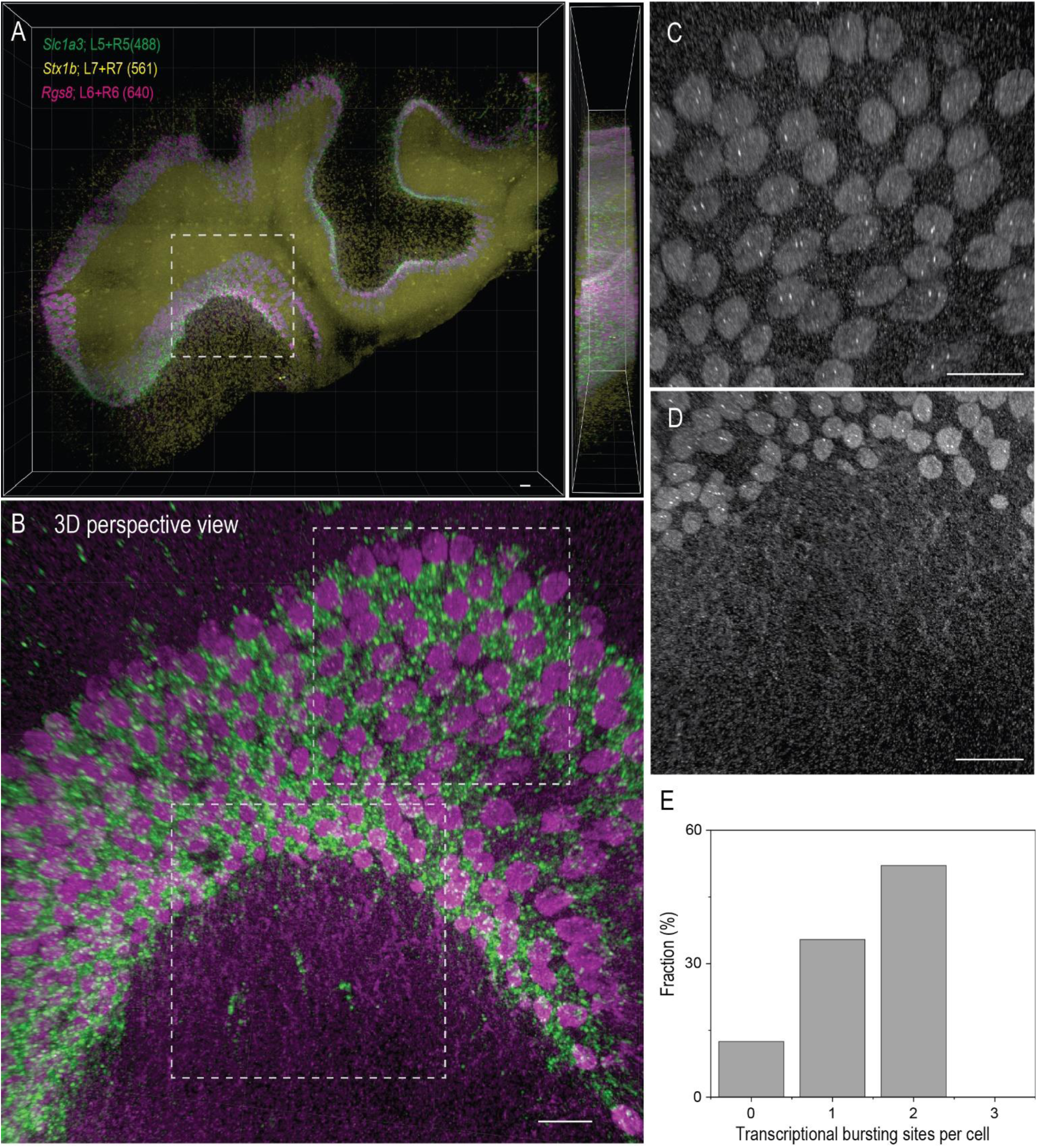
Detailed sub-cellular RNA imaging in thick cerebellum slice via cycleHCR. **(A)** Utilizing a three-color, single-round cycleHCR approach, we visualize RNAs from three genes specific to different cell types within a cerebellum slice measuring 2.4mm x 1.8mm x 0.2mm. The genes imaged include *Slc1a3* (L5+R5 at 488nm) for Bergmann glia cells, *Stx1b* (L7+R7 at 561nm) for the granule layer, and *Rgs8* (L6+R6 at 640nm) for Purkinje cells. This panel shows the capability of cycleHCR to delineate gene expression patterns across cell types in thick tissue sections. **(B)** A three-dimensional perspective view of *Slc1a3* (green) and *Rgs8* (magenta) RNA localizations within a specified region from (A). **(C)** cycleHCR reveals transcriptional bursting sites of the *Rgs8* gene within Purkinje cells, visible in a designated upper box region in (B). **(D)** *Rgs8* mRNA molecules localize along the dendrites of Purkinje cells, as shown in the lower box region in (B). This detail emphasizes the ability of cycleHCR to map mRNA distributions within specific cellular compartments. **(E)** The statistical analysis of the number of transcriptional bursting sites of *Rgs8* gene (Chr 1) per cell across populations of Purkinje cells, with approximately 200 cells quantified. Images are rendered using the 3D perspective view of maximal intensity projection (MIP) with Imaris. Scale bars: 50µm.

**Fig. S9.**
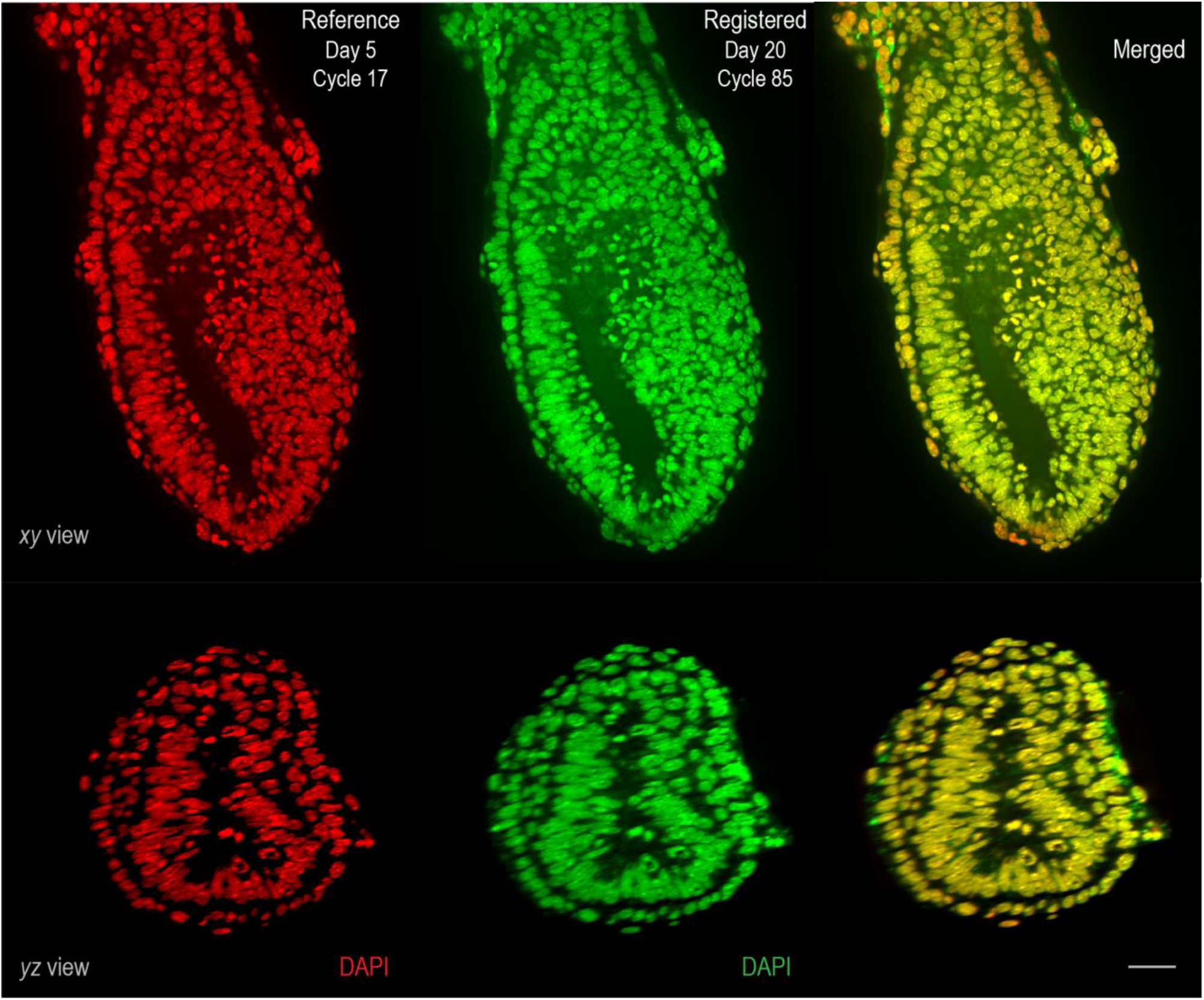
Verifying cross-cycle registration in whole-embryo transcriptomics imaging. This figure demonstrates the reliability and precision of our image registration throughout the process of whole-embryo transcriptomics imaging, as detailed in Figures 2-3. By comparing the reference DAPI image from Cycle 17 (presented in Red) with that from the final Cycle 85 (shown in Green), both in *xy* and *yz* planes, we observe the excellent alignment achieved between these disparate imaging cycles. The consistent overlay of the two images indicates both the stability of our imaging setup and the efficacy of the registration method employed across multiple cycleHCR rounds, ensuring accurate and coherent data integration throughout the study. Scale bar: 50µm.

**Fig. S10.**
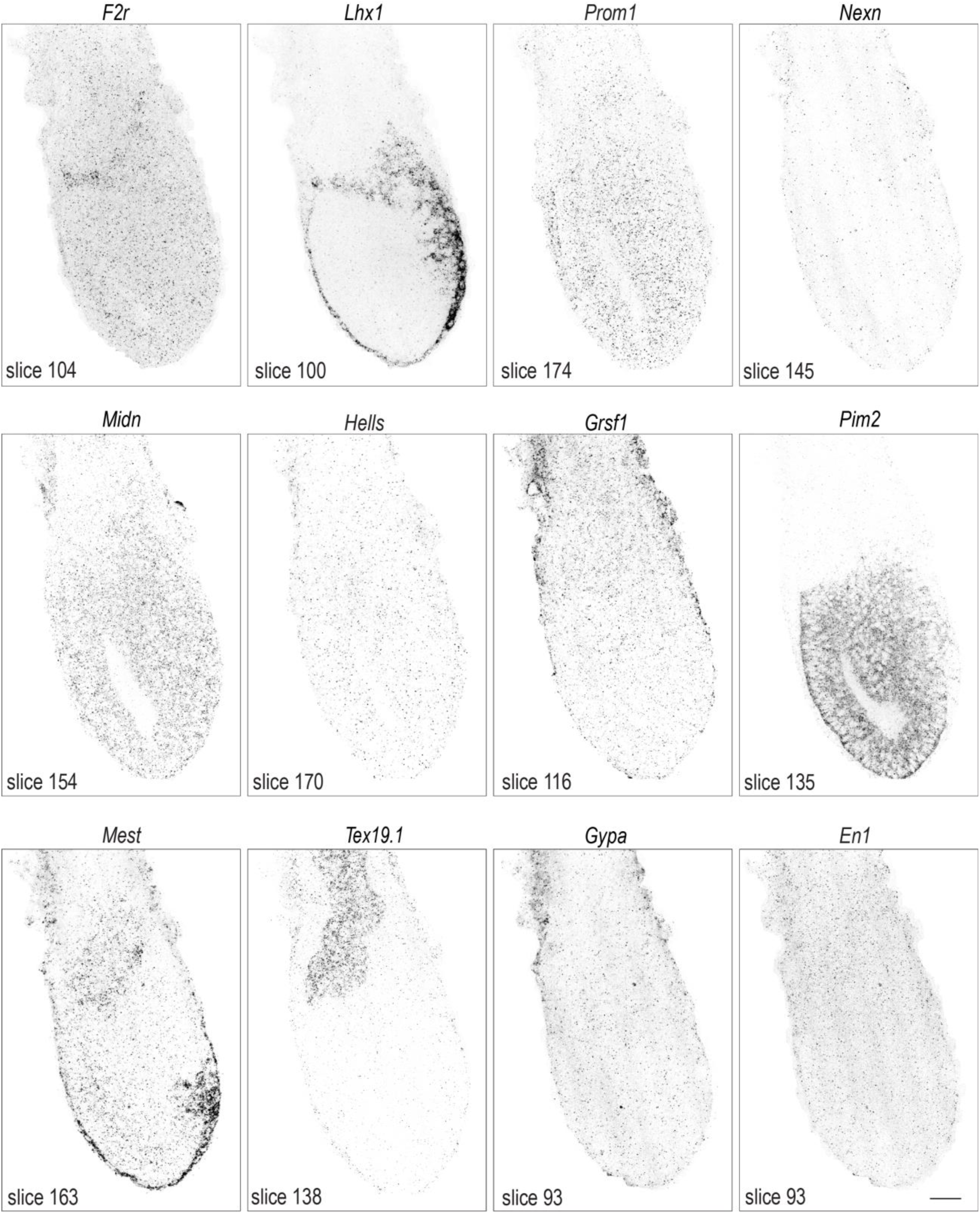
Whole-embryo transcriptomics imaging – gene panel 1. This figure features a series of inverted raw images captured during whole-embryo transcriptomics imaging, highlighting the mRNA expression patterns of indicated genes detected by the cycleHCR technique. Each panel displays a slice of the embryo, with the slice number indicated. The images are taken at 1 µm intervals. Scale bar: 50µm.

**Fig. S11.**
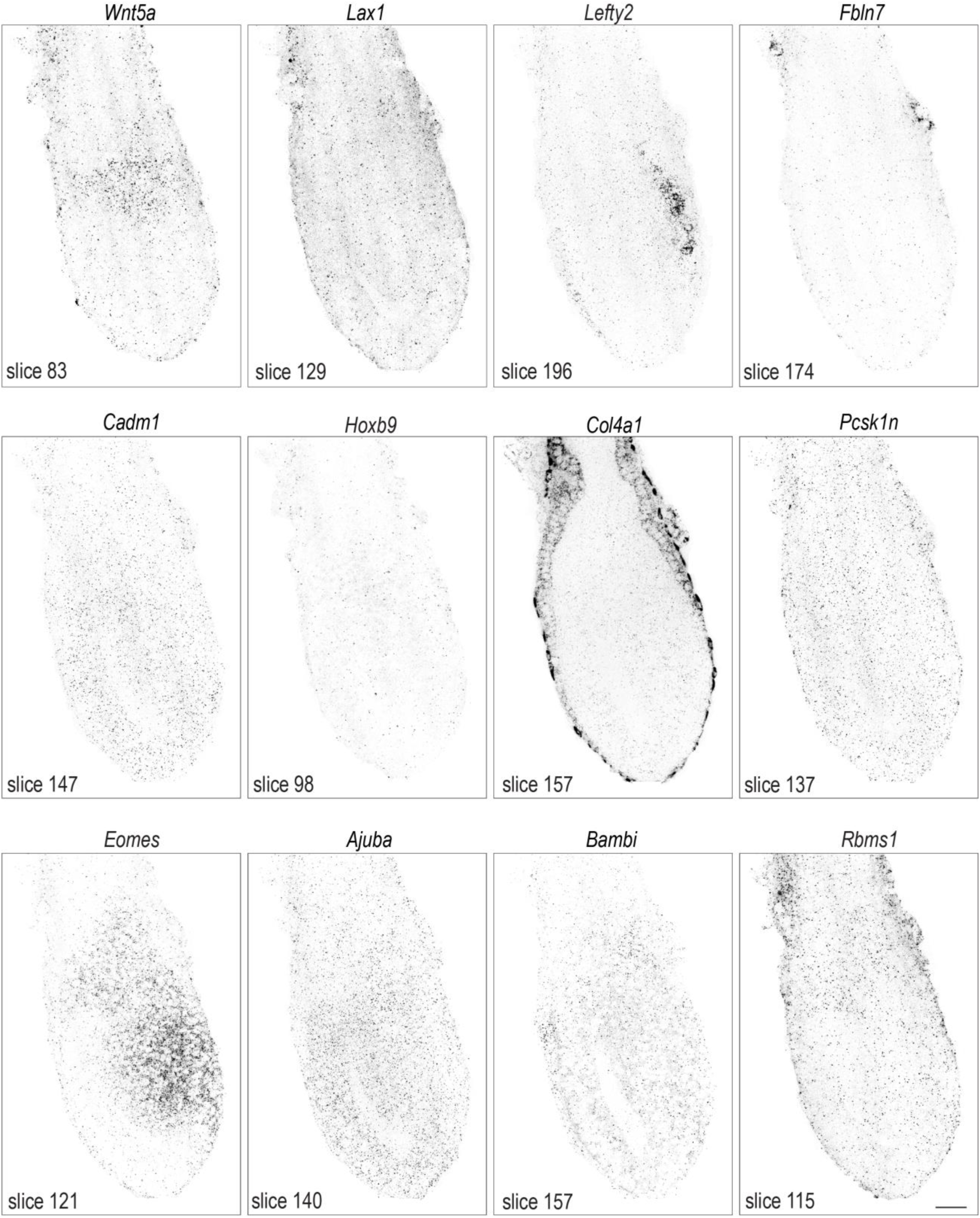
Whole-embryo transcriptomics imaging – gene panel 2. This figure features a series of inverted raw images captured during whole-embryo transcriptomics imaging, highlighting the mRNA expression patterns of specified genes detected by the cycleHCR technique. Each panel displays a slice of the embryo, with the slice number indicated. The images are taken at 1 µm intervals. Scale bar: 50µm.

**Fig. S12.**
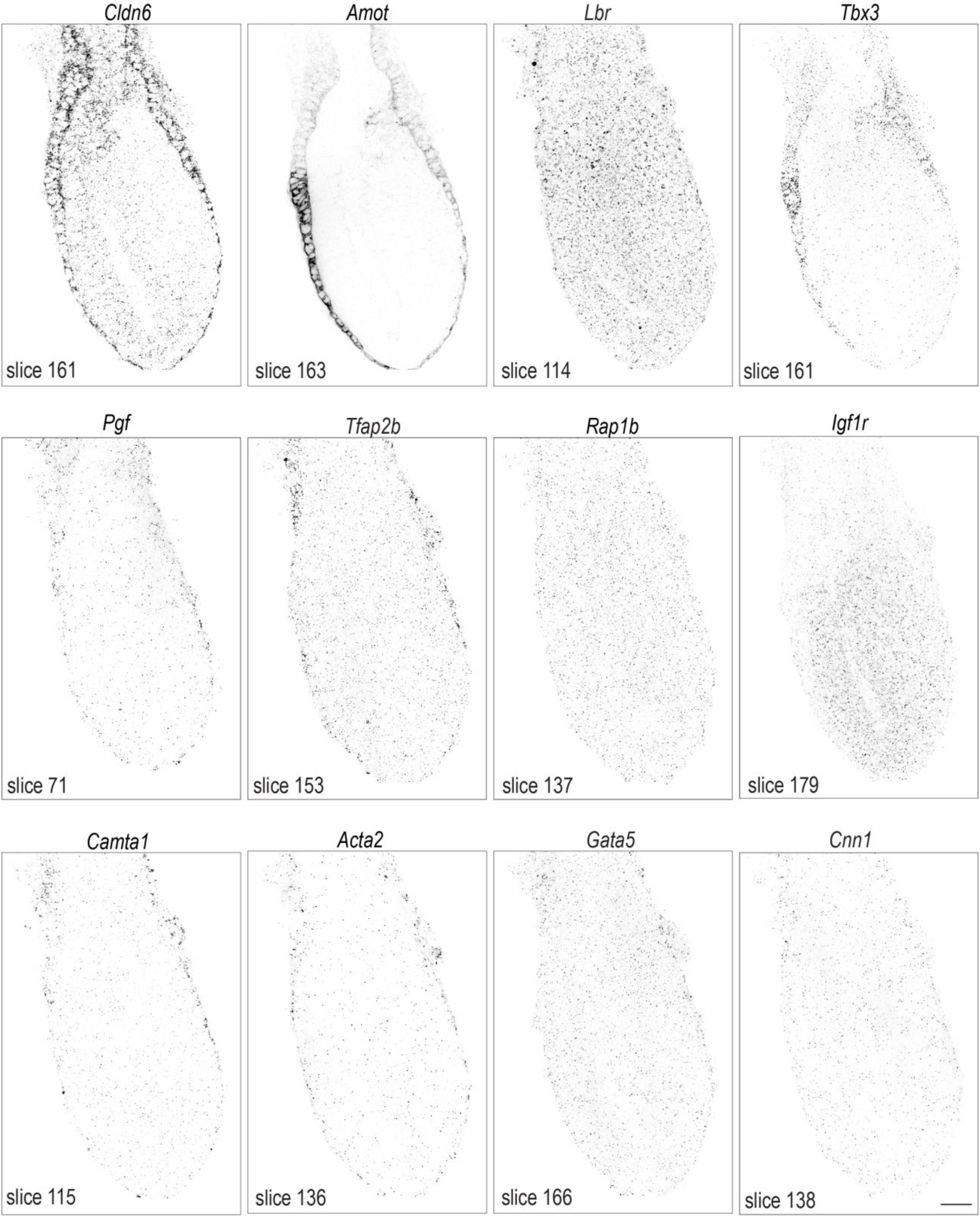
Whole-embryo transcriptomics imaging – gene panel 3. This figure features a series of inverted raw images captured during whole-embryo transcriptomics imaging, highlighting the mRNA expression patterns of specified genes detected by the cycleHCR technique. Each panel displays a slice of the embryo, with the slice number indicated. The images are taken at 1 µm intervals. Scale bar: 50µm.

**Fig. S13.**
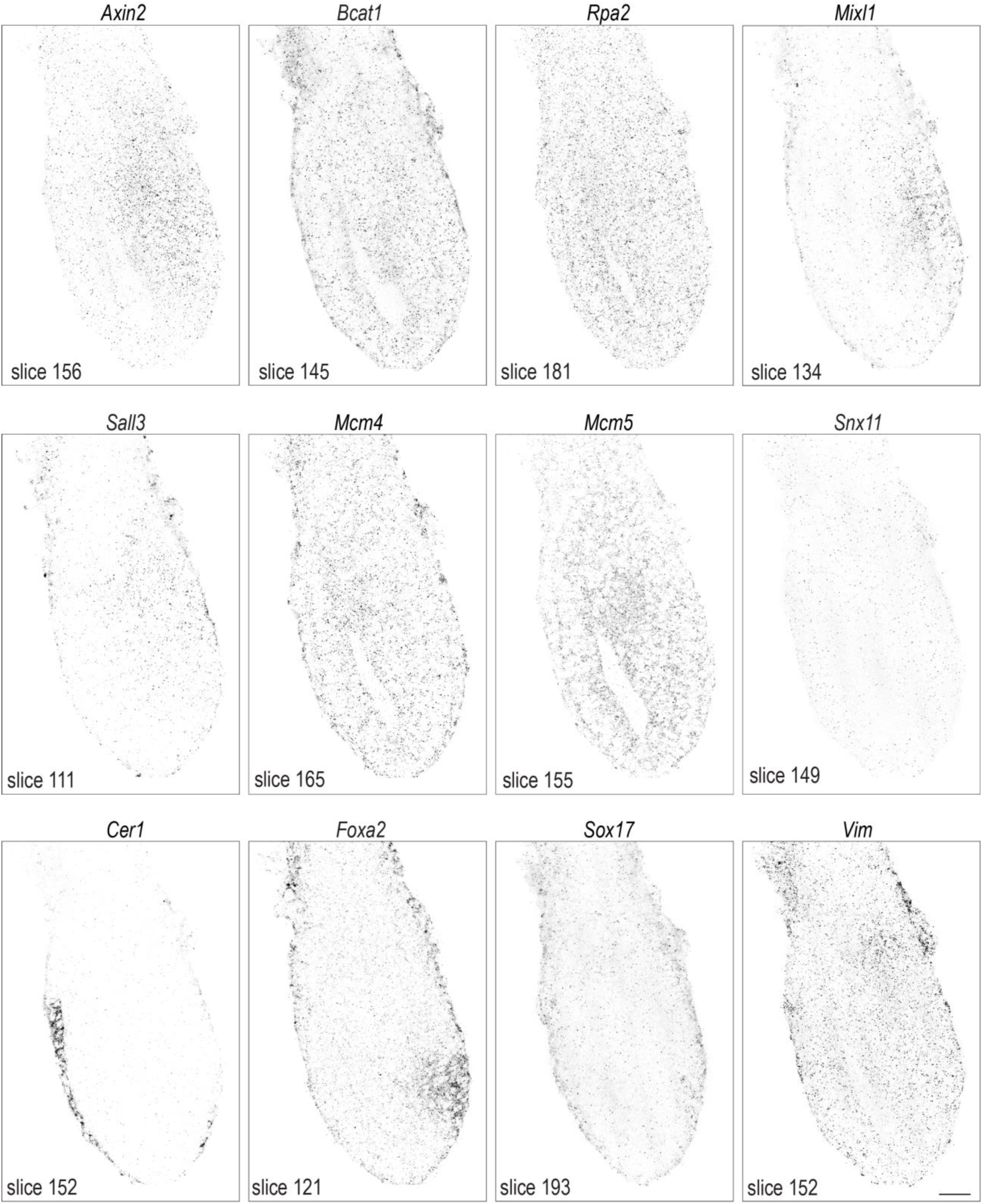
Whole-embryo transcriptomics imaging – gene panel 4. This figure features a series of inverted raw images captured during whole-embryo transcriptomics imaging, highlighting the mRNA expression patterns of specified genes detected by the cycleHCR technique. Each panel displays a slice of the embryo, with the slice number indicated. The images are taken at 1 µm intervals. Scale bar: 50µm.

**Fig. S14.**
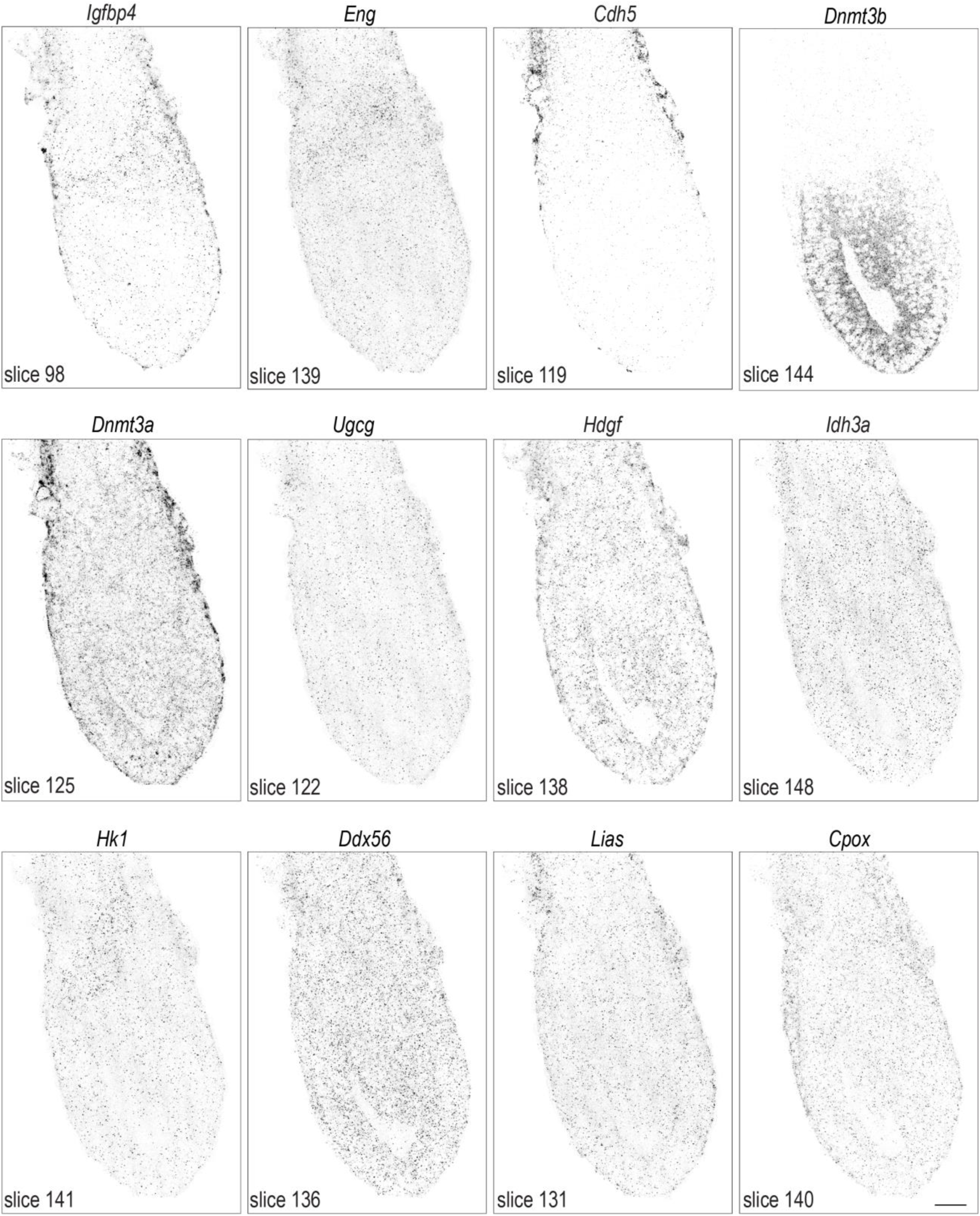
Whole-embryo transcriptomics imaging – gene panel 5. This figure features a series of inverted raw images captured during whole-embryo transcriptomics imaging, highlighting the mRNA expression patterns of specified genes detected by the cycleHCR technique. Each panel displays a slice of the embryo, with the slice number indicated. The images are taken at 1 µm intervals. Scale bar: 50µm.

**Fig. S15.**
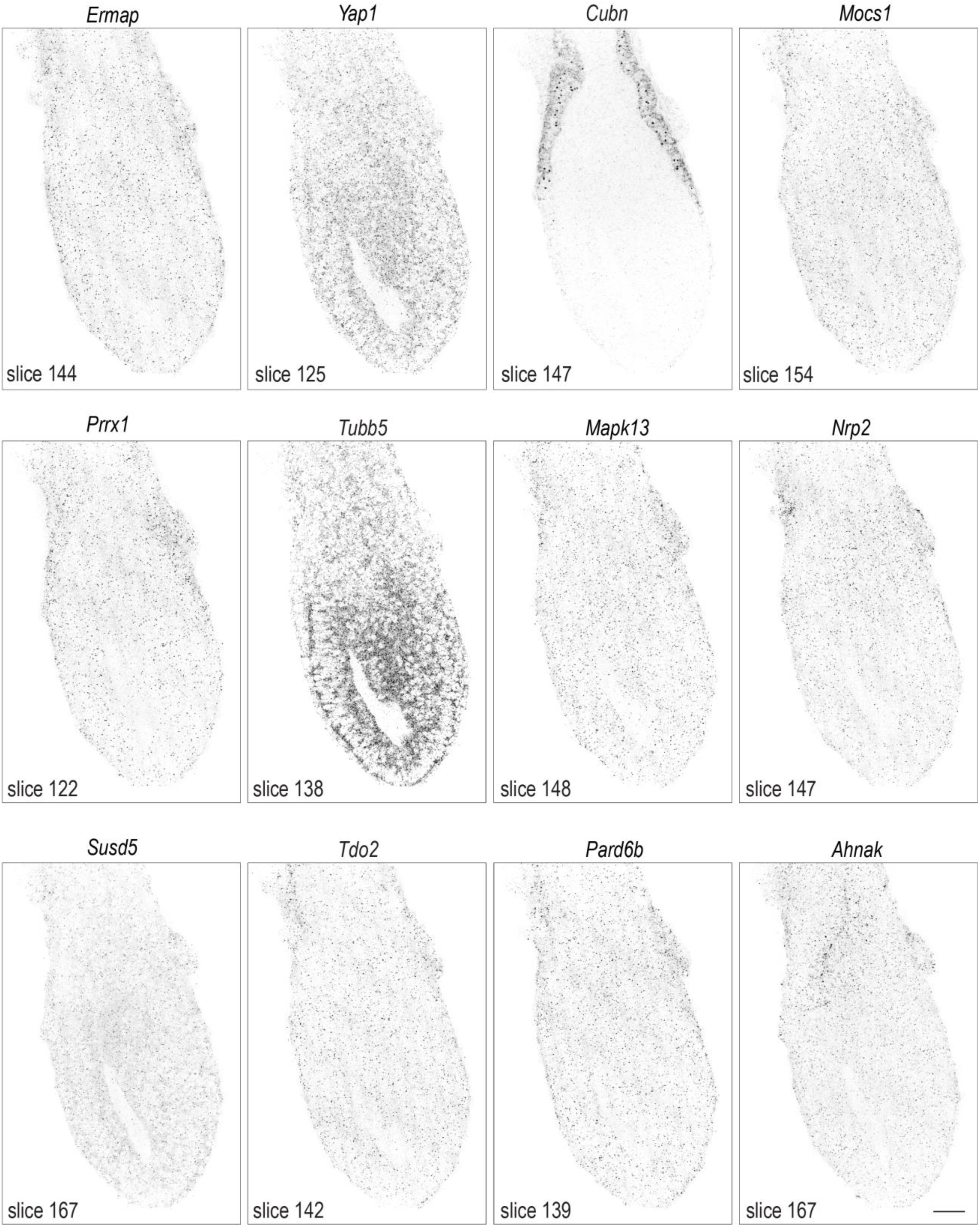
Whole-embryo transcriptomics imaging – gene panel 6. This figure features a series of inverted raw images captured during whole-embryo transcriptomics imaging, highlighting the mRNA expression patterns of specified genes detected by the cycleHCR technique. Each panel displays a slice of the embryo, with the slice number indicated. The images are taken at 1 µm intervals. Scale bar: 50µm.

**Fig. S16.**
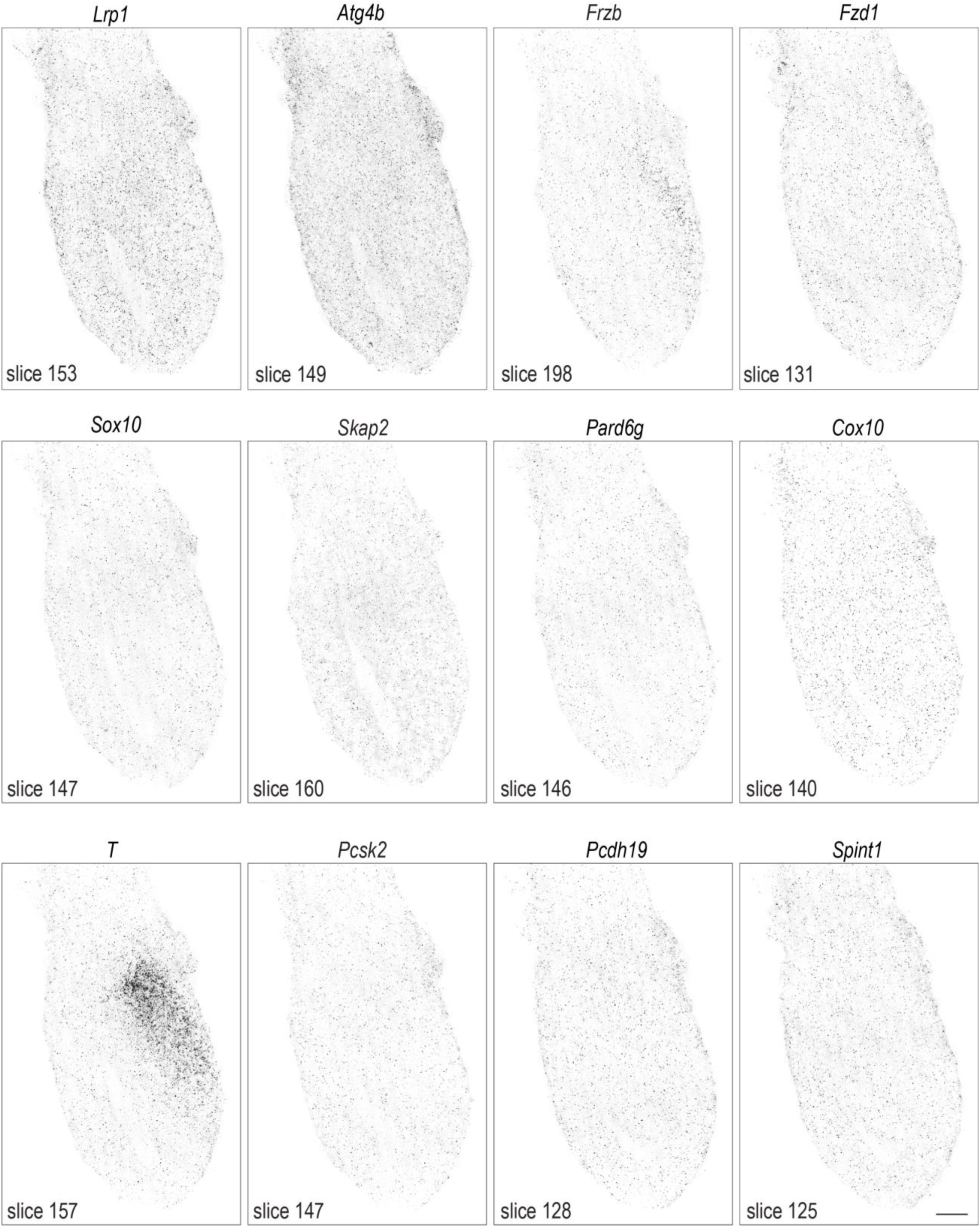
Whole-embryo transcriptomics imaging – gene panel 7. This figure features a series of inverted raw images captured during whole-embryo transcriptomics imaging, highlighting the mRNA expression patterns of specified genes detected by the cycleHCR technique. Each panel displays a slice of the embryo, with the slice number indicated. The images are taken at 1 µm intervals. Scale bar: 50µm.

**Fig. S17.**
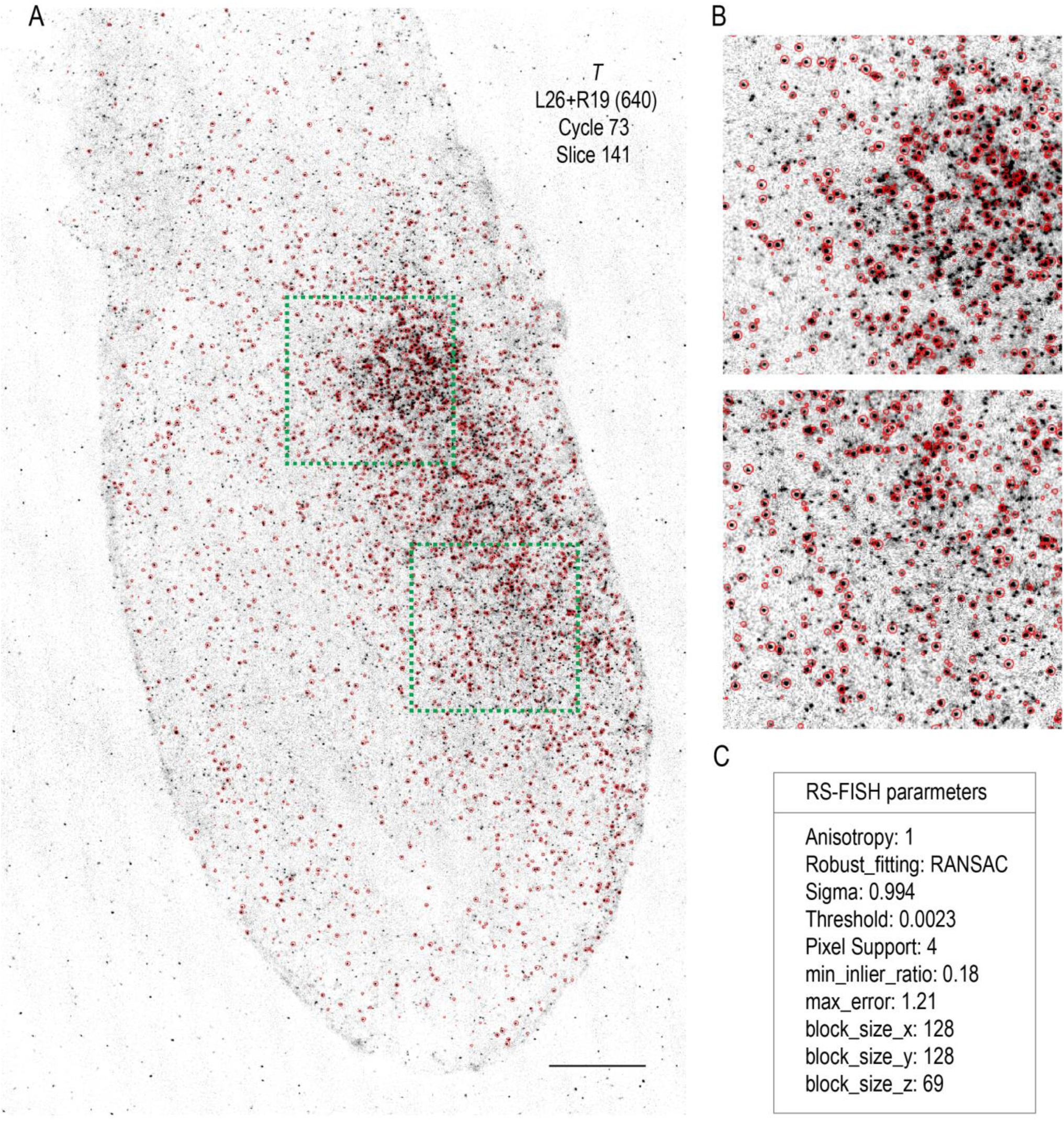
Single-molecule localization for whole-embryo transcriptomics imaging. **(A)** This panel demonstrates the precision of single-molecule localization in imaging mRNA transcripts of *T* gene. Red circles denote the positions of localized single molecules detected by RS-FISH, overlaid on an inverted raw image captured using the cycleHCR barcode L26 + R19 at the 640nm channel at cycle 73. Scale bar: 50µm. **(B)** A zoomed-in view of areas within green boxes from (A) provides a closer examination of the single-molecule localization specificity, highlighting the ability of cycleHCR in capturing transcriptomic details at the molecular level. **(C)** Single-molecule localization parameters with RS-FISH for localizing single-molecules. These parameters were selected to ensure minimal false positive detections.

**Fig. S18.**
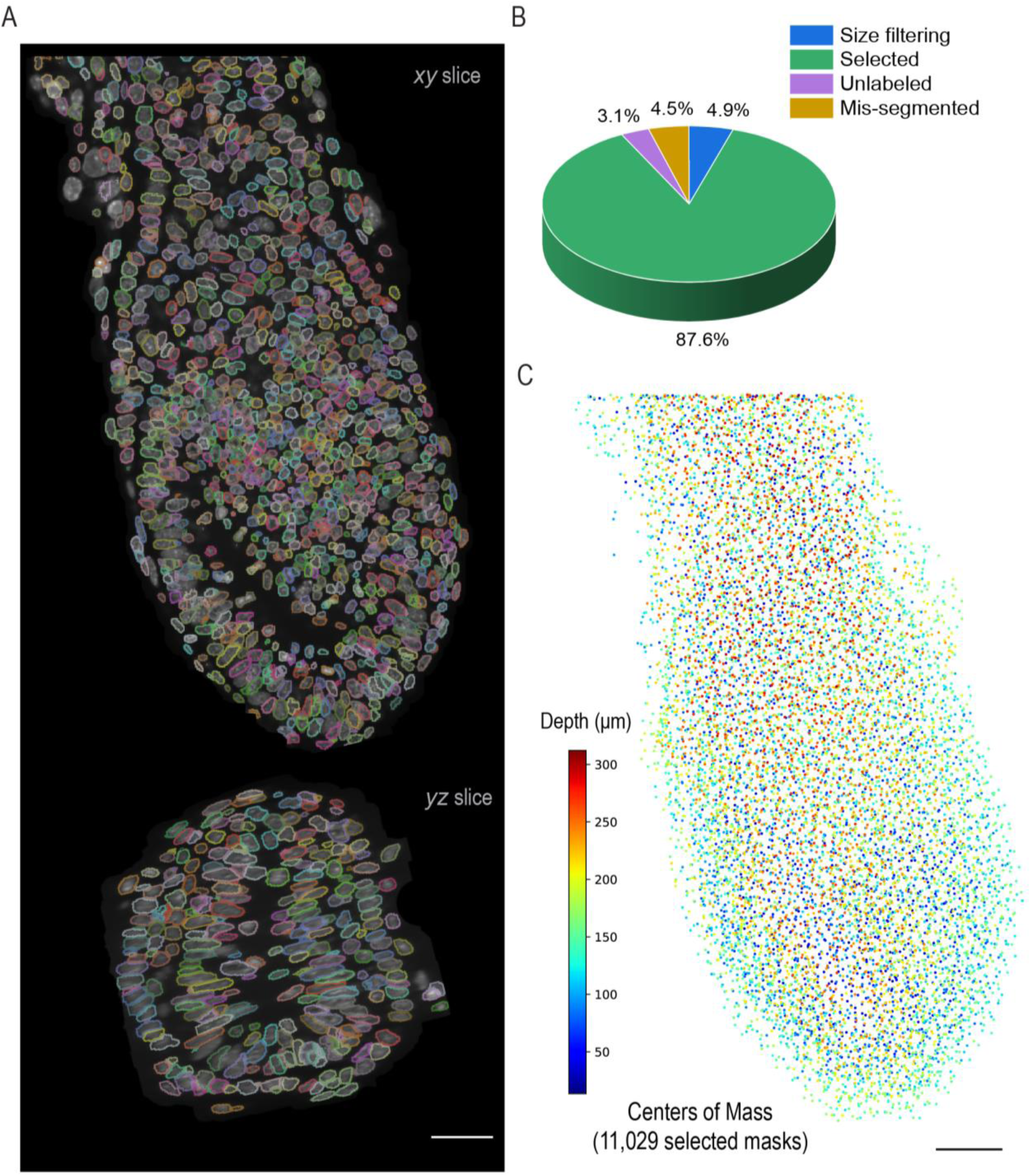
3D cell segmentation with a specialist model by Cellpose. **(A)** This panel presents both *xy* and yz slice views of the segmented cells within the embryo by Cellpose using a human-in-the-loop custom trained model. The contours delineate DAPI stained nuclei, allowing for detailed examination of cell segmentation result. The images were rendered using ORS dragonfly. **(B)** A pie chart details the results of the cell segmentation, categorizing the masks into groups: those filtered out by size criteria, selected, unlabeled, and mis-segmented. **(C)** A scatterplot representing the centers of mass for all 11,029 selected masks, with the *z*-axis depth of each point color-coded according to the accompanying color bar.

**Fig. S19.**
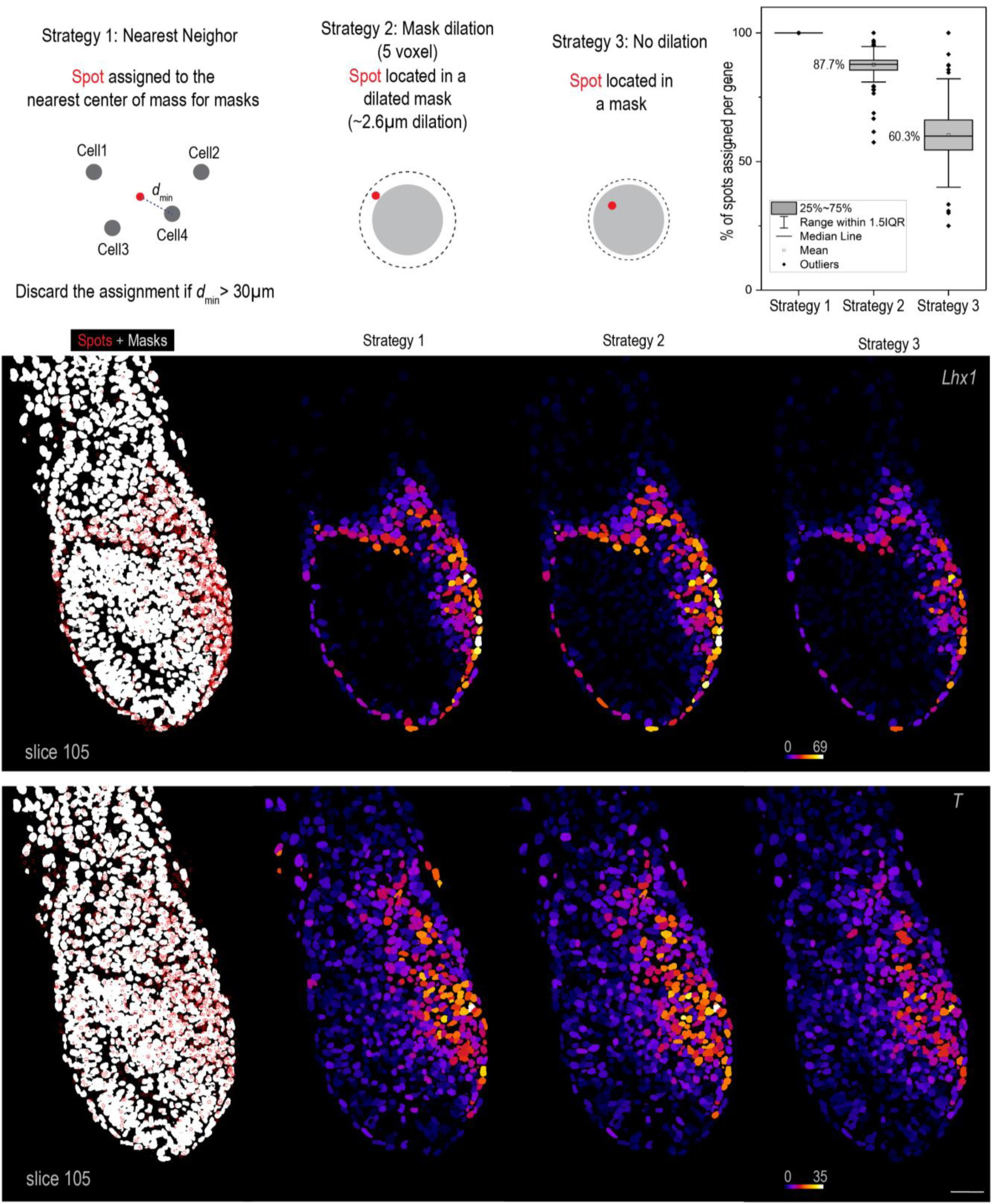
Spot-to-cell assignment and visualization. Following single-molecule localization of cycleHCR RNA signals and cell segmentation outlined in Fig. S17 and S18, three spot-to-cell assignment strategies were evaluated. The first strategy assigns each RNA spot to the nearest center of mass for masks, discarding assignments if the minimal distance exceeds 30 µm. The second strategy utilizes dilated masks (achieved through 5 voxel iterative non-overlapping dilations in an isotropically resampled volume with 0.52 µm voxel size), while the third assigns spots based on original masks without dilation. It is worth noting that cellpose masks are slightly dilated compared to DNA counterstained nuclei (Fig. 2E). The percentage of spots assigned per gene per strategy is showed in the box plot (n = 254) with mean levels noted in the panel. Visual representation includes localized RNA spots as red dots against white-colored nucleus masks (Left panel). Spot counts are integrated into cell mask labels within the image stack. Representative *xy* slice views (slice 105) for genes *Lhx1* and *T* under different assignment strategies show single-cell gene expression levels, color-coded for each cell according to a color map located at the bottom left of the image. Scale bar: 50µm.

**Fig. S20.**
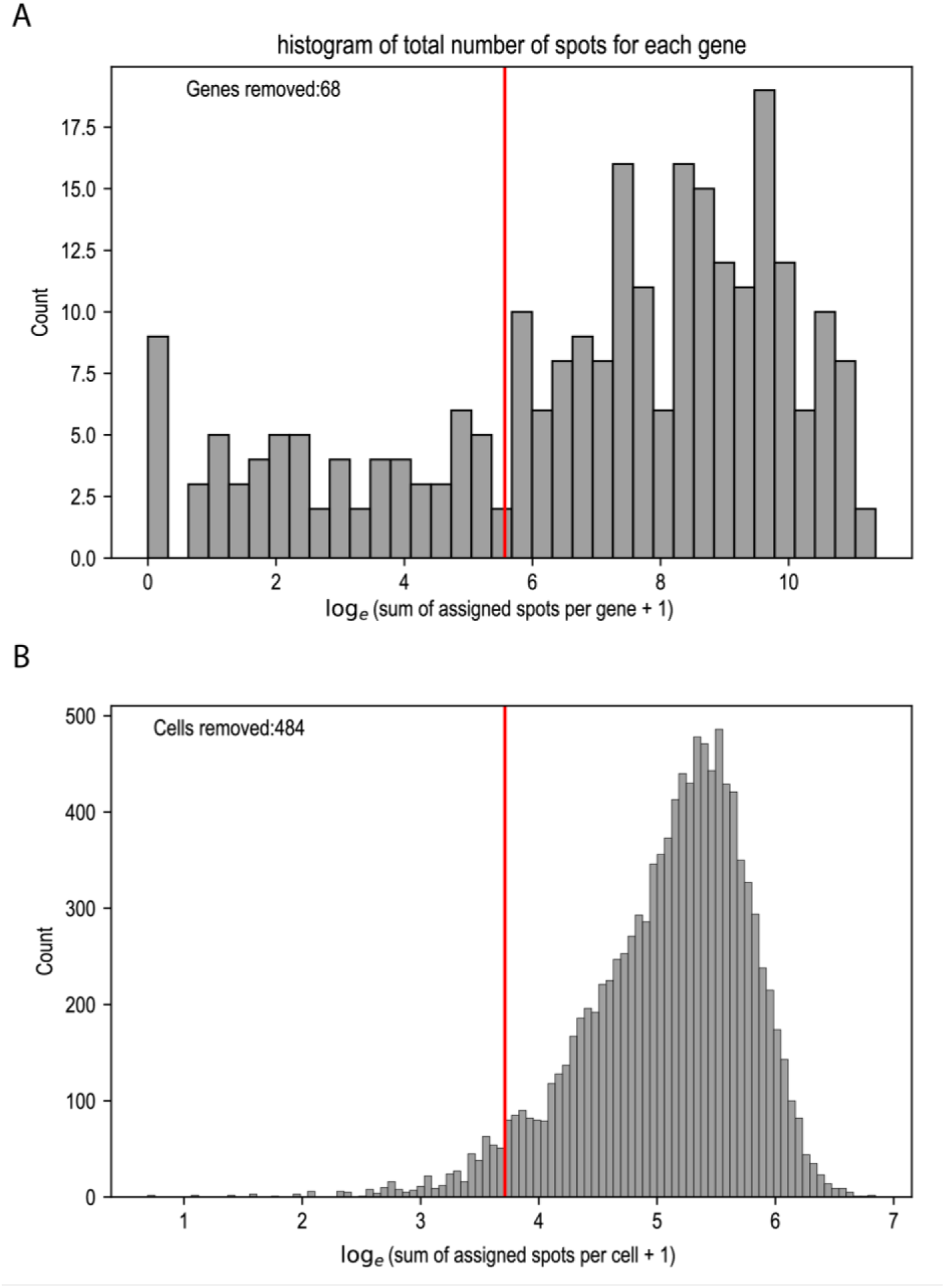
Gene expression thresholding and cell filtering. **(A)** A histogram showing the distribution of total spot counts for each gene across all cells on a logarithmic scale to accommodate the broad range of expression levels. Genes with fewer than 410 total spots are removed based on Otsu’s method, retaining 186 genes for subsequent analysis. **(B)** A histogram presenting the total number of spots assigned per cell, also on a logarithmic scale. Cells with spot counts below 40, approximately 484 cells (5% of the total), were excluded from further analysis.

**Fig. S21.**
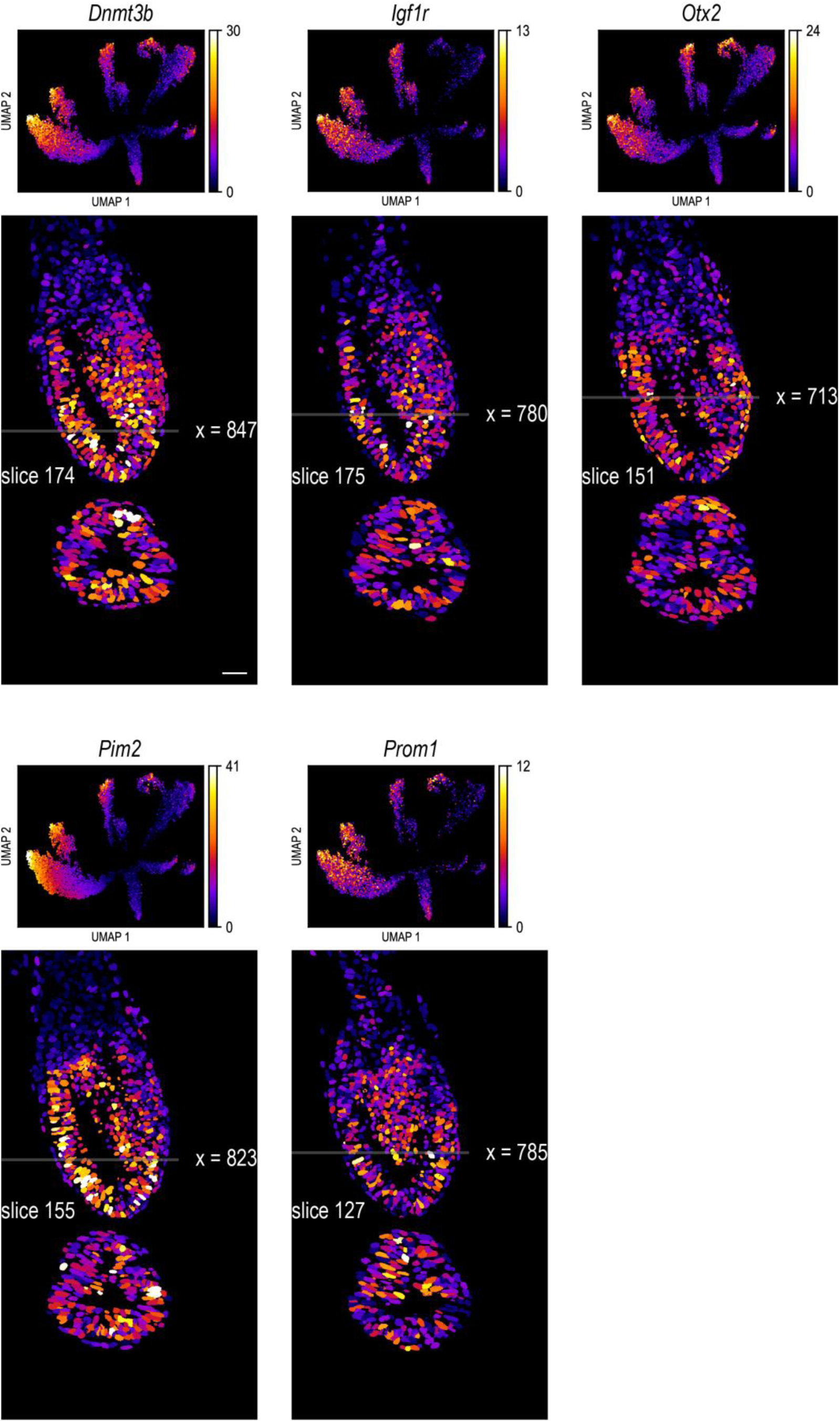
UMAP imputation and 3D spatial gene expression – cluster 1. Gene expression levels for five selected genes are color-coded in the upper panel, providing a visual representation of gene expression variations among clusters. The lower panel offers selected *xy* and *yz* views of the embryo, illustrating the spatial distribution of gene expression throughout the embryo. A unified color map adjacent to the UMAP panel aids in interpreting the gene expression levels. Scale bar: 50µm.

**Fig. S22.**
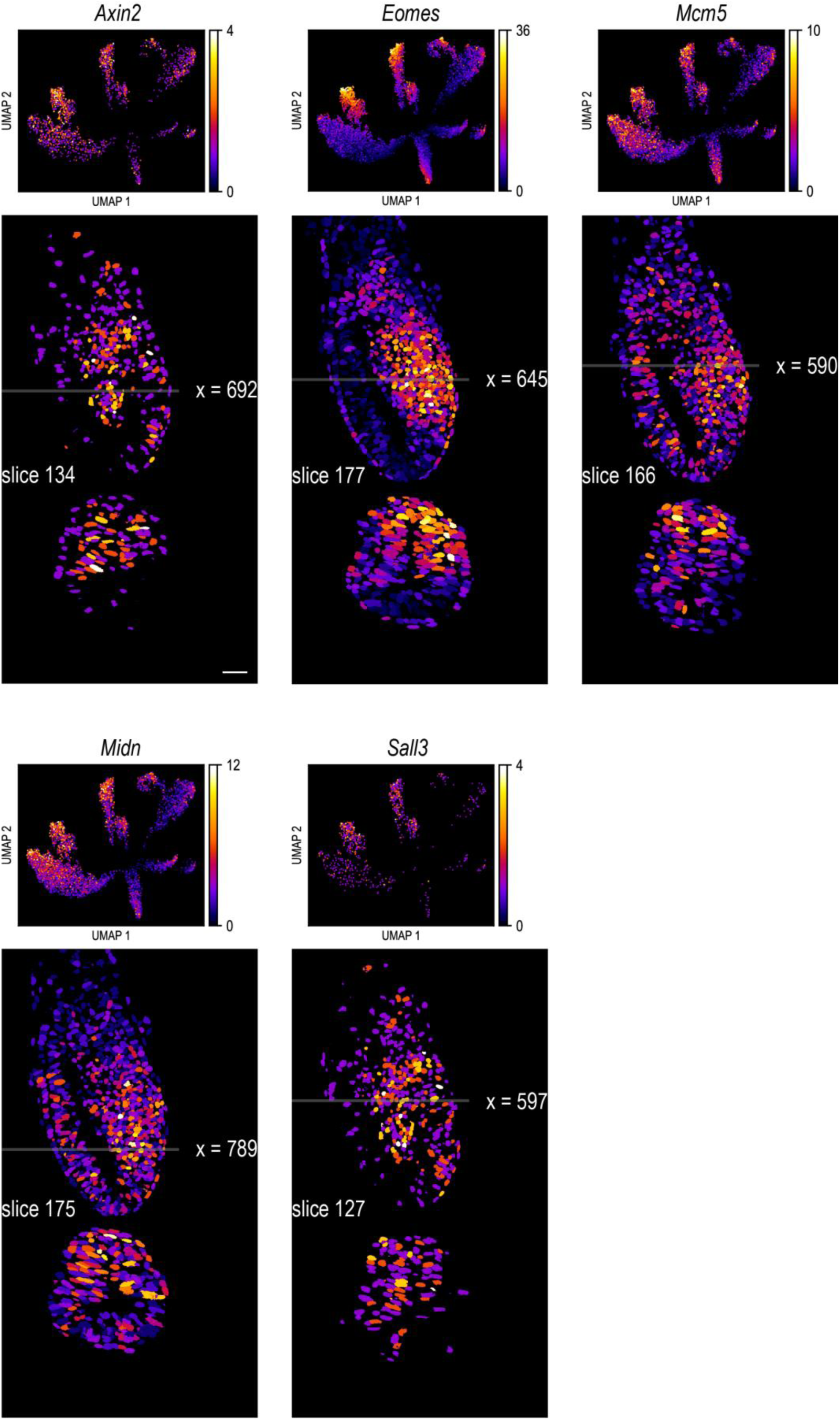
UMAP imputation and 3D spatial gene expression – cluster 2. Gene expression levels for five selected genes are color-coded in the upper panel, providing a visual representation of gene expression variations among clusters. The lower panel offers selected *xy* and *yz* views of the embryo, illustrating the spatial distribution of gene expression throughout the embryo. A unified color map adjacent to the UMAP panel aids in interpreting the gene expression levels. Scale bar: 50µm.

**Fig. S23.**
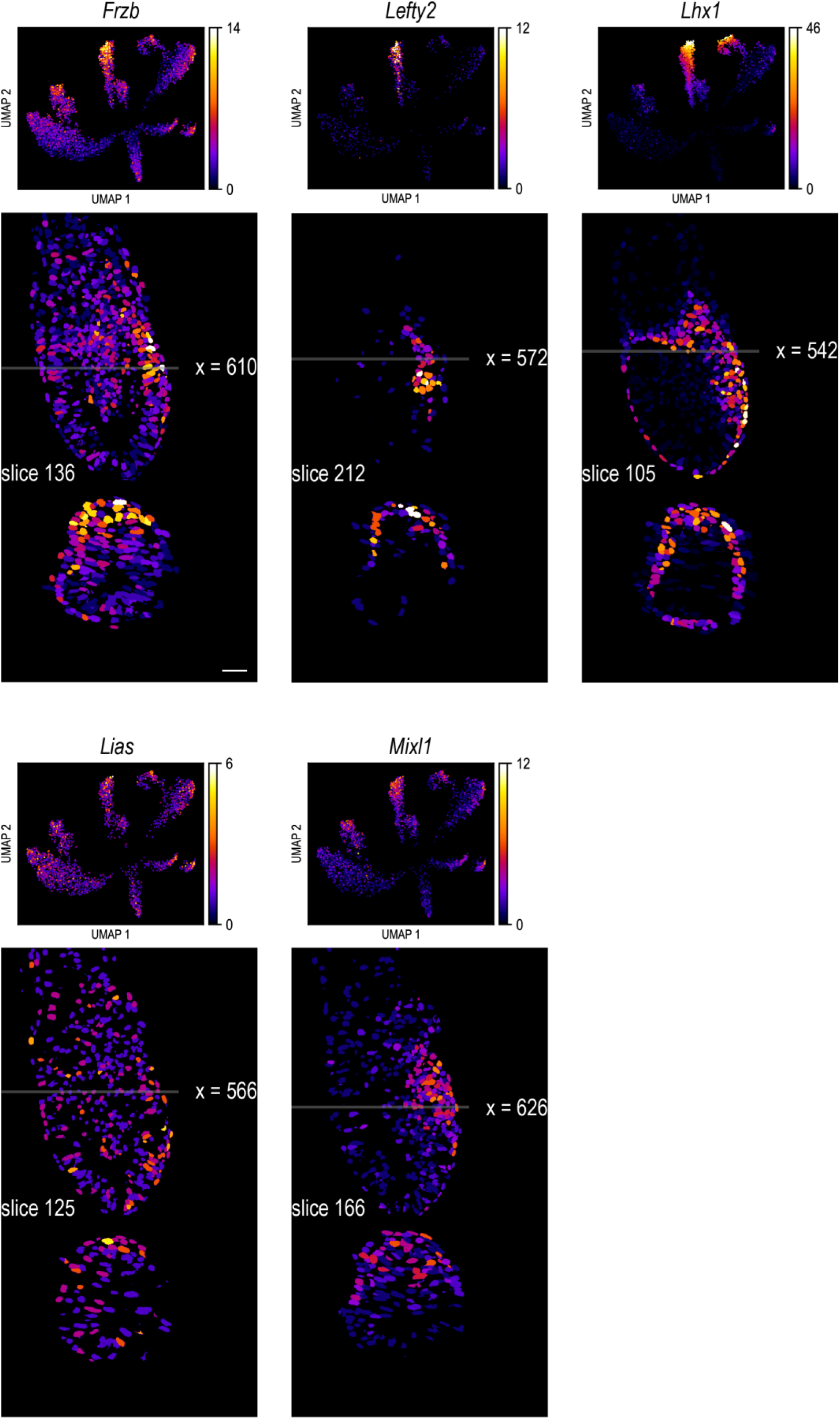
UMAP imputation and 3D spatial gene expression – cluster 3. Gene expression levels for five selected genes are color-coded in the upper panel, providing a visual representation of gene expression variations among clusters. The lower panel offers selected *xy* and *yz* views of the embryo, illustrating the spatial distribution of gene expression throughout the embryo. A unified color map adjacent to the UMAP panel aids in interpreting the gene expression levels. Scale bar: 50µm.

**Fig. S24.**
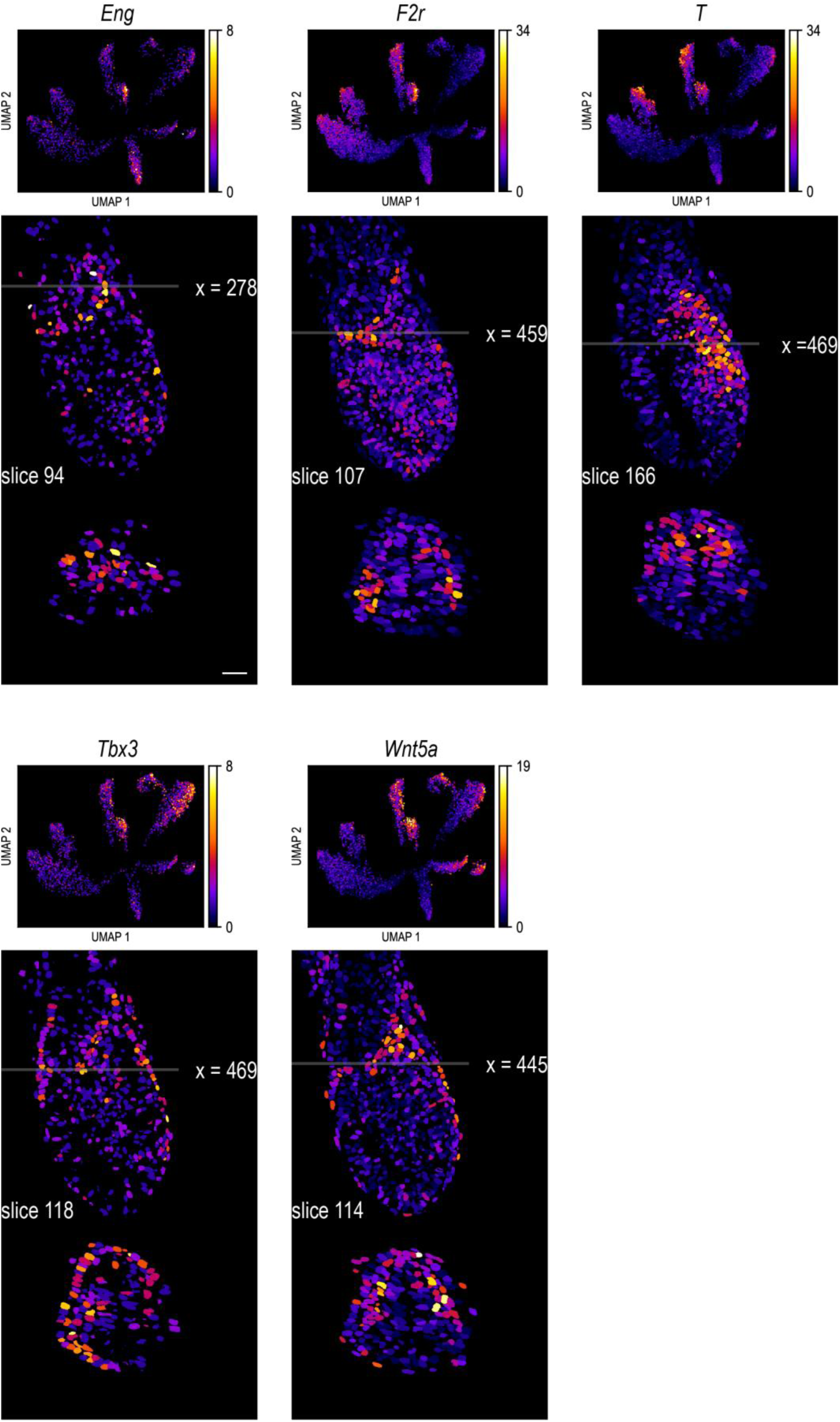
UMAP imputation and 3D spatial gene expression – cluster 4. Gene expression levels for five selected genes are color-coded in the upper panel, providing a visual representation of gene expression variations among clusters. The lower panel offers selected *xy* and *yz* views of the embryo, illustrating the spatial distribution of gene expression throughout the embryo. A unified color map adjacent to the UMAP panel aids in interpreting the gene expression levels. Scale bar: 50µm.

**Fig. S25.**
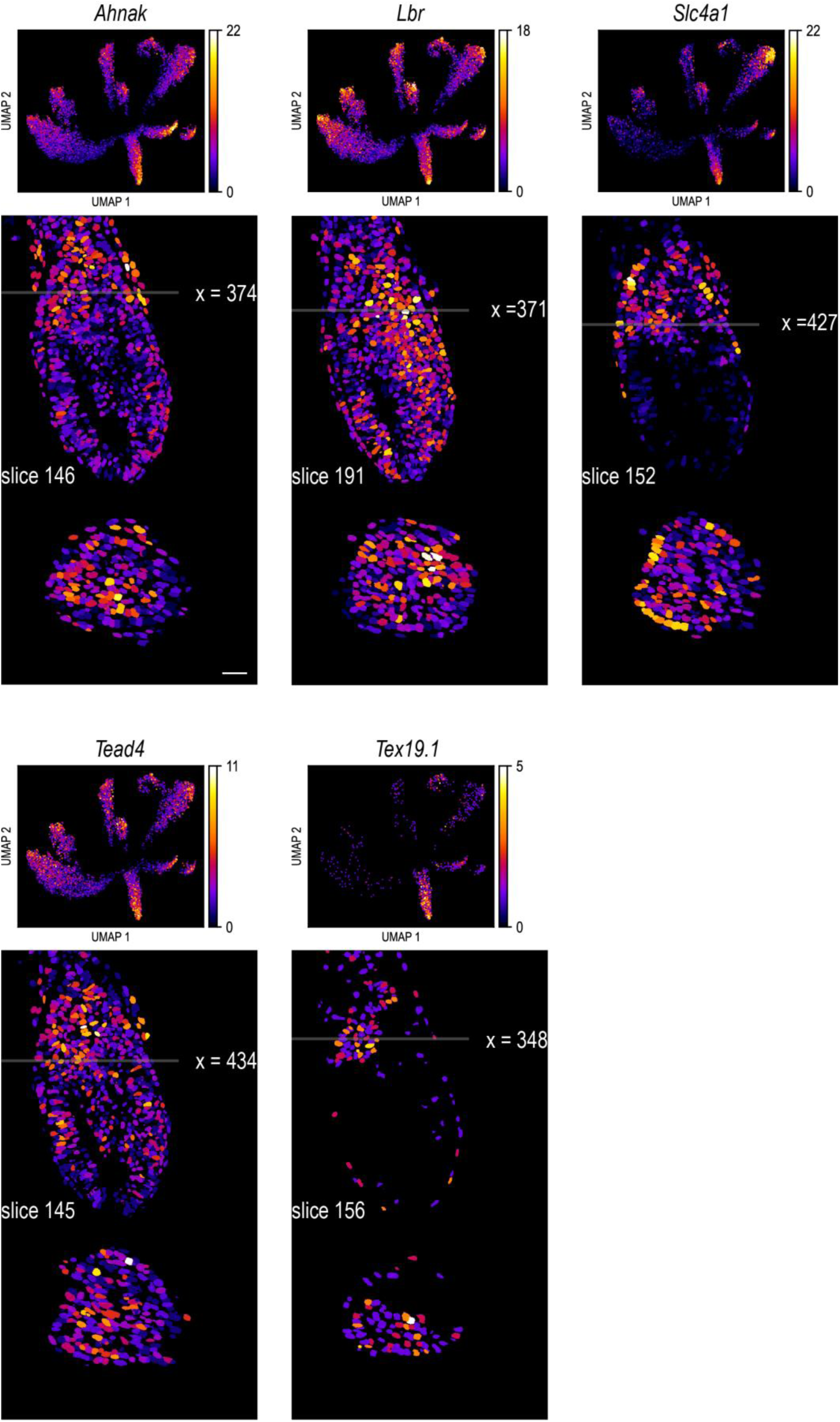
UMAP imputation and 3D spatial gene expression – cluster 5. Gene expression levels for five selected genes are color-coded in the upper panel, providing a visual representation of gene expression variations among clusters. The lower panel offers selected *xy* and *yz* views of the embryo, illustrating the spatial distribution of gene expression throughout the embryo. A unified color map adjacent to the UMAP panel aids in interpreting the gene expression levels. Scale bar: 50µm.

**Fig. S26.**
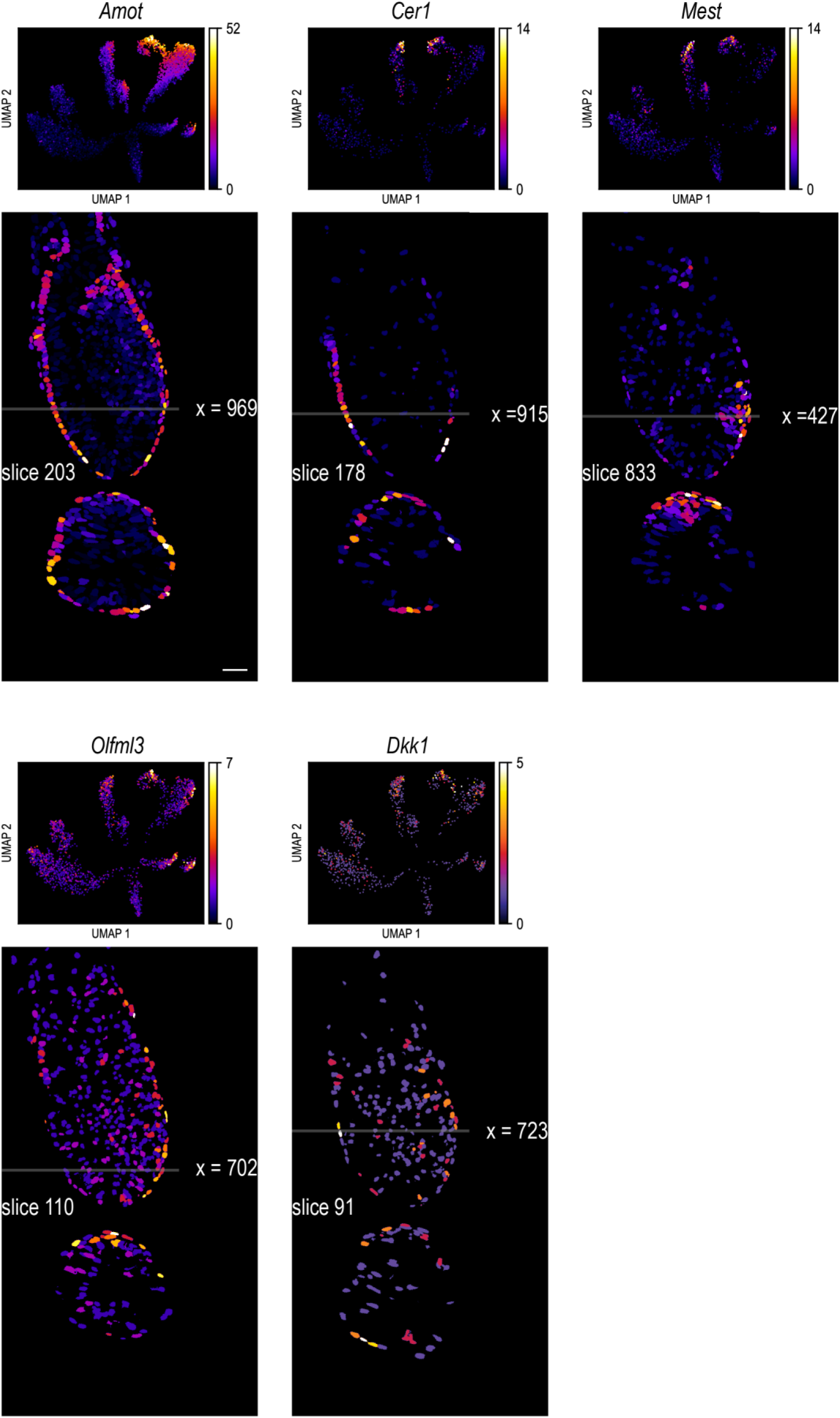
UMAP imputation and 3D spatial gene expression – cluster 6. Gene expression levels for five selected genes are color-coded in the upper panel, providing a visual representation of gene expression variations among clusters. The lower panel offers selected *xy* and *yz* views of the embryo, illustrating the spatial distribution of gene expression throughout the embryo. A unified color map adjacent to the UMAP panel aids in interpreting the gene expression levels. Scale bar: 50µm.

**Fig. S27.**
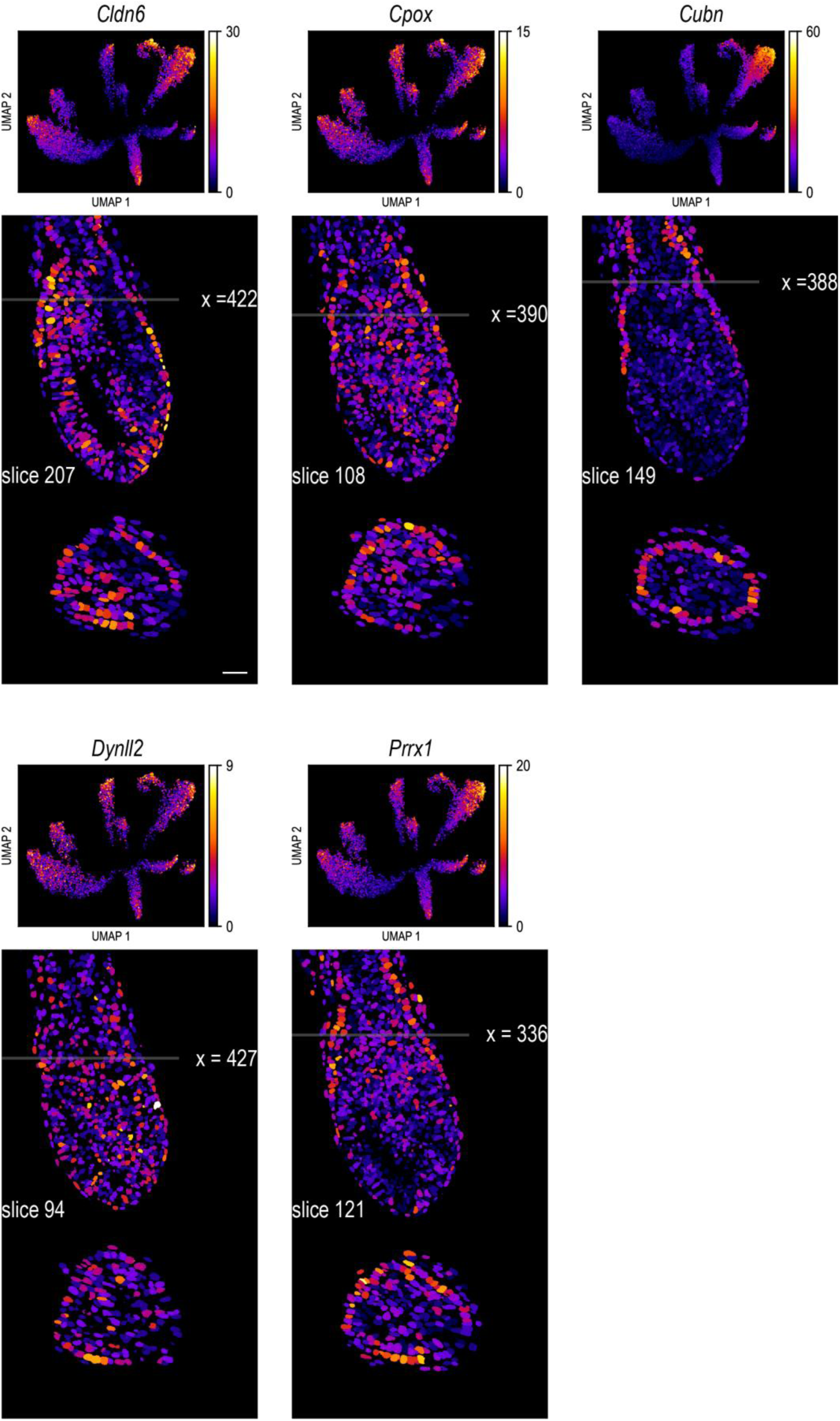
UMAP imputation and 3D spatial gene expression – cluster 7. Gene expression levels for five selected genes are color-coded in the upper panel, providing a visual representation of gene expression variations among clusters. The lower panel offers selected *xy* and *yz* views of the embryo, illustrating the spatial distribution of gene expression throughout the embryo. A unified color map adjacent to the UMAP panel aids in interpreting the gene expression levels. Scale bar: 50µm.

**Fig. S28.**
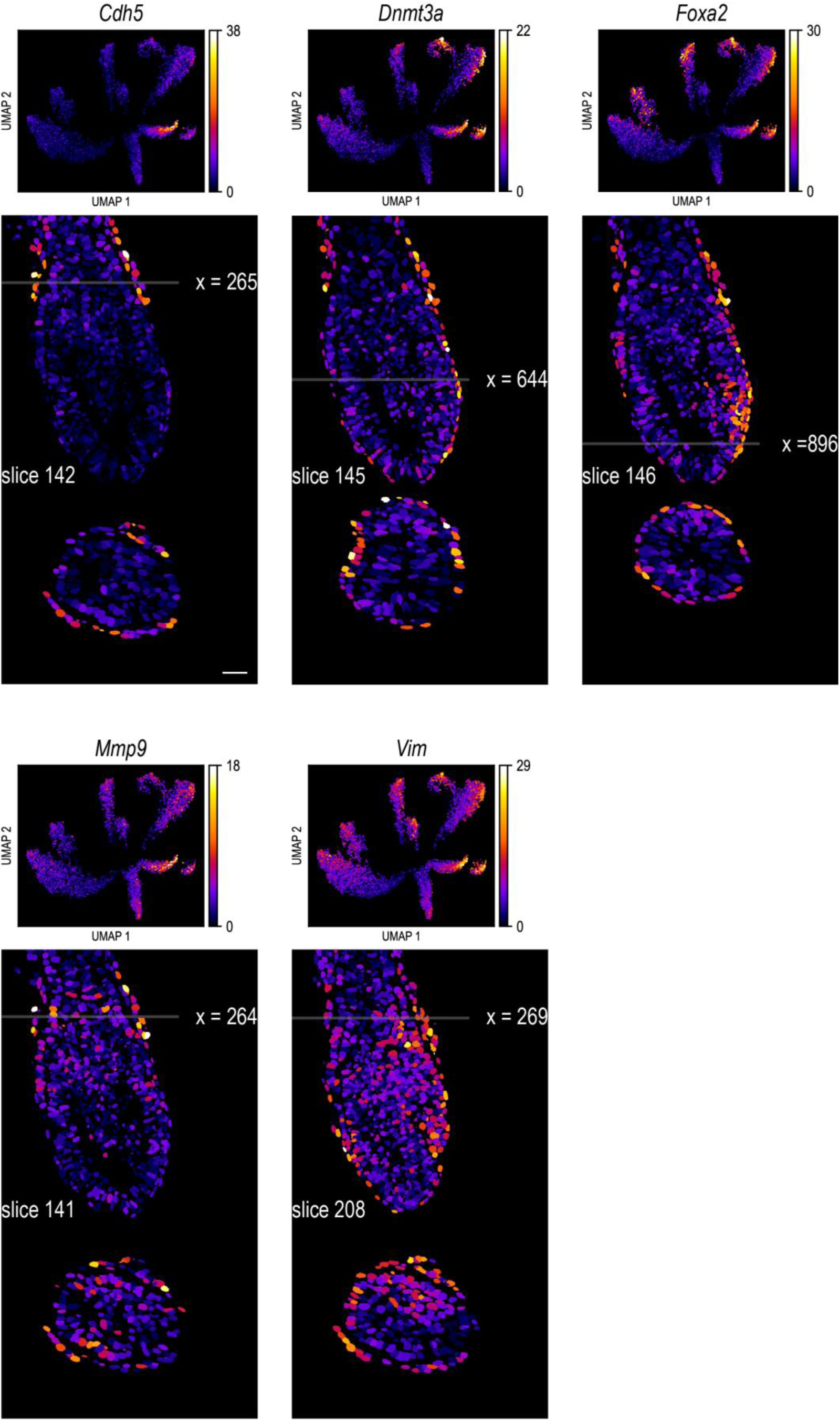
UMAP imputation and 3D spatial gene expression – cluster 8. Gene expression levels for five selected genes are color-coded in the upper panel, providing a visual representation of gene expression variations among clusters. The lower panel offers selected *xy* and *yz* views of the embryo, illustrating the spatial distribution of gene expression throughout the embryo. A unified color map adjacent to the UMAP panel aids in interpreting the gene expression levels. Scale bar: 50µm.

**Fig. S29.**
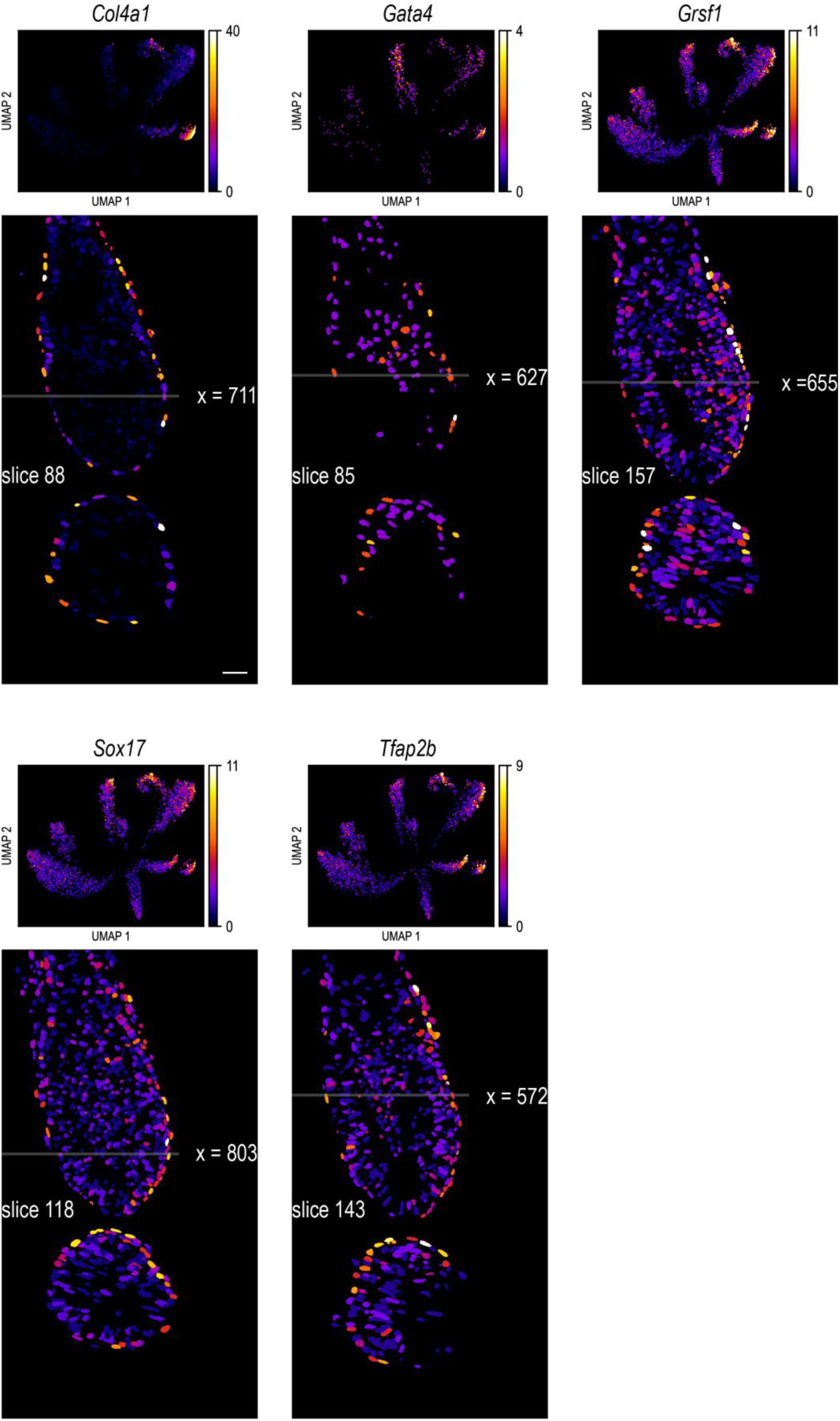
UMAP imputation and 3D spatial gene expression – cluster 9. Gene expression levels for five selected genes are color-coded in the upper panel, providing a visual representation of gene expression variations among clusters. The lower panel offers selected *xy* and *yz* views of the embryo, illustrating the spatial distribution of gene expression throughout the embryo. A unified color map adjacent to the UMAP panel aids in interpreting the gene expression levels. Scale bar: 50µm.

**Fig. S30.**
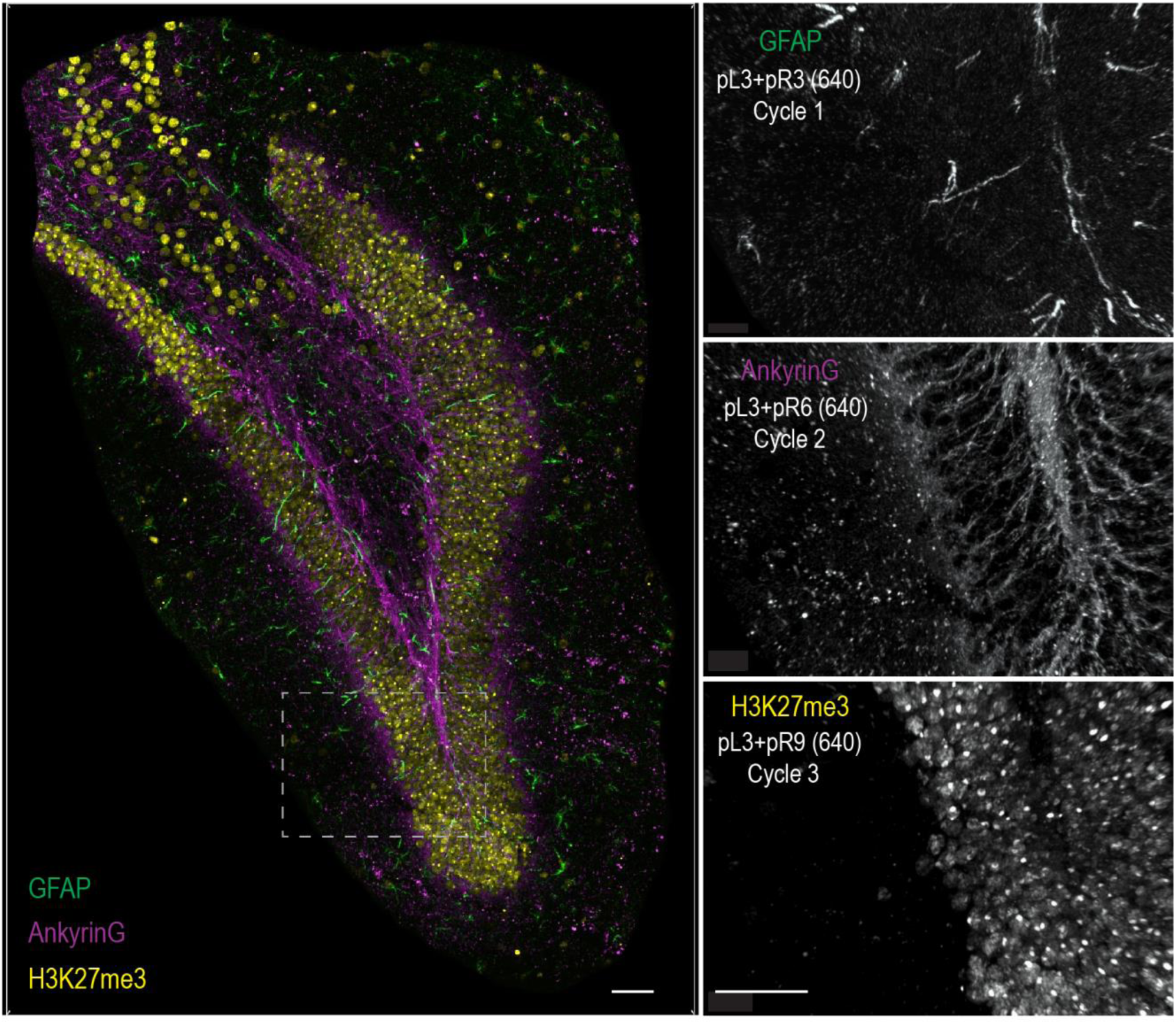
Demonstrating cycleHCR protein labeling specificity in tissue. This figure shows the capacity of cycleHCR to achieve protein labeling specificity within a complex tissue matrix, specifically in the dentate gyrus of a 40 µm thick hippocampal slice. By maintaining a constant left readout probe (pL3) and sequentially changing the right readout probes, cycleHCR shifts its target from the glial cell marker GFAP (pL3+pR3 at 640nm, Cycle 1) to the axonal initial segment marker ankyrin G (pL3+pR6 at 640nm, Cycle 2), and finally to the heterochromatin marker H3K27me3 (pL3+pR9 at 640nm, Cycle 3). The composite image is on the left panel and zoom-in cross-cycle views of the boxed region are displayed on the right. Scale bar: 50µm.

**Fig. S31.**
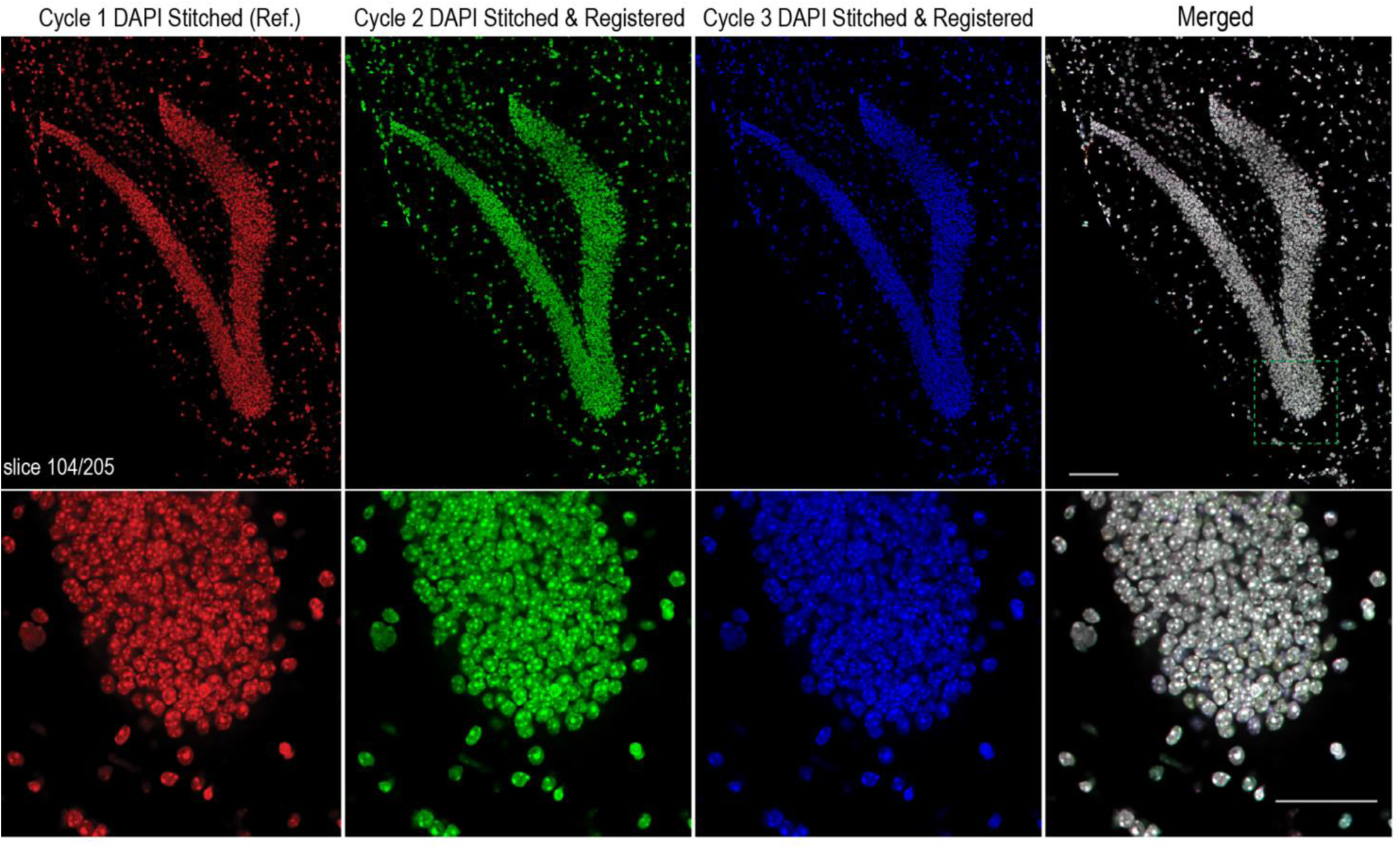
Cross-cycle registration validation for protein cycleHCR in mouse hippocampal slice. This figure illustrates the cross-cycle registration for protein cycleHCR imaging using a mouse hippocampal slice showed in Figure S31. The alignment of the reference DAPI image (depicted in red) with the DAPI images from subsequent cycles—Cycle 2 (shown in green) and Cycle 3 (in blue). A detailed examination within the boxed area in the upper panel underscores the precise alignment of DAPI signals across the three consecutive cycleHCR imaging rounds, confirming the reliability of the imaging and registration process over multiple cycles. Scale bar: 100µm (the upper panel); 50µm (the lower panel).

**Fig. S32.**
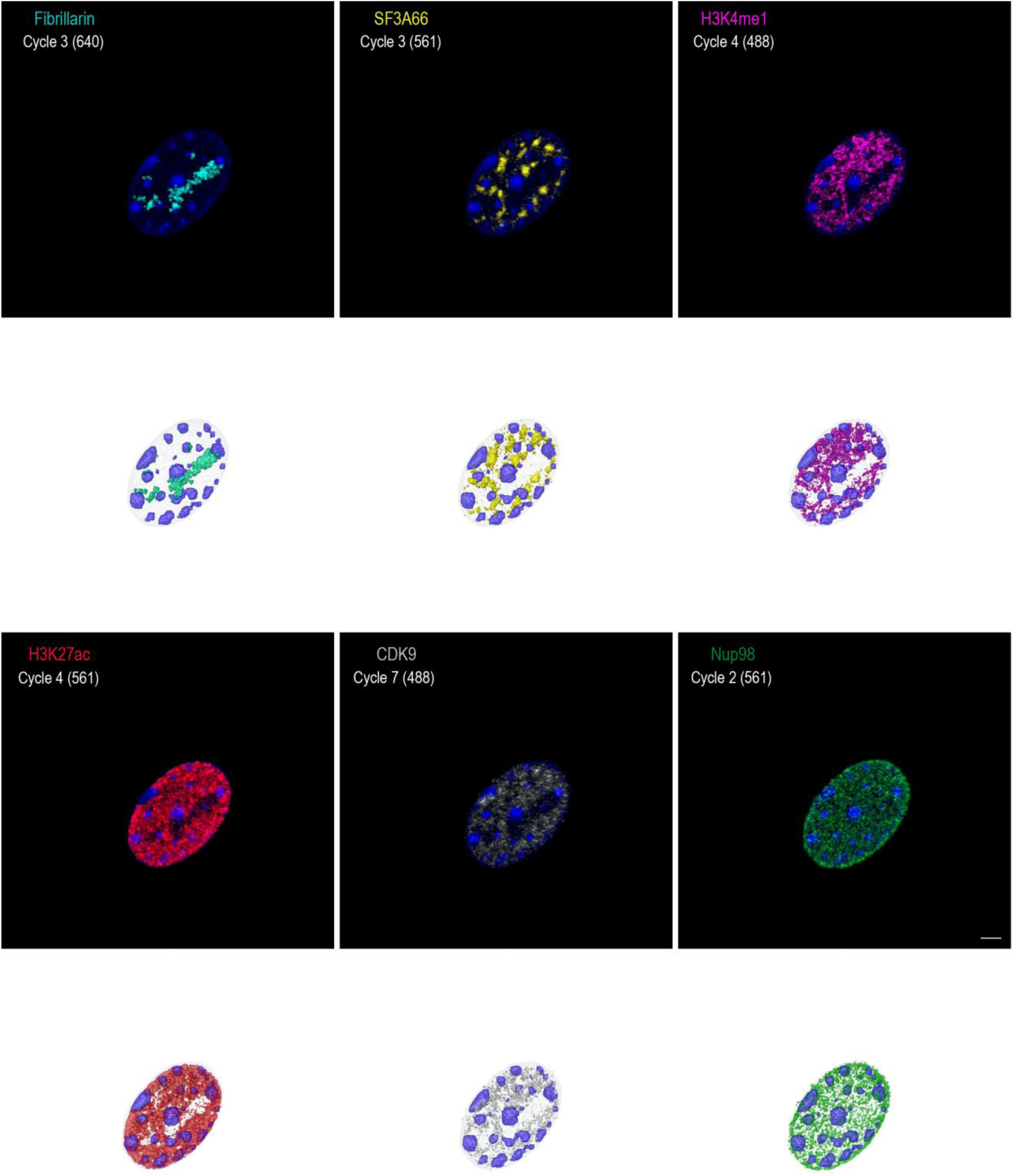
3D segmented models of subcellular structures – panel 1. Protein cycleHCR images of specific protein targets within a mouse embryonic fibroblast after ∼4X expansion are presented by Imaris Maximum Intensity Projection (MIP) volume rendering on the top. Accompanying each fluorescent image, 3D segmentation masks are displayed in mesh format, rendered by ORS Dragonfly. The proteins visualized include Fibrillarin (marking nucleolar fibrillar regions in the nucleolus), SF3A66 (highlighting nuclear speckles), H3K4me1 (associated with enhancers), H3K27ac (indicating active chromatin regions), CDK9 (involved in Pol II elongation), and Nup98 (marking nuclear pores). Scale bar: 10µm.

**Fig. S33.**
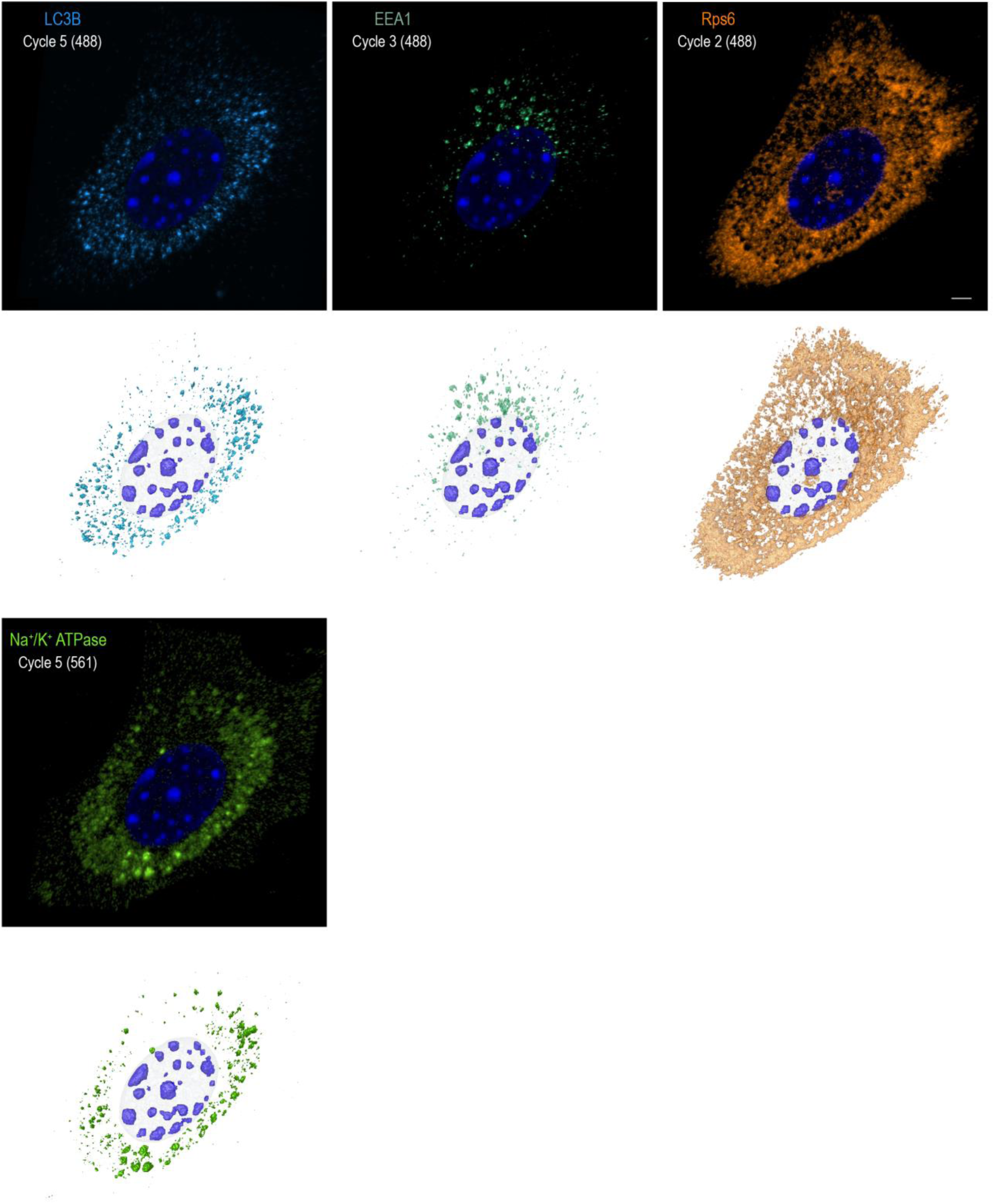
3D segmented models of subcellular structures – Panel 2. This figure continues the display of subcellular architectures in an expanded mouse embryonic fibroblast obtained by protein cycleHCR, including LC3B (autophagosomes), EEA1 (early endosomes), Rps6 (ribosomal subunit), and Na^+^/K^+^ ATPase. The distribution of each protein within the mouse embryonic fibroblast is visualized through Imaris MIP volume rendering, with 3D segmentation masks in mesh format rendered by ORS Dragonfly displayed below. Scale bar: 10µm.

**Fig. S34.**
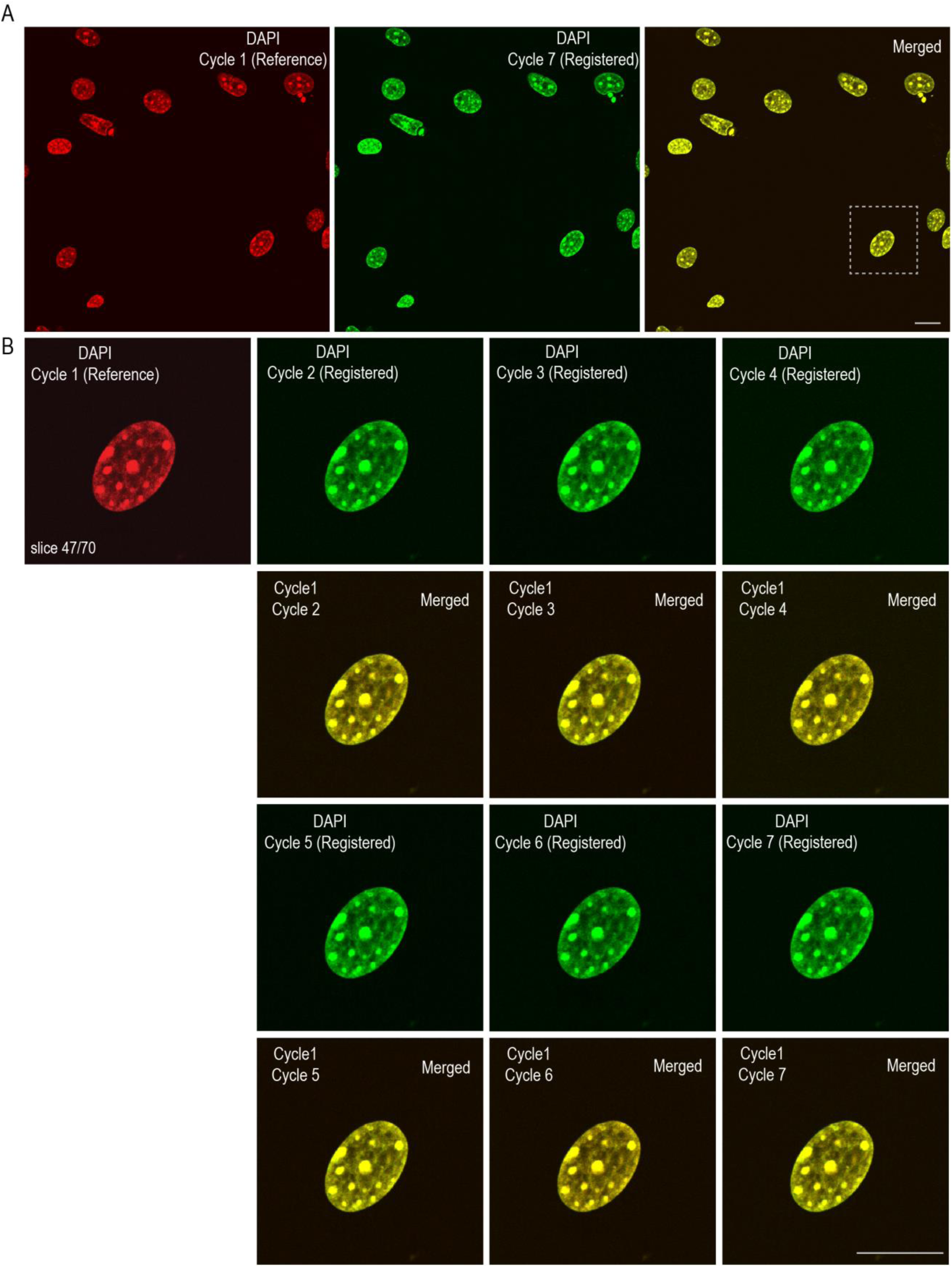
Verifying cross-round registration accuracy in protein cycleHCR imaging of mouse embryonic fibroblasts. **(A)** The precision of cross-cycle registration in protein cycleHCR imaging applied to ∼4X expanded mouse embryonic fibroblasts associated with Fig. 4, Fig. S32 and Fig. S33. The alignment of the reference DAPI image (depicted in red Cycle 1) with the DAPI images from last cycle — Cycle 7 (shown in green). **(B)** A detailed examination of the cell within the boxed area in (A): The alignment of the reference DAPI image from Cycle 1 (rendered in red) with DAPI images from subsequent cycles—Cycles 2 through 7 (each displayed in green). The composite image by fusing the reference image with the registered image are showed below, offering a visual assessment of the accuracy of cross-round registration. Scale bars: 50µm.

**Fig. S35.**
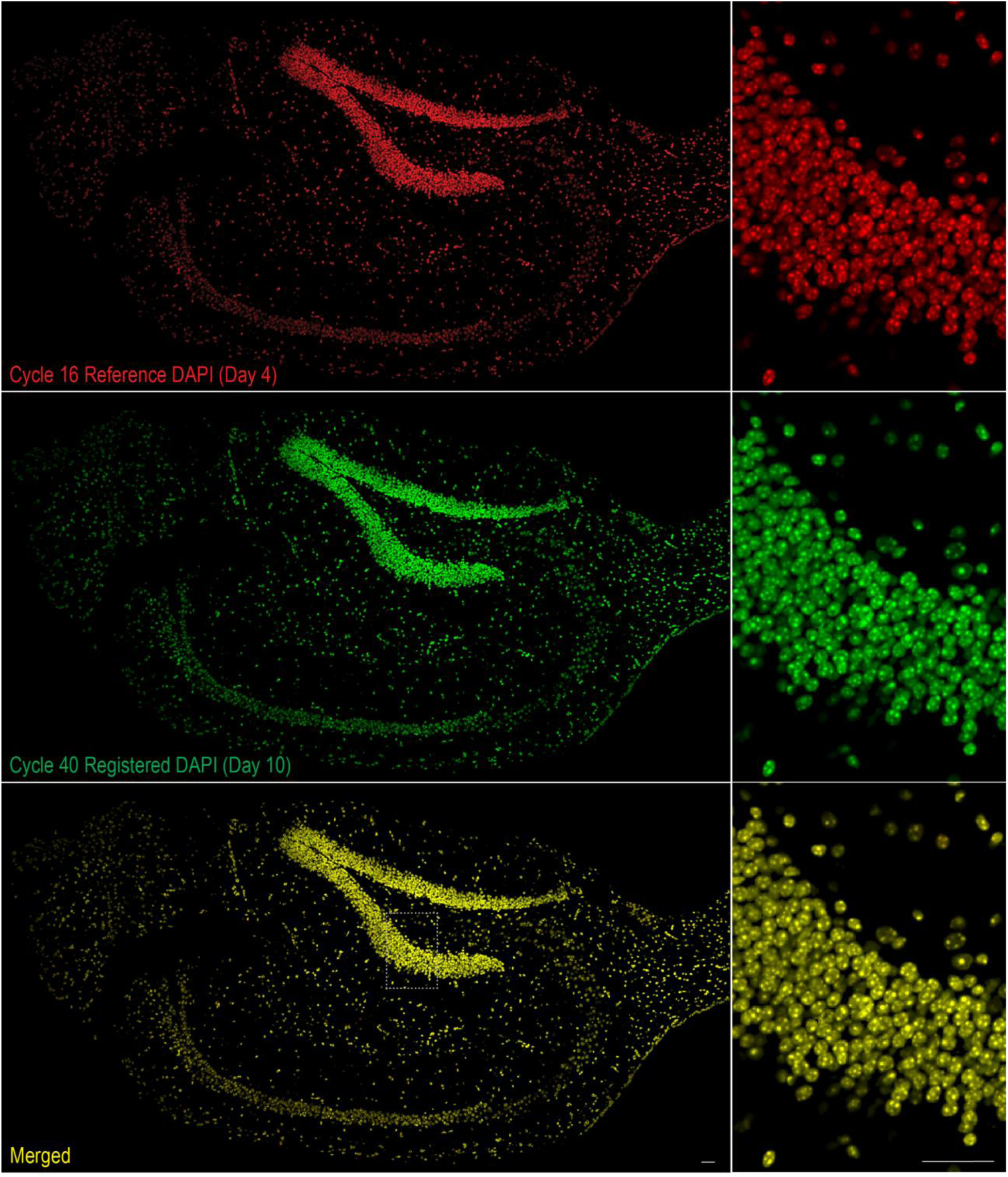
Verifying cross-round registration accuracy in joint cycleHCR protein and RNA imaging of hippocampal slice. This figure demonstrates the precision of cross-cycle registration in joint cycleHCR protein and RNA imaging of the hippocampal slice associated with Fig. 5. The alignment of the reference DAPI image (depicted in red for Cycle 15) with the DAPI images from the last RNA cycle (Cycle 40, shown in green) is illustrated. The composite image, created by fusing the reference image with the registered image, is shown below, providing a visual assessment of the accuracy of cross-round registration. Zoomed-in views of the boxed region are displayed on the left. Scale bars: 50µm.

**Fig. S36.**
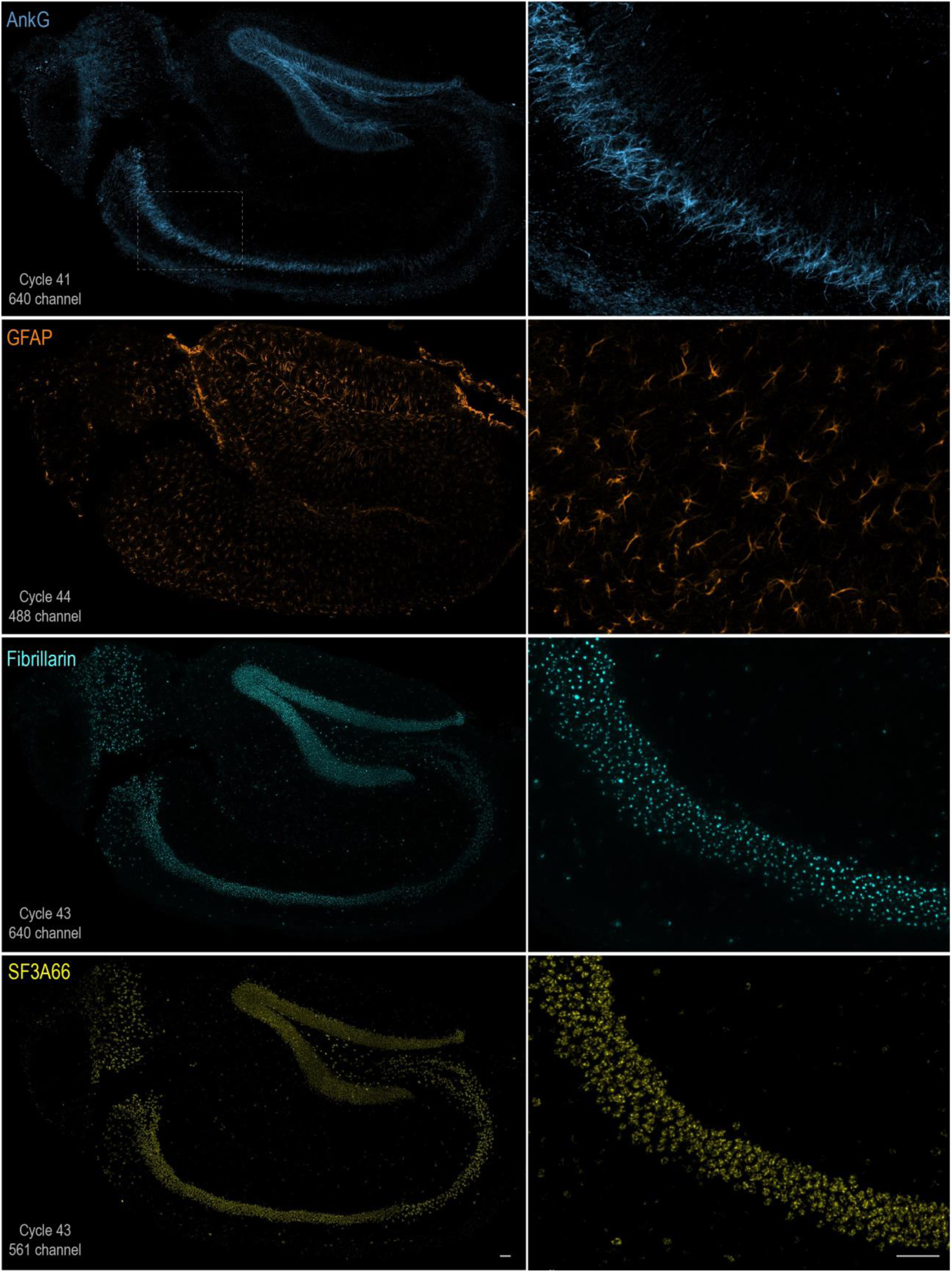
Protein cycleHCR in hippocampal slice – Panel 1. This panel presents stitched and registered images from protein cycleHCR targeting the axon initial segment marker AnkG, astrocyte marker GFAP, nucleolar fibrillar center marker Fibrillarin, and nuclear speckle marker SF3A66 in the hippocampal slice. Zoomed-in images of the boxed region are provided on the right panel. Scale bars: 50µm.

**Fig. S37.**
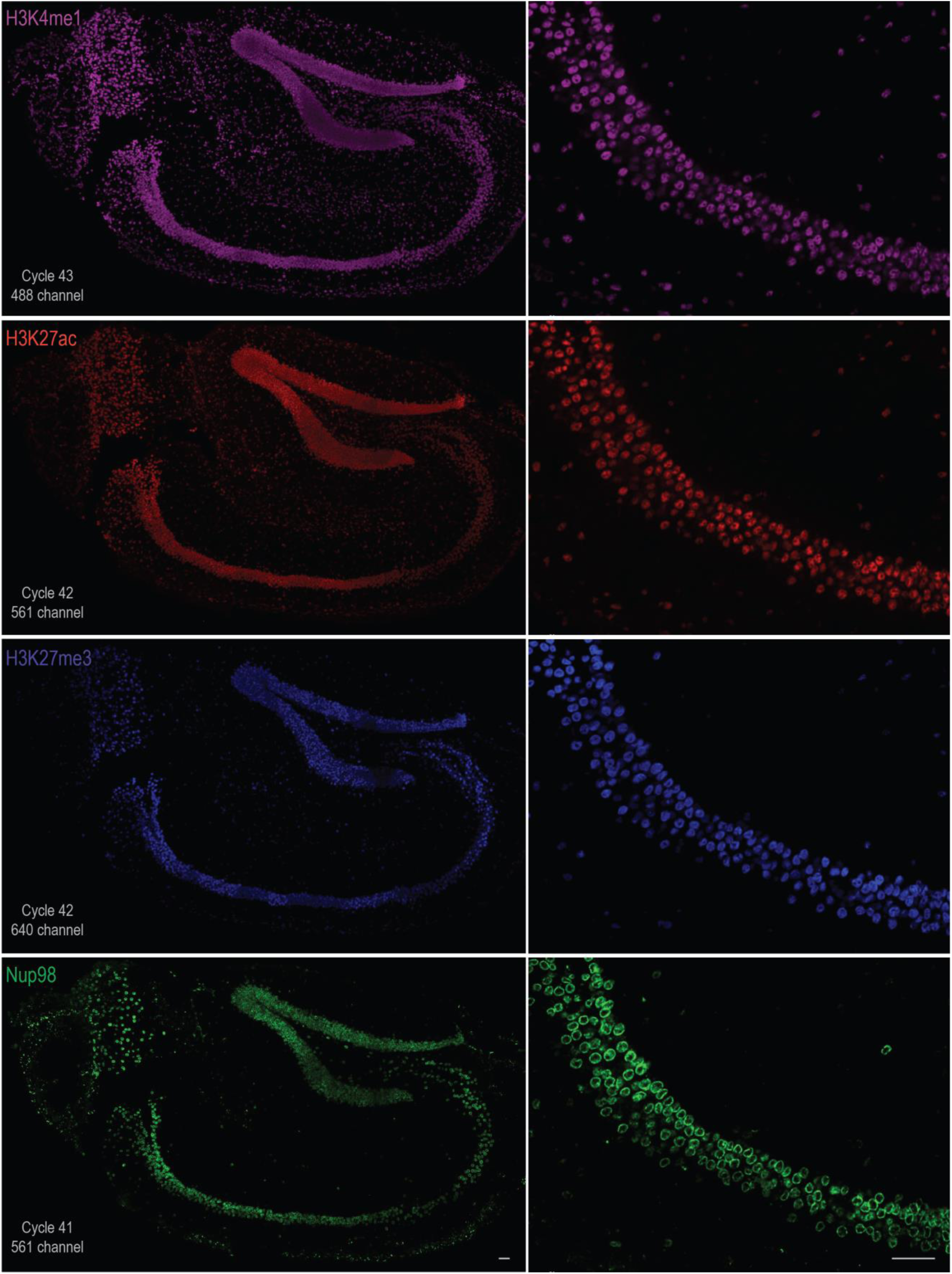
Protein cycleHCR in hippocampal slice – Panel 2. This panel displays stitched and registered images from protein cycleHCR targeting the active enhancer H3K4me1, active chromatin H3K27ac, inactive chromatin H3K27me3, and nuclear pores Nup98 in the hippocampal slice. Zoomed-in images of the boxed region showed in Fig. S36 are provided on the right panel. Scale bars: 50µm.

**Fig. S38.**
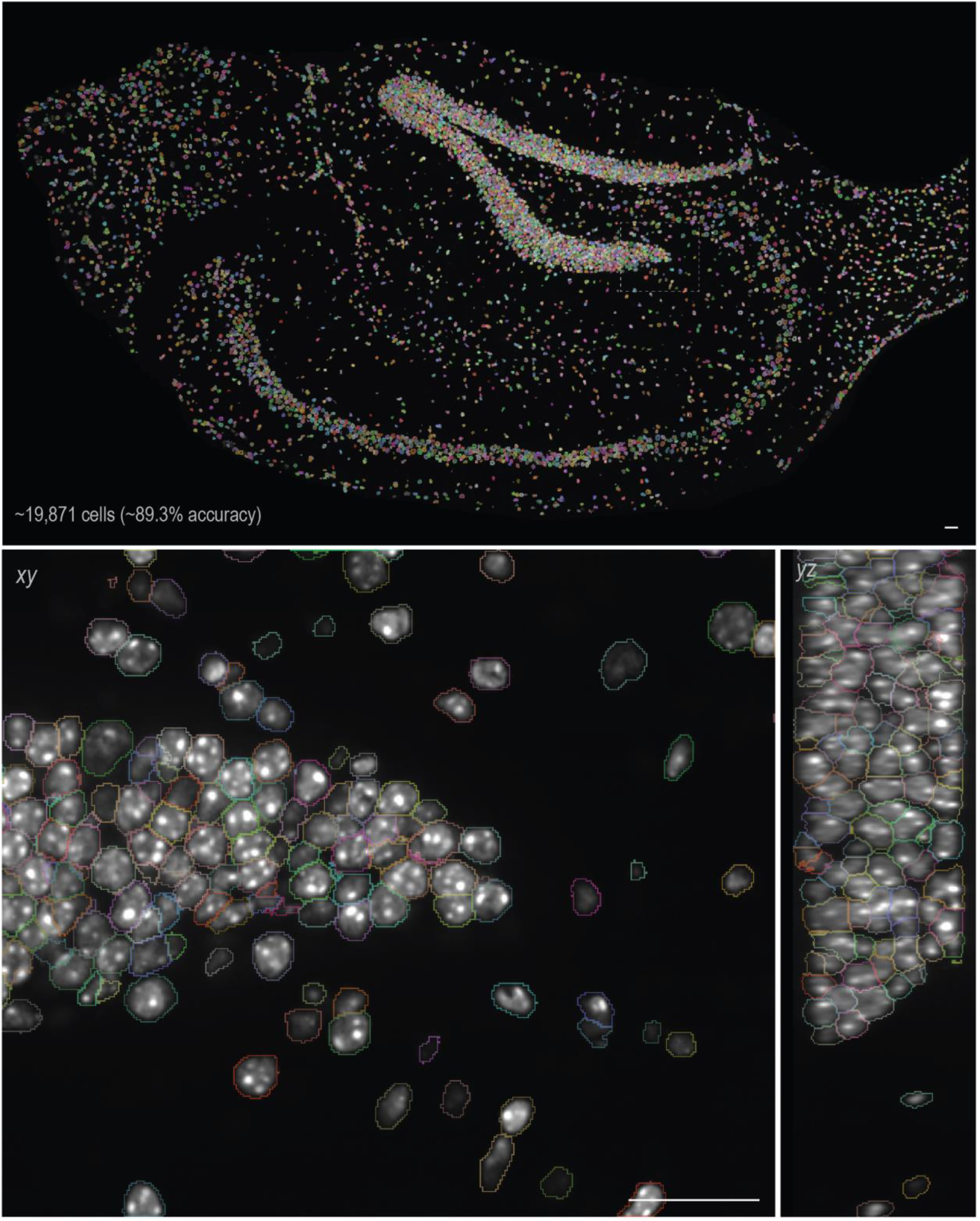
3D segmentation of hippocampal slice for joint protein and RNA cycleHCR imaging. The upper panel displays a slice view of segmented cells within the hippocampal slice by Cellpose, utilizing three-view segmentation of human-in-the-loop custom-trained models trained on orthogonal views (*xy* and *yz*). Contours delineate DAPI-stained nuclei. The lower two panels show the *xy* and *yz* views of the boxed region in the upper panel. Images were rendered using ORS Dragonfly. Scale bars: 50µm.

**Fig. S39.**
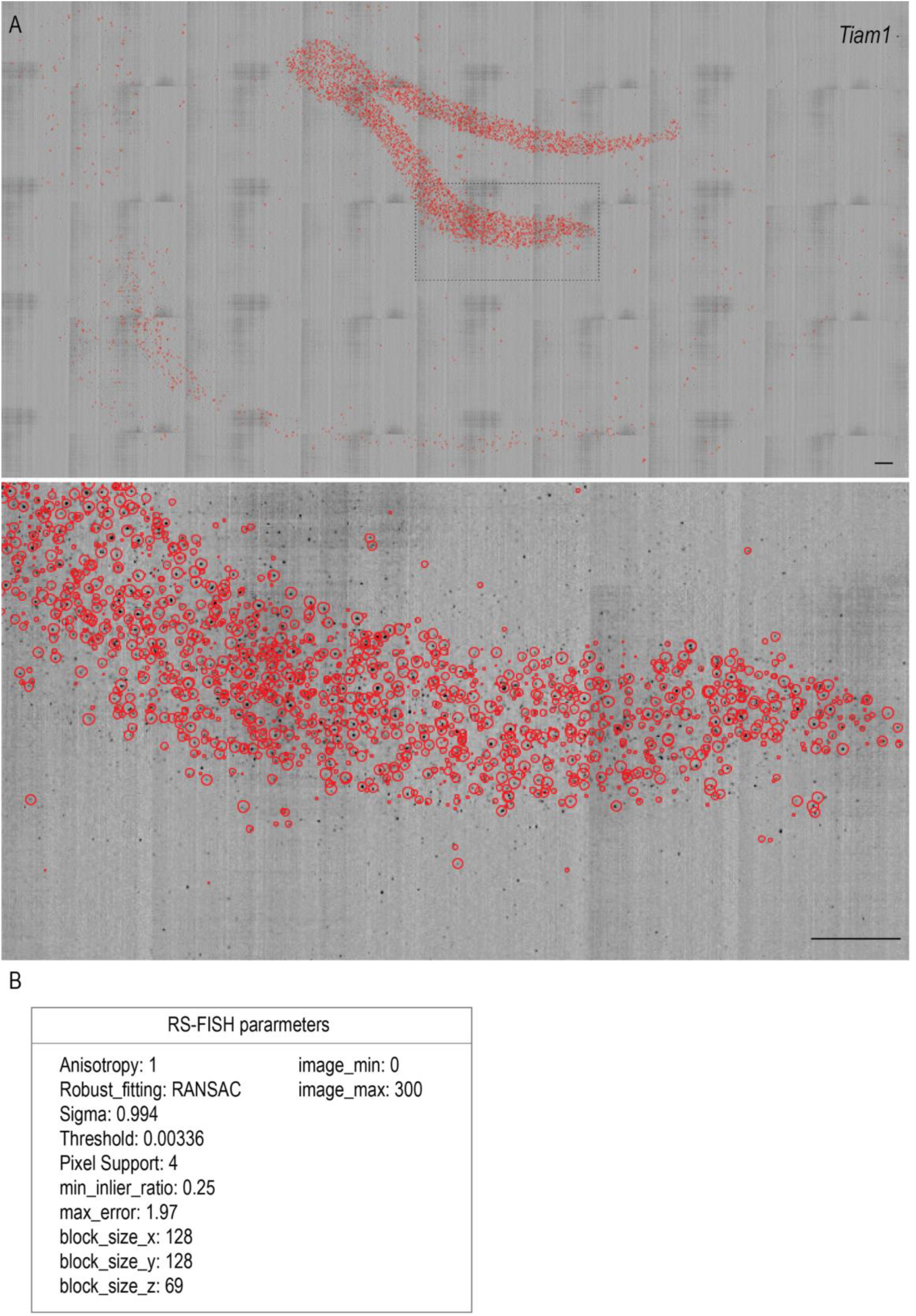
Single-molecule localization for joint protein and RNA cycleHCR imaging of hippocampal slice. **(A)** Single-molecule localization in imaging mRNA transcripts of the *Tiam1* gene in the hippocampal slice. Red circles indicate the positions of localized single molecules detected by RS-FISH, overlaid on an inverted raw image. A zoomed-in view of the boxed region is shown below. Scale bar: 50µm. **(B)** Single-molecule localization parameters with RS-FISH for localizing single molecules. These parameters were carefully selected to ensure minimal false positive detections and were maintained consistent for processing data from all cycles. Scale bars: 50µm.

**Fig. S40.**
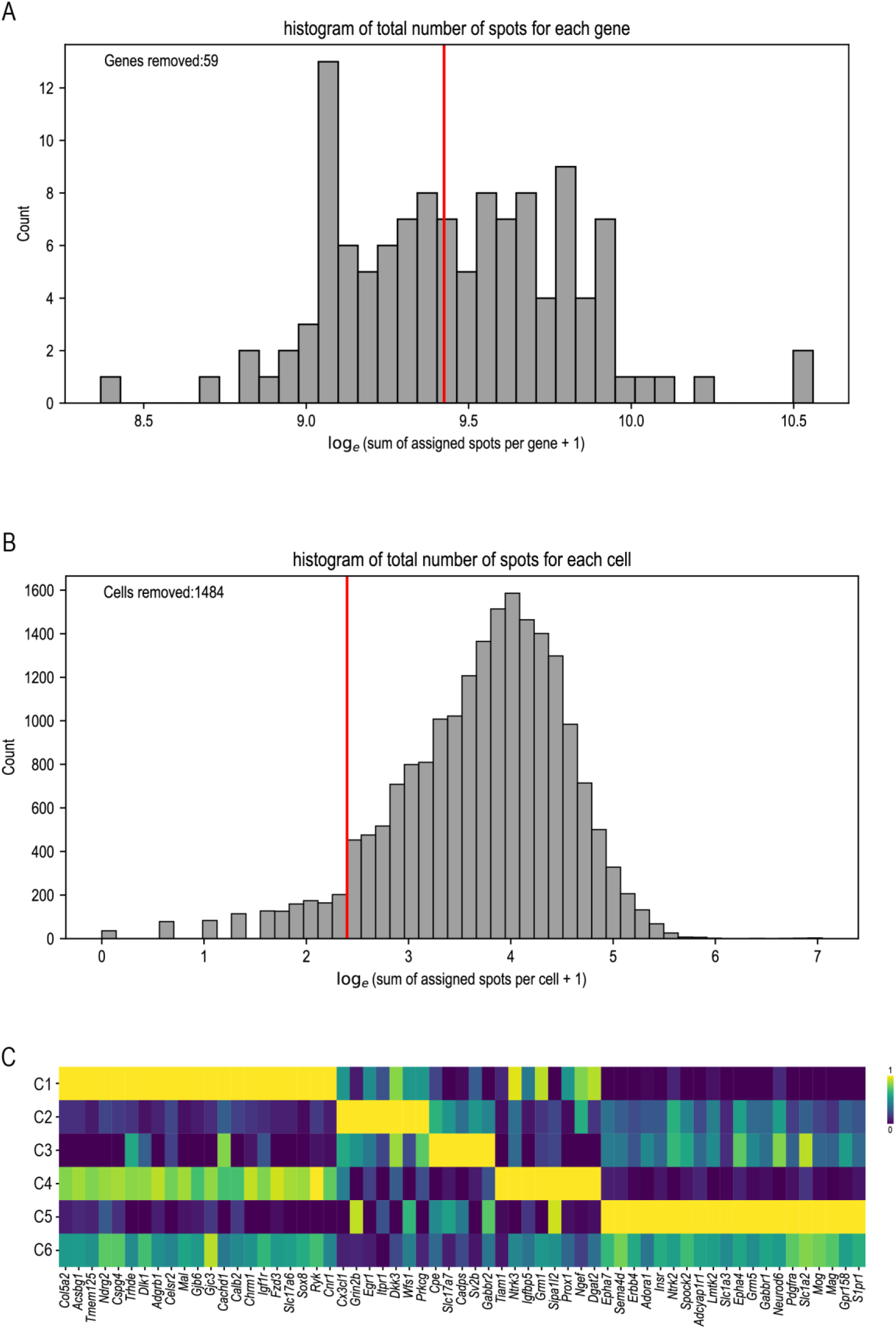
Gene and cell filtering prior to UMAP analysis. **(A)** Histogram displaying the distribution of total spot counts for each gene across all cells on a logarithmic scale. Genes with fewer than 12,386 spots were removed based on Li’s method, retaining 61 genes for subsequent analysis. **(B)** Histogram illustrating the total number of spots for each cell across the retained 61 genes on a logarithmic scale. Cells with spot counts less than 10, approximately 1,484 (7.5% of total), were excluded from further analysis. **(C)** Heatmap presenting the min-max scaled mean expression per cluster for each gene used in the UMAP analysis.

**Fig. S41.**
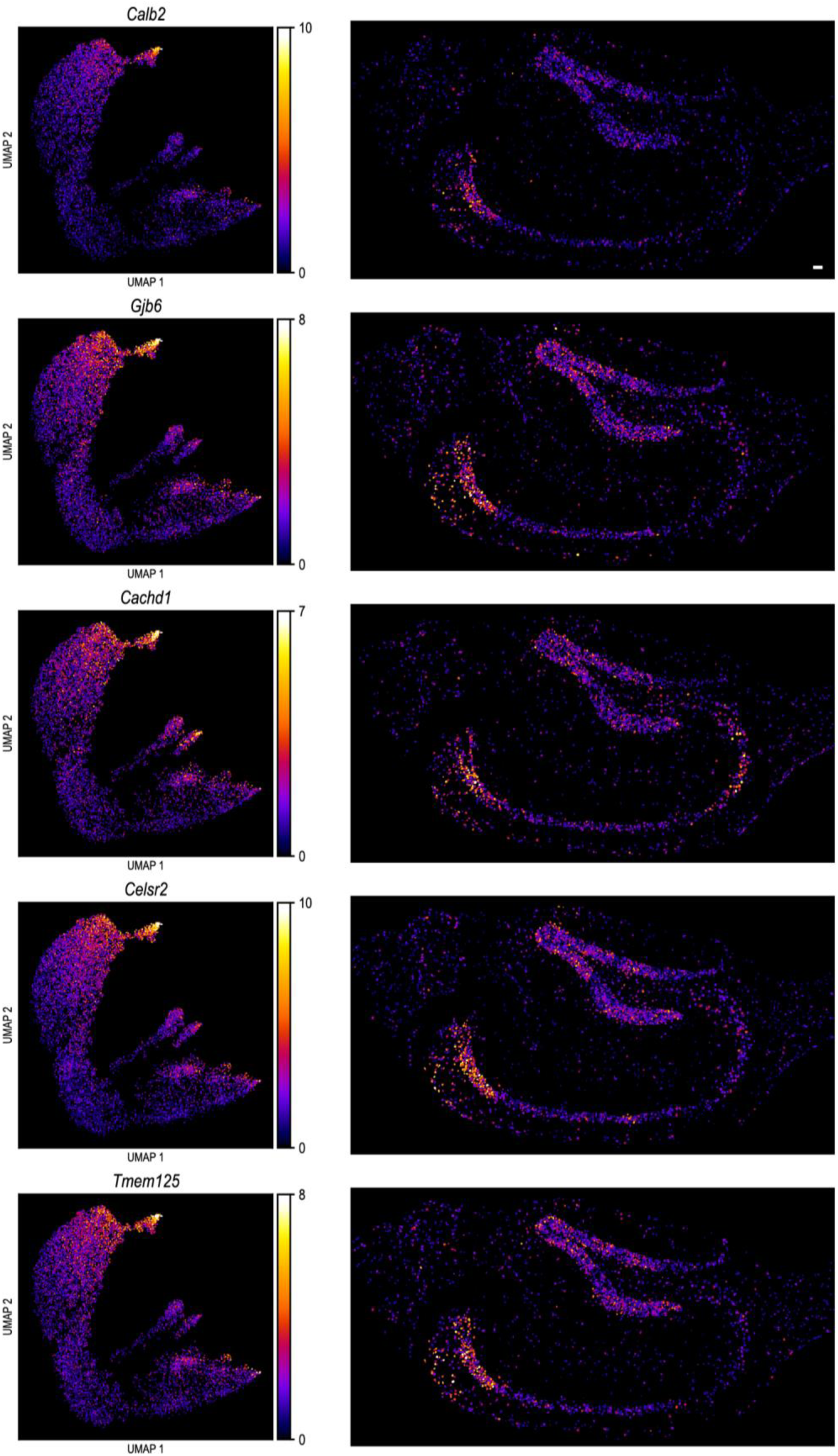
UMAP imputation of genes for hippocampal cluster 1. Gene expression levels for five selected genes are color-coded in the upper panel, providing a visual representation of gene expression variations among clusters. The left panel offers selected xy views of the hippocampal slice, illustrating the spatial distribution of gene expression. A unified color map adjacent to the UMAP panel aids in interpreting the gene expression levels. Scale bar: 50µm.

**Fig. S42.**
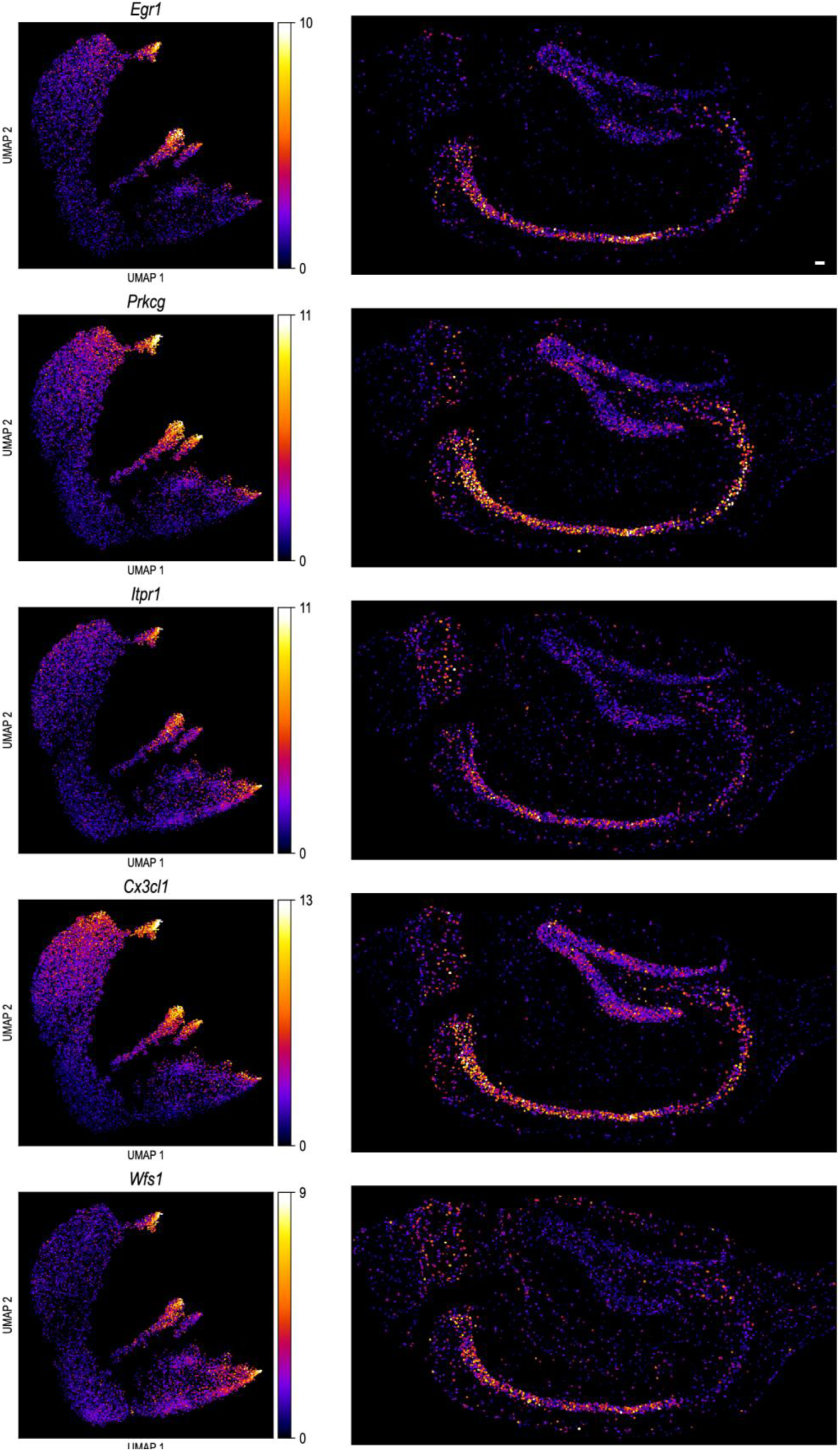
UMAP imputation of genes for hippocampal cluster 2. Gene expression levels for five selected genes are color-coded in the upper panel, providing a visual representation of gene expression variations among clusters. The left panel offers selected xy views of the hippocampal slice, illustrating the spatial distribution of gene expression. A unified color map adjacent to the UMAP panel aids in interpreting the gene expression levels. Scale bar: 50µm.

**Fig. S43.**
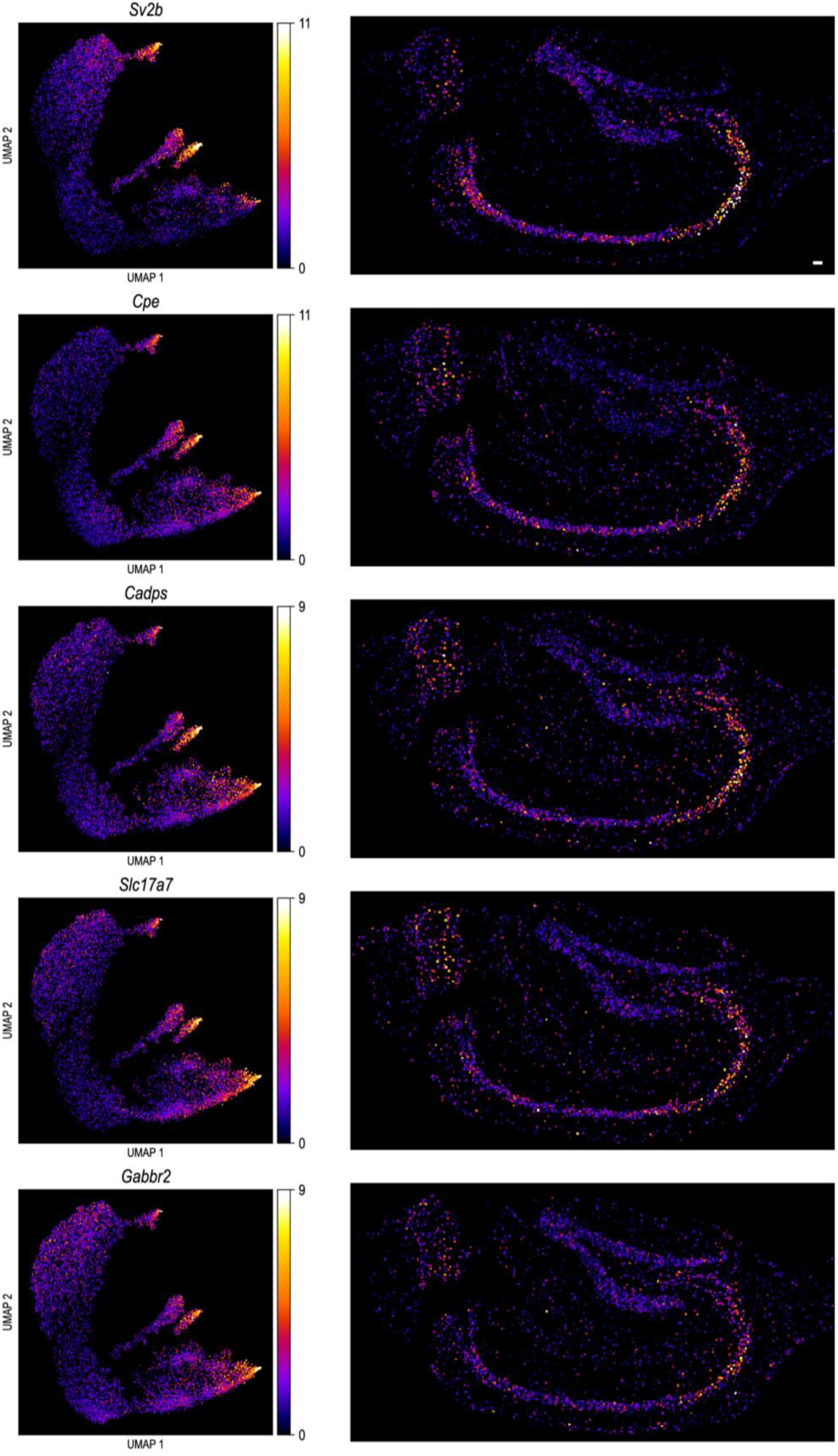
UMAP imputation of genes for hippocampal cluster 3. Gene expression levels for five selected genes are color-coded in the upper panel, providing a visual representation of gene expression variations among clusters. The left panel offers selected xy views of the hippocampal slice, illustrating the spatial distribution of gene expression. A unified color map adjacent to the UMAP panel aids in interpreting the gene expression levels. Scale bar: 50µm.

**Fig. S44.**
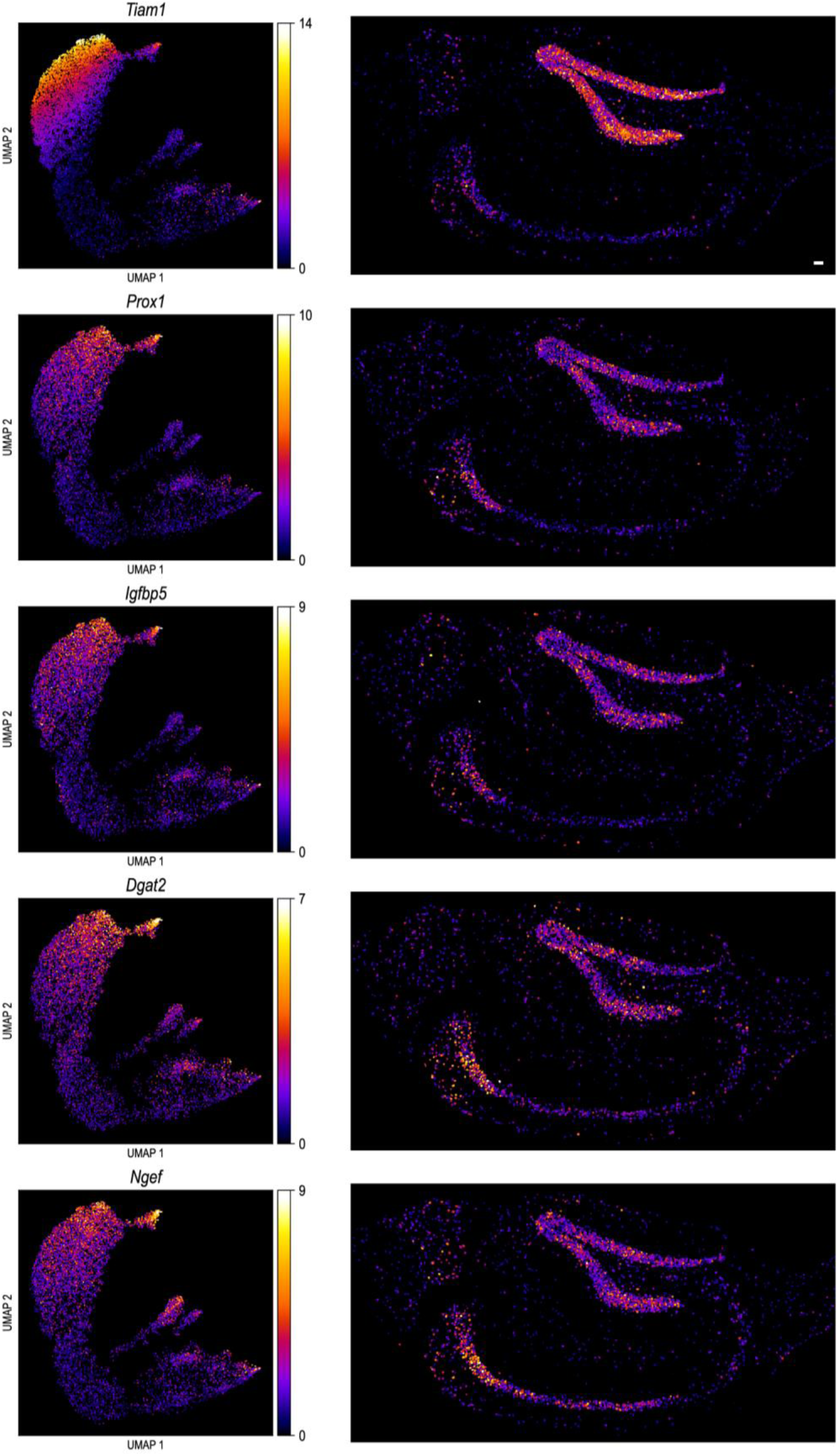
UMAP imputation of genes for hippocampal cluster 4. Gene expression levels for five selected genes are color-coded in the upper panel, providing a visual representation of gene expression variations among clusters. The left panel offers selected xy views of the hippocampal slice, illustrating the spatial distribution of gene expression. A unified color map adjacent to the UMAP panel aids in interpreting the gene expression levels. Scale bar: 50µm.

**Fig. S45.**
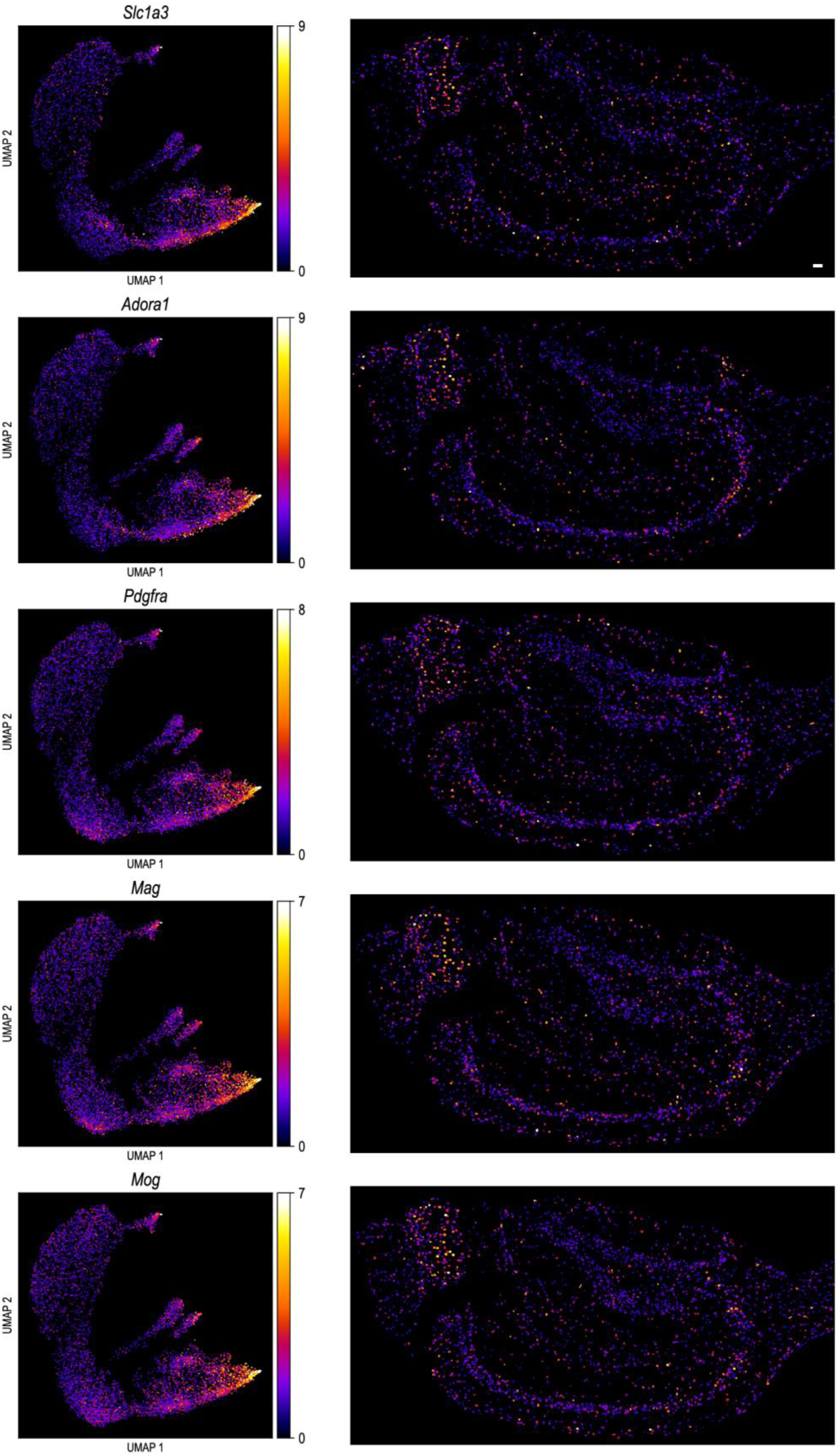
UMAP imputation of genes for hippocampal cluster 5. Gene expression levels for five selected genes are color-coded in the upper panel, providing a visual representation of gene expression variations among clusters. The left panel offers selected xy views of the hippocampal slice, illustrating the spatial distribution of gene expression. A unified color map adjacent to the UMAP panel aids in interpreting the gene expression levels. Scale bar: 50µm.

**Fig. S46.**
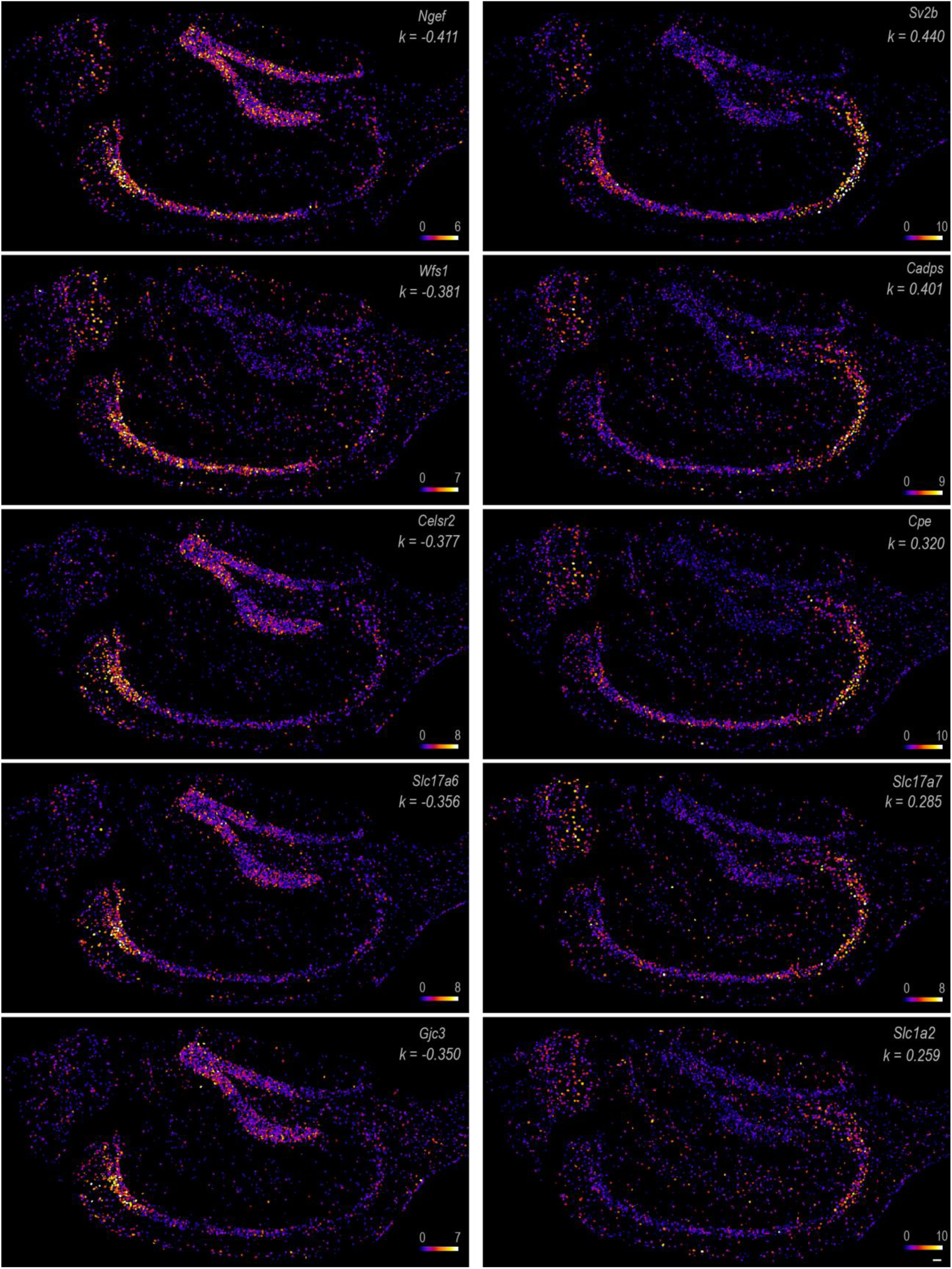
Gene expression gradients along the C1-C2-C3 axis. On the left are gene expression maps for 5 genes with the steepest decreasing slopes along the C1-2-3 segments showed in Fig. 5F, while on the right are 5 genes with the steepest increasing slopes. Cells are color-coded according to spot count, as indicated by the color bar. Scale bars: 50µm.

**Fig. S47.**
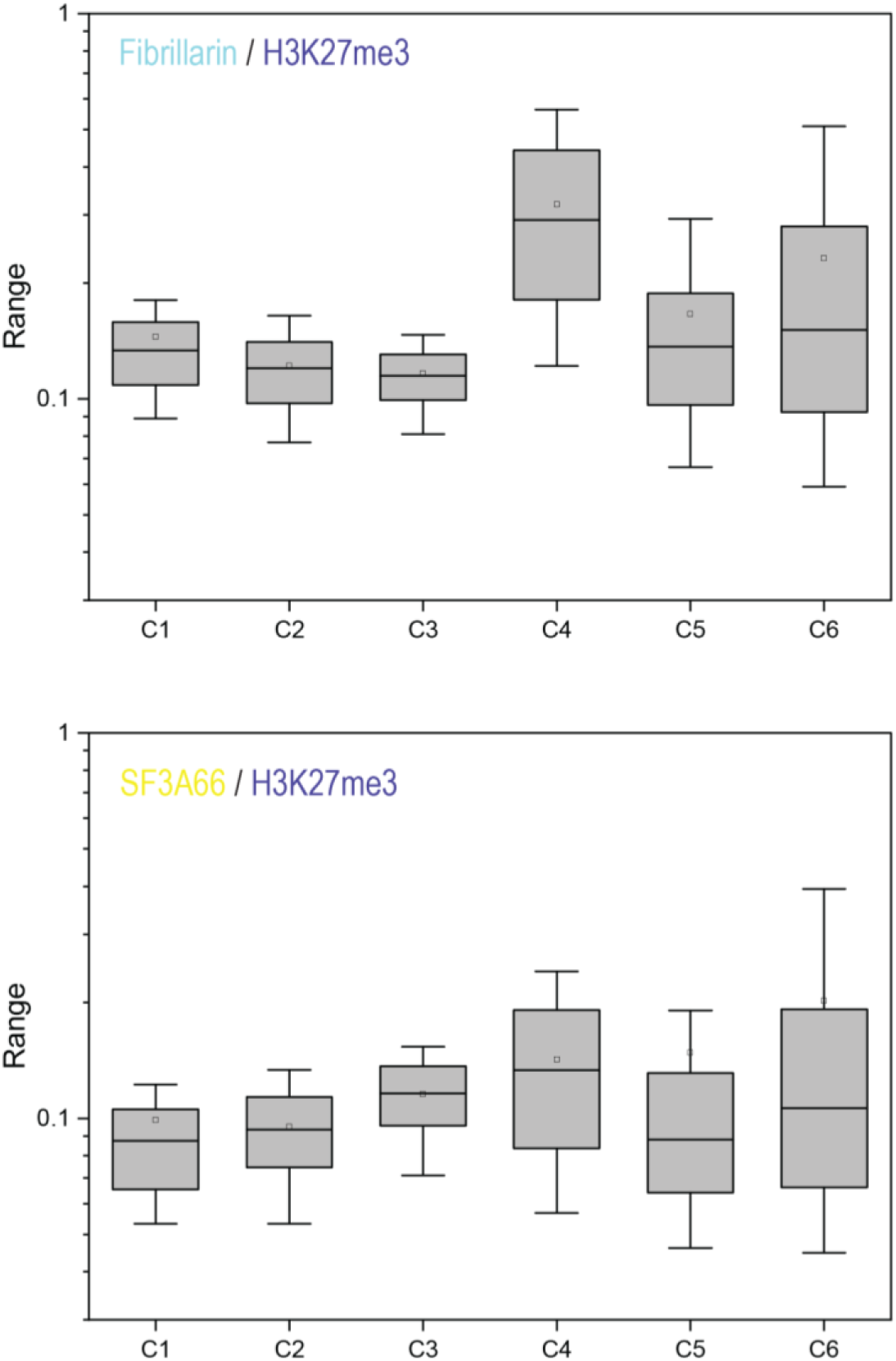
Nuclear structural variances among UMAP clusters in hippocampus. Boxplots depicting nuclear antigen levels (Fibrillarin – nucleolar fibrillar region; SF3A66 – nuclear speckles) normalized to inactive chromatin (H3K27me3) intensity for all clusters identified in the UMAP analysis of hippocampal cells. The lower and upper whiskers represent the 10th and 90th percentile values, respectively; the box represents the interquartile range (IQR) from the 25th to 75th percentile; the center line denotes the median; and the dotted line indicates the mean. The number of cells for each cluster is annotated in Fig. 5E.

## Movie Legends 1 to 6

**Movie 1**, 3D representation of RNA transcripts from three genes imaged using a single-round, three-color cycleHCR method in ∼200 µm thick cerebellar brain slice. The genes imaged are *Slc1a3* (L5+R5 at 488nm; green) marking Bergmann glia cells, *Stx1b* (L7+R7 at 561nm; yellow) for the granule layer, and *Rgs8* (L6+R6 at 640nm; magenta) for Purkinje cells. The movie is rendered using the 3D view in Imaris.

**Movie 2**, A detailed slice-by-slice examination of transcripts from ten genes within E6.5∼E7.0 mouse embryos, captured using cycleHCR technology over an axial depth of approximately 310 µm. Each gene is represented by a specific color, as detailed in Fig. 2A. The movie is rendered by using the ortho slice view in Imaris. Scale bar: 30 µm

**Movie 3**, 3D reconstruction of nine UMAP clusters within a mouse embryo, where cells from each cluster are colored according to the palette used in Fig. 3A. The movie is rendered by using Python image viewer napari with the plugin napari-animation.

**Movie 4**, cycleHCR images for 10 protein targets as illustrated in Fig. 4B. The movie is rendered using Imaris software.

**Movie 5**, 3D segmented models for 10 protein targets showed in Fig. 4B. The movie is rendered using ORS Dragonfly software.

**Movie 6**, cycleHCR images for 8 protein targets as shown in Fig. 5A-D. The movie is rendered using Imaris software.

## Supplementary Table Legends S1 to S4

**Table S1**, Barcodes and probe sequences used in cycleHCR imaging of 254 genes in the E6.5∼E7.5 mouse embryo

**Table S2**, Cell positions, transcript count per cell for 254 genes and UMAP assignment in the E6.5∼E7.5 mouse embryo

**Table S3**, Barcodes and probe sequences used in cycleHCR imaging of 120 genes in hippocampal slices.

**Table S4**, Cell positions, transcript count per cell for 120 genes and UMAP assignment in mouse hippocampal slice.

## Acknowledgments

We thank Monique Copeland and Benjamin Foster from Histology Shared Resources in Janelia Research Campus for preparing cerebellar and hippocampal slices. We also acknowledge Melanie Radcliff for her administrative support throughout this project. Special thanks to Andrew Woehler, Danial Smith, and Jeff Talbot at Advanced Instrumentation at Janelia Research Campus for their expertise in manufacturing custom-built components for our microscopy set-up. We appreciate the support from the HHMI Janelia Open Science Software Initiative (OSSI) for maintaining BigStitcher(-Spark), CellPose, and RS-FISH. This article is subject to HHMI’s Open Access to Publications policy. HHMI lab heads have previously granted a nonexclusive CC BY 4.0 license to the public and a sublicensable license to HHMI in their research articles. Pursuant to those licenses, the author-accepted manuscript of this article can be made freely available under a CC BY 4.0 license immediately upon publication.

## Author contributions

Z.J.L. conceived and designed the study, conducted robotic arm, fluidics, and microscope assembly and programming, and performed probe design and assembly. V.G. conducted cycleHCR RNA and protein imaging experiments with mouse embryonic stem cells, cerebellar, and hippocampal slices. J.K. and Z.J.L. performed image processing and single-cell analysis, including image stitching, registration, spot detection, 3D cell segmentation, spot-to-cell assignment, cell mask-based intensity calculation, UMAP analysis, and gene-to-cluster assignment. L.Y. and A.A.C. prepared the E6.5-7.0 mouse embryo for cycleHCR RNA imaging. V.G. and Y.L. validated antibodies for cycleHCR protein imaging. Y.L., A.H., and P.T. prepared the expanded mouse embryonic fibroblast for cycleHCR protein imaging. C.S. wrote custom codes for Cellpose-based 3D segmentation incorporating both *xy* and *yz* models and compatible with large datasets. T.K., K.R., G.F., and S.P. developed the Nextflow-orchestrated image processing workflow. Z.J.L. wrote the manuscript with inputs from other authors.

## Methods

### Cell lines

mouse ESCs were maintained in Knockout DMEM (GIBCO 10829-018) supplemented with 10% fetal bovine serum ES cell qualified (ATCC® SCRR-30-2020™), 1X GlutaMAX™ Supplement 100X (GIBCO, 35050-061), 1X MEM non-essential amino acids solution, 100X (GIBCO, 11140050), 1X Antibiotic-Antimycotic (GIBCO, #15240062), 0.1 mM 2-mercaptoethanol (GIBCO, 21985-023), 1000 U/mL LIF (EMD Millipore, ESG1106), 1μM PD03259010 (Millipore Sigma, PZ0162), 3μM CHIR99021 (StemCell Technologies, 72052). Primary mouse embryonic fibroblasts (pMEFs) were maintained in low glucose DMEM (GIBCO, 12320032) supplemented with 10% fetal bovine serum, 1X GlutaMAX™ Supplement and 1X Antibiotic-Antimycotic. Cell lines were maintained at 5% CO2 at 37 °C.

### Mice and tissue sections

C57BL/6J mice (3 months old) were used in our study. Primary rodent work was performed in accordance with protocols approved by the Janelia Research Campus Institutional Animal Care and Use Committee (IACUC) guidelines. Mice were housed in a 12 h light/dark cycle. Animals were anesthetized with isoflurane and perfused with RNase-free PBS (15ml) followed by 50 mL 4% paraformaldehyde (PFA) buffered to 0.1M Phosphate Buffer (pH 7.4). After dissection, brains were fixed in 4% PFA overnight, rinsed into 1x PBS, immersed in 30% (w/v) RNase-free Sucrose (Sigma-Aldrich, S7903) in 1x PBS to shake overnight, until the brain sank. Cerebellum was dissected out and embedded in OCT (Fisher Scientific, 23-730-571). 200 mm cryosections of the cerebellum were cut around Bregma -5.80mm. Cryosections were directly mounted onto PDL-coated silanized 40mm coverslips.

For flash frozen tissue, animals were anesthetized with isoflurane, checked for toe-pinch pain response, and decapitated before the hippocampus was dissected out. The hippocampus was oriented in OCT within a Peel-A-Way™ embedding molds (Millipore Sigma, E6032), and snap frozen by placing the mold into a dry ice/ethanol mixture (an approximate ratio of 5:1) until the OCT had frozen solid (about 5 min). OCT blocks were acclimated to -14°C in Leica CM 3050S Cryostat for 1 hour before 40 mm cryosections were collected coronally around Bregma -1.46mm. Cryosections were directly mounted onto PDL-coated silanized 40mm coverslips.

Mouse embryos were obtained from non-hormone primed Hsd:ICR(CD-1) females (Envigo) mated in house with B6D2F1 (C57BL/6xDBA) males. Insemination was verified the next morning by the presence of a copulatory plug, and this day was defined as E0.5 days post fertilization (d.p.f). Timed pregnant females were euthanized by cervical dislocation, and the embryos were recovered in 4% PFA (Electron microscopy sciences) diluted in RNase-free PBS at 4°C. Embryos were removed from the uterus and dissected from the decidua and parietal yolk sac using fine forceps, leaving the ectoplacental cone attached to the egg cylinder, as described previously (*35*). Subsequently, the embryos were fixed in fresh 4% PFA (RNase-free) overnight at 4°C, washed twice for 5 minutes with RNase-free PBS-T (1% PBS; Tween 20) on ice, and dehydrated in a series of methanol/PBS-T dilutions (10 minutes each) on ice as follows: 1) 25% MeOH/75% PBS-T; 2) 50% MeOH/50% PBS-T; 3) 75% MeOH/25% PBS-T; 4) 100% MeOH. Lastly, embryos were incubated at -20°C overnight (>16 hours).

### Coverslip cleaning and coating

40mm coverslips (Bioptechs, 40-1313-0319) were cleaned as described previously (*36*) with the following modifications. For the silanized coating, coverslips were immersed in a solution of 0.2% Triethylamine (Sigma Aldrich, 471283) and 0.3% allyltrichlorosilane (Sigma Aldrich, 107778) in Chloroform for 30 minutes. For tissue adhesion, silanized coverslips were coated with 100 mg/ml poly-D-lysine (Sigma P7280) dissolved in 1XPBS overnight. For cell adhesion, silanized coverslips were coated with 1 mg/ml Laminin (Gibco, A29248) for 2 hours at 37°C.

### cycleHCR probe design and assembly

#### cycleHCR primary probe selection

For precise single-shot RNA imaging using cycleHCR, a comprehensive multi-step probe selection process is applied to each target transcript, ensuring high specificity and robust signal amplification. This process encompasses: 1) A sliding window approach is employed to identify an optimal 92bp sequence within the target transcript for probe design. This sequence is divided into two segments of 45bp each for the left and right probe pairs, with a 2bp gap between them. The selection criteria for these probe halves include: a maximum secondary structure melting temperature (Tm) of less than 76°C; a maximum cross-sequences with six consecutive identical nucleotides (GGGGGG, CCCCCC, TTTTTT, AAAAAA) and a minimal gap of 2bp between any two consecutive probes along the 92bp region. 2) The Tm for DNA:RNA duplexes is typically 10–15°C higher than that for DNA:DNA interactions (*37*). Thus, Tm(probe:RNA) is conservatively estimated using the formula: **Tm(probe:RNA) = Tm(probe:DNA) + 10°C**, where **Tm(probe:DNA)** is calculated using the nearest-neighbor method (*38*) with a salt concentration of 1 mol. Additionally, RNA:DNA duplexes are less sensitive to formamide denaturation, particularly at high formamide concentrations (*39*). These selection criteria ensure probe-target interaction stability under stringent stripping conditions. 3) The final step in probe selection involves a thorough specificity check against the entire genome. This involves screening for 26bp junction sequences, representing the unique overlapping region between left and right probe halves. Any probe pair matching more than once in the genome is discarded to eliminate the risk of non-specific targeting.

### L and R barcoding probe sequence selection

For the selection and assembly of L + R barcoding probes, 14bp RNA and protein left and right readout probes was initially generated by using the randseq() function in Matlab 2022. The probe sequences underwent rigorous screening to adhere to specific criteria, aiming to enhance their specificity and minimize non-specific interactions. These criteria included: 1) Exclusion of sequences containing runs of four or more identical nucleotides (GGGG, CCCC, TTTT, AAAA) to prevent formation of secondary structures that could impair hybridization efficiency. 2) Elimination of sequences with the potential to form dimers or hairpin structures, which could interfere with probe-target interactions. 3) Ensuring a GC content between 30% and 60%, balancing hybridization strength and specificity. 4) A melting temperature constraint of less than 45°C to facilitate selective binding under the assay conditions. 5) Screening each new sequence against all existing sequences and their complementary sequences in the library to ensure a maximum cross-hybridization melting temperature (Tm) lower than 25°C, thereby reducing the likelihood of non-specific cross reaction. Through this method, a total of 230 probes were successfully generated, with 180 designated for barcoding cycleHCR RNA targets and the remaining 50 for barcoding cycleHCR protein targets.

#### L and R barcoding probe assembly

The assembly of L and R barcoding probes for cycleHCR involves a division and pairing process to enable specific and efficient Hybridization Chain Reaction (HCR) for each color channel. The process is outlined as follows: 1) The 180 RNA barcoding probes are divided into two sets of 30 Left (L) and 30 Right (R) probes for each color channel. This division ensures a wide range of unique barcode combinations, facilitating the multiplexing capability of the cycleHCR system. 2) To prepare the L and R probes for HCR initiation, we link a specific 18bp split HCR initiator sequence to the 5’ end of each left barcode, separated by a short spacer (AA). Similarly, the right barcode is linked to the 5’ end of the other 18bp split HCR initiator separated by a short spacer (AA), specific to the same color channel. This initiator sequence was designed to trigger the HCR reaction in a color channel specific manner as previously described (*11*). 2) Following the assembly, the L and R barcoding probes, now equipped with their respective HCR initiator sequences, are synthesized using Integrated DNA Technology.

#### Primary probe library assembly

For each RNA target, we chose between 10 to 25 probe pairs. The construction of these probe pairs involves the assembly of left and right primary probes as follows: 1) Left Primary Probe Assembly: A forward PCR sequence containing a T7 promoter (TAATACGACTCACTATAGCGTCATC) initiates the assembly. This is followed by the first 45bp sequence of the selected 92bp segment. A spacer sequence ’TT’ is inserted next. The 14bp left barcode sequence comes after the spacer. The assembly concludes with a reverse oligo sequence (CGACACCGAACGTGCGACAA). 2) Right Primary Probe Assembly: Similar to the left probe, it starts with the same forward PCR sequence with the T7 promoter. The 14bp right barcode sequence is placed immediately after the promoter. A spacer ’TT’ follows the barcode. The last 45bp of the 92bp primary probe sequence is then added. The assembly is completed with the same reverse oligo sequence (CGACACCGAACGTGCGACAA). Each assembled probe, whether left or right, ultimately spans a total length of 92bp. The primary probe sequences used for the 254 gene and 120 gene libraries for the mouse embryo and hippocampal experiments are in Table S1 and S3.

#### Protein cycleHCR probe design and synthesis

The 20 bp oYo docking sequences were generated using the randseq() function in Matlab 2022 and screened for cross-hybridization with a melting temperature (Tm) below 40°C. DNA sequences linked to oYo (5’ linked) were manufactured by AlphaThera, Inc. The gel anchoring sequence was constructed by concatenating the reverse complementary sequence of the oYo docking (18-20 bp), a 5 bp random sequence, the reverse complementary sequence of the right barcode, 2 bp of AA, and the reverse complementary sequence of the left barcode. The 5’ Acrydite modification (/5Acryd/) was added during oligo synthesis by Integrated DNA Technology (IDT).

### DNA synthesis and purification

ssDNA libraries were synthesized by Twist Bioscience. dsDNA was amplified using KAPA HiFi HotStart Polymerase (Roche, KK2502) using the following two primer sequences (T7 Forward: 5’-TAATACGACTCACTATAGCGTCATC-3’; Reverse: 5’-TTGTCGCACGTTCGGTGTCG-3’) and purified with DNA Clean and Concentrator-5 Kit (Zymo Research, 11-302). dsDNA was converted to RNA with HiScribe™ T7 High Yield RNA Synthesis Kit (NEB, E2040S) and purified using Monarch® RNA Cleanup Kit (NEB, T2040). An RNA/DNA mix was obtained by retrotranscribing 200pmol of RNA + 200pmol of the Reserve primer (5’-TTGTCGCACGTTCGGTGTCG-3’) with Maxima H Minus Reverse Transcriptase (Thermo Scientific, EP0753) as per manufacturer instructions. The RNA was digested with Thermolabile USER (Uracil-Specific Excision Reagent) II (NEB, M5508) overnight before alkaline hydrolysis in 1M NaOH at 65°C for 15 minutes. Hydrolysis was neutralized with 1M acetic acid before purifying the libraries using ssDNA/RNA Clean & Concentrator^TM^ (Zymo Research, 50-444-498).

### RNA cycleHCR in tissues and cells

#### Fixation and permeabilization

Fixed tissues were washed 3 times in 1X PBS (for 5 minutes each, then incubated in ice-cold methanol (Fisher Chemical., A454-1), on ice, for 20 minutes, 5 minutes in ice-cold [50% methanol, 50% 2X SSC -0.1% Tween] and 5 minutes in cold-[25% Methanol, 75% 2XSSC - 0.1% Tween]. 25% methanol was then removed by rinsing three times with 1X PBS and washing once in 1X PBS for 5 minutes (tissue ≤ 40um) or 10 minutes (tissue ≥ 100um). Tissue was permeabilized in [0.5% Triton, 1XPBS] for 30 minutes at room temperature, followed by 3 washes in 1X PBS for 5 minutes each.

Cells were fixed in 4% paraformaldehyde (PFA) in 1XPBS (Invitrogen, AM9625) for 10 minutes and permeabilized in 0.5% Triton for 15 minutes.

#### Gel embedding and Proteinase K digestion

After permeabilization, samples were incubated in 0.1mg/ml Acryloyl-X SE (AcX) for 1 hour at room temperature. AcX (Invitrogen, A20770) was prepared as described (*40*). Samples were then washed twice in 1XPBS, 10 minutes each time. Before proceeding with the gelation step, samples were incubated for 5 minutes (tissue ≤ 40μm) or 10 minutes (tissue ≥ 100μm) in polyacrylamide (PA) solution [4% acrylamide/bis acrylamide (BioRad., 1610154), 60mM Tris HCl pH 8 (Corning, 460131), 0.3M NaCl (Corning, 46032]. For gelation, APS (Sigma, A3678) and TEMED (Sigma, T7024) were added to the PA solution with a final concentration of 0.03% and 0.15% respectively. Gelation chamber was assembled as described (*36*). Briefly, a pre-cleaned slide 75 x 50 mm, thickness 0.96 to 1.6 mm (Corning 2947-75X50) was coated with Gel Slick (Lonza, 182369) for 15 minutes and let air dry for at least 30 minutes. 100μL of PA solution with APS and TEMED were added to the dry-glass plate and sample was slowly overlaid to avoid air bubbles. After 1.5 hour at room temperature, the gel fully polymerized and sample was gently detached from the glass plate. The silanized coating allows covalent binding of the hydrogel to the 40mm coverslips, while the gel slick coating prevents the gel from sticking to the glass plate during the polymerization. As a result, when the 40mm coverslip is gently detached from the glass plate, the gel-embedded sample will remain stably bound to the coverslips during further processing.

Tissue samples were then digested with Proteinase K (NEB, P8107S) as described (*41*) with some modifications. Tissue was incubated in [2X SSC, 2% SDS, 0.5% Triton, 1:100 dilution of Proteinase K (800units/ml; NEB P8107S)] for 16 hours in a humid chamber at 37°C.

Cells were digested as described (*36*).

#### RNA quality control and hybridization with ssDNA libraries

Steps that required small volumes (100-200mL) were performed in custom-made hybridization chambers of the following dimensions: [0.12mm chamber depth /20mm diameter] for samples ≤ 40µm; [0.5 mm chamber depth /19 mm diameter] for tissue 100-200 µm; [1mm chamber depth/ 10mm diameter] for embryos. To assemble the hybridization chambers, a 13-in-1 heavy duty hollow punch sets was used to cut the internal diameters indicated above from a round-silicon sheet (BIOPTECHS, 1907-1422-500 and 1907-1422-500). The prepared gasket was then securely affixed to a pre-cleaned slide 75 x 50 mm, thickness 0.96 to 1.6 mm (Corning 2947-75X50) using 100% silicone sealant (GORILLA). For the 0.12mm x 20mm chamber, am imaging spacers (Grace Bio-Labs Cat.# 654006) was attached to the pre-cleaned slide as described above.

Before incubating the sample with primary probes, RNA quality was assessed using our Alexa Fluor 488-conjugated-readout probe (RO22: 5’-GCCAAGATGGAGTTA-3’) targeting ribosomal RNAs. After proteinase K digestion, samples were washed twice in 5XSSC-T [5X SSC, 0.1% Tween] for 15 minutes. Ribosome probes were diluted in 10% EC Buffer [10% ethylene carbonate (Sigma, E26258), 10% dextran sulfate (Sigma, D4911), 2X SSC) at the following concentrations: 100nM RO22 probes (samples ≤ 40µm) or 200nM (tissue ≥ 100μm). Samples were incubated in the appropriate concentration of RO22 for 1 hour at room temperature and washed once (samples ≤ 40µm) or twice (tissue ≥ 100μm) in 10% formamide solution [10% deionized formamide (Ambion, AM9342), 0.1% triton, 2X SSC) for 10 minutes. Before imaging, samples were then washed in 5X SSC-T before staining nuclei with 5mg/ml DAPI.

After assessing RNA quality, samples were incubated with primary probe libraries. Primary probes were added to a total volume of 100µL (for 0.12mm x 20mm or 1mm x 10mm diameter chambers) or 250µL (for 0.5 mm /19 mm chamber) hybridization solution [50% deionized formamide, 10% Dextran Sulfate, 2X SCC] in the following amount: ∼2µg of ssDNA libraries targeting 10 RNA species or less, ∼25 µg of ssDNA libraries targeting RNA transcripts from 120 genes, ∼50 µg of ssDNA libraries targeting RNA transcripts from 279 genes. Samples were hybridized with primary probes at 37°C for about 20 hours in RapidFISH Slide Hybridizer Oven (Boekel Scientific, Cat. #13-245-230). High stringent wash was carried out in 80% formamide solution [80% formamide, 4X SSC, 0.1% Triton) at 32°C for 20 minutes before readout probe hybridization.

### Readout probe hybridization and hybridization chain reaction (HCR)

Samples were incubated with 200nM readout probes in 10% EC Buffer for an hour at room temperature, rinsed three times in 5X SSC-T and subsequently washed once (samples ≤ 40µm) or twice (tissue ≥ 100μm) in 10% formamide solution [10% deionized formamide, 0.1% triton, 2X SSC) for 10 minutes. Samples were then washed three times in 5X SSC-T before HCR. Expanded samples were incubated with 200nM readout probes for 3 hours and washes performed as described above.

For HCR, H1 and H2 amplifiers (HCR^TM^ Amplifiers: B4 fluorophore 488, B3 fluorophore 647, B2 fluorophore 560, Molecular Instruments, Inc.) were activated according to the manufacturer’s instructions. After activation, H1 and H2 were added to the Amplification buffer [10% dextran sulfate, 0.1% Tween, 5XSSC) in a 1:100 ratio. In non-expanded samples, HCR was carried out for 1.5 hour at 32°C and washed twice in 5X SSC-T for 10 minutes before imaging. In expanded samples, HCR was carried out for 3 hours at 32°C.

### Stripping and reprobing

Readout probes and HCR chains were removed by incubating the sample in 80% formamide solution (80% deionized formamide, 0.1% triton, 4XSSC) for 20 minutes at 32°C. Samples were then rinsed three times in 5X SSC-T and washed once in 5X SSC-T for 10 minutes before reprobing with readout probes for the consecutive round.

### Sample preparation for cycleHCR protein imaging

#### Fixation and permeabilization

40μm cryosections were removed from -80°C and immediately fixed. Tissues and cells were both fixed in 4% PFA for 10 minutes and permeabilized in 0.5% Triton in 1XPBS for 15 minutes at room temperature. After permeabilization, samples were incubated for 1 hour in blocking solution [0.25% Triton, 0.5mg/ml salmon sperm DNA (Fisher Scientific AM9680), 10mg/ml nucleases- and proteinases-free BSA (Sigma, 126609), 1X PBS]. During blocking, the antibodies conjugated with a unique docking oligo were assembled with the acrydite^TM^ 5’-gel docking oligos (IDT).

#### In vitro assembly of antibodies-oYo-oligos with acrydite^TM^ 5’-gel docking oligos

oYo Link® conjugated with each unique docking oligos were purchased from AlphaThera and crosslinked to the antibodies of interest according to the manufacturer’s instructions. For in vitro assembly of gel docking complex, a mix of 1 μl oYo-crosslinked antibodies, 7 μM acrydite^TM^ 5’-gel docking oligos and 10% dextran sulfate in 1X PBS was incubated for 1 hour at room temperature on an orbital shaker at 100rpm.

#### Antibody staining

Pre-assembled gel docking complexes were combined in 100μl blocking solution and incubated with the sample overnight in a humid chamber at 4°C. To preserve RNA during incubation with primary antibodies, 6 units of SUPERase•In™ RNase Inhibitor (Invitrogen, AM2694) were added to the antibody mix. The next day, samples were washed three times in 1X PBS, before post-fixation in 4% PFA for 10 minutes. Samples were washed in 1X PBS and then incubated with 0.1mg/ml Acryloyl-X SE (AcX) for 1 hour at room temperature. After Acx modification, non-expanded samples were embedded in a thin-hydrogel layer, digested with proteinase K and incubate with readout probes as described in RNA cycleHCR section. For simultaneous detection of protein and RNA, samples were incubated with primary probes after proteinase K digestion as described in RNA cycleHCR section.

### Protein cycleHCR with expansion

Protein cycleHCR was combined with expansion microscopy using a protocol modified from a previous report (*42*). Samples were prepared as described in Protein cycleHCR. After overnight incubation with the mixture containing the pre-assembled gel docking complexes, samples were treated with 200 μg/mL acryloyl-X SE in 1XPBS for 1 hour at room temperature and rinsed twice with 1XPBS for 15 minutes each. Samples were then embedded in TREx1000 gelation solution [TREx1000: 1 M sodium acrylate (Sigma, 408220), 14% acrylamide (Bio-Rad, 1610140), 1000 ppm N,N’-methylenebisacrylamide (bis, Sigma, M7279), 1XPBS, 2000 ppm APS (Sigma, A3678), 2000 ppm TEMED (Sigma, T7024), and 100 ppm 4-hydroxy TEMPO (4HT, Sigma, 176141)] on a chamber composed of a Gel Slick-treated glass slide and 1-layer Scotch Magic Tape spacers (∼56 µm, 3M, #810) (see details in (*42*)). Gel was polymerized for 3 hours at room temperature to prevent denaturing the gel docking strand from the antibody docking strand. The cell-embedding gel was then detached from the coverslip, cut into smaller pieces, and digested in proteinase K (NEB, P8107S) diluted 1:100 in proteinase K digestion buffer (50 mM Tris-HCl pH 8, 500 mM NaCl, 1 mM EDTA, 0.5% Triton X-100, and 1% SDS) overnight at 37 °C. The gel was washed 3 times in 1XPBS and three times in nuclease-free water for 30 minutes each time to fully expand. The expanded gel was then incubated in re-embedding gelation solution for 30 minutes on ice. The re-embedding gelation solution contains 44% acrylamide, 0.1% N,N’-methylenebisacrylamide (bis), 0.2% TEMED, 0.01% 4HT, and 0.2% APS. The re-embedding gel was then allowed to polymerize on a cleaned, silanized, and poly-D-lysine coated 40 mm coverslip in a chamber with 4-layer-Scotch Magic Tape spacers for 1 hour at 37°C. After re-embedding, the samples were loaded on our automated microfluidics imaging system to acquire protein cycleHCR data.

### Antibodies

The following rabbit antibodies were used at 1:100 dilution unless otherwise indicated. Anti-Fibrillarin (Abcam, ab5821), Anti-Histone H3 (mono methyl K4) (Abcam, ab8895), Anti-Signaling, 9733), anti-GFAP (Abcam, ab278054) used at 1:200, anti-Iba1 (Abcam, ab178846), anti-Nup98 (Cell Signaling, 2598) used at 1:30 dilution, anti-Tomm20 (Abcam, ab186735), anti-Calnexin (Abcam, ab22595), anti-RPS6 (Abcam, ab225676), anti-RAB7 (Abcam, ab126712), anti-EEA1 (Abcam, ab2900), anti-LAMP2A (Abcam, ab18528), anti-LC3B (Abcam, ab192890), anti-Na/K ATPase (Abcam, ab76020), anti-Fibrillin (Abcam, ab53076), anti-CDK9 (Abcam, ab239364).

The following mouse antibodies were used at 1:100 dilution: anti-SF3a66 (Abcam, ab77800), anti-SC35 (Abcam, ab11826), anti-Ankyrin-G (Antibodies Inc., 75-146). anti-Syntaxin 6 (BD Biosciences, 610636), anti-Lamin B1 (Abcam, ab8982).

### cycleHCR fluidics and imaging system

The fluidics system is designed for precision in flow control and mixing, essential for cycleHCR imaging processes. It incorporates an OB1 flow controller for managing the flow rate through the system, two MUX distribution valves for directing flow to specific channels and a BFS Coriolis flow sensor to provide feedback for accurate flow control. This setup accommodates 10 probe tubes for L+R readout probes, 4 solution types including buffers and washing solutions and 2 hairpin mix tubes - One for H1 hairpin mix and another for H2 hairpin mix. The real-time mixing of H1 and H2 hairpin mixes is regulated by alternating the flow between them using a 2 to 1 MUX valve, ensuring efficient HCR reaction initiation. Flow Rate: Optimized at 150 µl/min to balance efficiency and sample integrity. After primary probe hybridization, samples on a 40mm coverslip are placed in a closed-top FCS2 chamber for imaging. This chamber is then positioned on a customized stage adapter on the microscope. All buffers used in the following fluidics steps were filtered through 0.22mm vacuum filter (Corning, 431098).

#### Fluidics Steps

1. Stripping Buffer (80% formamide, 4X SSC, 0.1% Triton) Incubation (20 mins): Removes bound probes and prepares the sample for the next cycle.
2. Washing (20 mins): Uses 5X SSC and 0.1% Tween buffer with 5μg/ml DAPI to clean the sample.
3. Readout Probe Incubation (30 mins): Applies mixed Left and Right probes for a new cycle of target visualization. 200nM of each readout probes were added to 1.8mL of hybridization solution (10% Ethylene Carbonate, 5X SSC, 10% Dextran Sulfate)
4. Washing (20 mins): Another round with 5X SSC with DAPI.
5. Hairpin Mix Injection (5 mins): Introduces H1 and H2 mixes for signal amplification. H1 and H2 were added to the amplification buffer (5X SSC, 0.1% Tween, 10% Dextran Sulfate) at 1:50 ratio.
6. HCR Amplification (90 mins): Enables the fluorescent signal to build.
7. Washing (10 mins) as in step 4: Cleans excess reagents post-amplification.
8. Imaging Buffer (50mM Tris HCl pH8, 2mM Trolox (Sigma, 238813), 1mg/ml Glucose Oxidase (Sigma, G2133), 1:100 Catalase (Sigma, C3155), 0.8% D-Glucose, 5X SSC) Injection (20 mins): Prepares the sample with an oxygen scavenger to prevent photo-crosslinking and enhance signal strength.
9. TTL Signal Activation: Triggers the microscope for imaging after a final 10-minute incubation.

#### Microscopy

The Nikon CSU-W1 spinning disk microscope, equipped with advanced features, is utilized for high-resolution imaging in cycleHCR technology. This setup includes:

1) A 25X CFI PLAN Apochromat Lambda S silicone oil immersion objective with a numerical aperture (N.A.) of 1.05 and a working distance of 0.55 mm, ideal for imaging intact mouse embryos and protein cycleHCR imaging in expanded primary mouse fibroblasts.
2) A 40X CFI PLAN Apochromat Lambda S silicone oil immersion objective with an N.A. of 1.25 and a working distance of 0.3 mm, suited for imaging brain tissue slices.
3) An uniformizer: Ensures even illumination across the field of view.
4) 6 Laser Lines (405nm, 514nm, 561nm, 594nm, 640nm): Provide a range of excitation wavelengths for versatile fluorophore excitation.
5) A Hamamatsu BT Fusion Camera: Captures high-quality images with efficient signal detection.

For the 25X objective, the microscope operated in ultraquiet mode with a fixed framerate of 5.1Hz, optimizing conditions for sensitive samples like intact mouse embryos and expanded primary mouse fibroblasts. For the 40X objective, imaging of brain tissue slices was performed with a 100ms exposure in the standard camera readout mode. The Nikon’s Perfect Focusing System (PFS) maintained the *Z* position of the objective between imaging rounds, critical for long-term imaging experiments. TTL signals automated the transition between imaging rounds and fluidic cycles, streamlining the cycleHCR process. Temperature control during imaging was ensured by heating the objective with a Tokai Hit Lens Heater controlled by a TPi controller (TPiE-LH), while the imaging chamber temperature was regulated using a Bioptechs FCS controller.

### Image stitching

Large images composed of multiple 3D tiles were aligned using BigStitcher (*43*) and the corresponding BigStitcher-Spark framework for distributed execution (https://github.com/JaneliaSciComp/BigStitcher-Spark). After re-saving image data into the multi-resolution N5 format, alignment was performed independently for each imaging round, initialized using the 20% overlap (in xy) between neighboring tiles. The calculation of the final 3D affine transformation for each tile consists of several steps: pairwise shift calculation using phase correlation, pre-viewing and filtering pairwise shifts, global optimization of pairwise shifts, and affine refinement of the alignment using the Iterative Closest Point (ICP) algorithm (*44*). The pairwise shifts were calculated using 8x8x4 (xyz) downsampling and averaging intensity from all channels, ICP refinement was also computed using 8x8x4 downsampling. The tiles were fused either by weighted average fusion or the one-tile-wins strategy. The weighted average fusion strategy computes the values of pixels in the overlapping regions using a distance-weighted (from the tile boundaries) average, which was used for visualizing protein labeling. The one-tile-wins fusion strategy does not perform averaging in overlapping areas, but instead copies (and interpolates) pixel values from the input tile that was imaged first (out of all overlapping tiles at any given pixel). The one-tile-wins strategy was used for calling single-molecule spots from RNA labeling. This strategy solves decreased spot-detection frequency in the overlapping regions due to averaging pixel values when the same single-molecule spot is slightly misaligned in the different overlapping tiles. The Nextflow pipeline for image processing and stitching is available at https://github.com/liulabspatial/cycleHCR.

### Image registration

To adjust for shifts in the sample or field of view during multiple cycles of imaging, the images of different rounds were registered to the image from a reference round using the Python package bigstream (version 1.2.9) (*13*). The reference round was chosen based on manual inspection, and all other rounds were transformed such that the DAPI channels across different rounds align in 3D space.

For single-tile images, the global affine transformation was sufficient. The affine transformation matrix was obtained using the alignment_pipeline function with a matrix was then used in the apply_transform function to perform uniform translation, scale, sheer, and rotation on all pixels.

For stitched multi-tile images, the global affine and local deform transformations were sequentially performed. The affine matrix was calculated as above, and then the deform matrix was calculated using the distributed_piecewise_alignment_pipeline function initialized with the affine matrix. The two matrices were provided as sequential steps, affine and then deform, in the distributed_apply_transform function to first perform the global affine transformation to the whole image, followed by local deformable transformation uniquely defined for each pixel. The parameters and codes for bigstream are available at https://github.com/liulabspatial/cycleHCR.

### Cell segmentation

For 3D nucleus segmentation of the mouse embryo, a unified custom model was trained using the human-in-the-loop feature in Cellpose 2 (*17*) on *xy* and *yz* slices in which nuclei were manually labeled. Subsequently, 3D segmentation function (*45*) was utilized to perform the segmentation across all dimensions. Masks less than 1000 voxels were filtered out.

In contrast, for 3D nucleus segmentation of the hippocampal slice, two distinct custom models were trained separately for the *xy* and *yz* orthogonal views, again utilizing the human-in-the-loop feature in Cellpose 2. A custom 3D segmentation procedure was then implemented by computing the xy flows using the *xy* model, and computing the *yz* and *xz* flows using the *yz* model. Next, the consensus cell flow was calculated by averaging across *xy*, *yz* and *xz* flows and then the dynamics were run on the flows to compute the masks. Masks less than 1000 voxels were filtered out.

The segmentation accuracy was estimated by human inspection of raw DAPI images and corresponding masks. Further manual curation and size filtering to eliminate potential oversized doublets were performed using ORS Dragonfly software.

The parameters and codes for Cellpose segmentation are available at https://github.com/liulabspatial/cycleHCR/.

### Spot detection and spot-to-cell assignment

The spot detection process involved three-dimensional single molecule localization using RS-FISH (*16*). Specific parameters for each experiment were initially selected using the interactive feature provided by RS-FISH, as detailed in fig. S17 and fig. S39. These parameters were carefully chosen to minimize false positive detections. Importantly, the same parameters were consistently maintained across all images within each series.

Spot-to-cell assignment was achieved by rounding the x, y, z coordinates of each spot. Subsequently, cell assignment was determined by matching the rounded x, y, z voxel value of the labeled cell mask image.

The parameters and codes for RS-FISH localization and spot-to-cell assignment are available at https://github.com/liulabspatial/cycleHCR/.

### Mask-based image quantification

We utilized the *regionprops3()* function in Matlab 2023b, to compute center-of-mass and intensity measurements on labeled and grayscale image volumes.

### UMAP analysis

The cell-by-gene matrix was filtered based on the total number of spots per gene and the total number of spots per cell. The total counts per gene were plotted as a histogram on a logarithmic scale. For the embryo data, Otsu’s method of thresholding was used to remove genes with low total counts using the Python package scikit-image (version 0.23.1) (*46*). Cells with less than 40 total counts were removed, resulting in the removal of approximately 5% of cells. For hippocampus data, Li’s method of threshold was used to remove genes with low total counts. Cells with less than 10 total counts across the retained genes were removed, resulting in the removal of approximately 7.5% of cells.

The filtered counts matrix was used as input for generating UMAP using the Python package Uniform Manifold Approximation and Projection (UMAP) (version 0.5.5) (*18*). The cells were clustered in an unsupervised manner using Python package HDBSCAN (version 0.8.33) (*47*). The parameters and codes used for UMAP and HDBSCAN are available at https://github.com/liulabspatial/cycleHCR.

### Gene-to-cluster assignment

We assigned each gene to one of the identified clusters based on average counts per cell in each cluster. The filtered counts matrix is normalized across each cell by dividing each element of the gene vector by the total counts per cell, thereby normalizing for the potential overrepresentation of certain cells in gene vectors. For each gene, average counts per cell for each cluster are calculated. The gene is assigned to the cluster with the highest average counts. The unnormalized raw counts per cell were plotted in both UMAP and anatomical space to validate gene-to-cluster assignment.

